# The exhaustive genomic scan approach, with an application to rare-variant association analysis

**DOI:** 10.1101/571752

**Authors:** George Kanoungi, Michael Nothnagel, Tim Becker, Dmitriy Drichel

**Author notes:** Correspondence address: Dmitriy Drichel University of Cologne Cologne Center for Genomics Department of Statistical Genetics and Bioinformatics Weyertal 115b, 50931 Cologne, Germany.

## Abstract

Region-based genome-wide scans are usually performed by use of a priori chosen analysis regions. Such an approach will likely miss the region comprising the strongest signal and, thus, may result in increased type II error rates and decreased power. Here, we propose a genomic exhaustive scan approach that analyzes all possible subsequences and does not rely on a prior definition of the analysis regions. As a prime instance, we present a computationally ultra-efficient implementation using the rare-variant collapsing test for phenotypic association, the genomic exhaustive collapsing scan (GECS). Our implementation allows for the identification of regions comprising the strongest signals in large, genome-wide rare-variant association studies while controlling the family-wise error rate via permutation. Application of GECS to two genomic data sets revealed several novel significantly associated regions for age-related macular degeneration and for schizophrenia. Our approach also offers a high potential for genome-wide scans for selection, methylation and other analyses.

## Introduction

Genomic scans assess genomic regions (usually subsequences) with respect to some statistical measure and, ideally, quantify its consistency with the null hypothesis. Prominent applications include the detection of allele frequency differences between cases and controls in genetic association studies (CHRISTOPHERSEN *et al.* 2017), the departure of the site-frequency spectrum (SFS) from the expectation under neutral evolution in selection analysis (NIELSEN *et* al. 2005) and of differential methylation patterns in epigenomics (JAFFE *et al.* 2012). Although statistical tests differ, the basic procedure remains similar across these applications by comprising (1) the prior definition of a set of contiguous analysis regions (bins) *B*_*ij*_, characterized by start positions *i* and end positions *j* (“binning”), sometimes defined by setting scanning parameter values (“sliding window”); (2) the calculation of a suitable summary or test statistic, T(*B*_*ij*_), for each bin; (3) the distributional assessment of the statistics in order to identify extreme values, frequently including the calculation of p-values, and often, but not always, followed by control of the family-wise error rate (FWER).

With long chromosomal sequences, it is not known in advance which subset of possible subsequences is most suitable for statistical summarization and testing, i.e. which regions will provide the highest power. Use of a priori fixed regions, including sliding-window approaches with fixed bins, will result in a highly likely increase in the type II error rate and, correspondingly, reduced power, because regions comprising the strongest signal(s) will almost certainly not be chosen prior to the analysis. A more probable scenario is that a region of interest will only partially coincide with the chosen analysis region. As a consequence, the signal will be diluted by inclusion of non-relevant variants, split across multiple analysis regions, or both. Fixed, pre-determined binning therefore represents a major limitation of current genomic scans. Moreover, due to unknown correlation structures between regions, the correction for multiple testing is often performed in a conservative way, e.g. by use of Bonferroni correction for the number of tested regions (CHEN *et al.* 2017).

Here, we focus on the application of the exhaustive scan approach to rare-variant (RV) association studies based on sequenced or genotyped data. RV analysis is motivated by the observation that, although genome-wide association studies (GWAS) have usually identified common risk alleles for a wide range of complex diseases (MANOLIO *et al.* 2009), most of these alleles cause at most moderate increases in risk and contribute little to the overall heritability of diseases individually, leaving large portions of human diseases’ heritability unexplained (FELDMAN and RAMACHANDRAN 2018; MANOLIO *et al.* 2009). This observation motivated studies to focus on the role of RVs, aiming to deliver functionally interpretable variants of moderate to large effect sizes and explaining additional disease risk variability. While highly penetrant RVs play an important role in Mendelian disease etiology (GIBSON 2012), their role in complex disease etiology is less clear. Population-based study designs using single-variant phenotypic association tests are usually plagued by low power (CORDELL AND CLAYTON 2002) due to small observed allele counts for RV. Region-based RV association analyses are based on the assumption that multiple RVs in physical proximity have similar effects on the phenotype. Under this assumption, multiple RVs in a genomic region can be aggregated and analyzed as a unit. In this context, the most common approach is to define fixed bins by either using the locations of known protein-coding genes as regions of analysis or by using a sliding-window approach with two fixed parameters, namely the window size and the step size. Either choice is fundamentally limited in scope, and will consider only a tiny fraction of possible subsequences.

In RV analysis, “rareness” itself is another parameter that is usually defined by a threshold of the minor allele frequency, MAF_T_ Alternatively, weighting schemes have been proposed that assign lower weights to variants with higher allele counts. This does not fully solve the problem of rareness thresholds, as the shape of the weighting function is usually chosen somewhat arbitrarily and without a stringent justification of its usefulness.

Noteworthy progress towards non-parametric RV analysis has been made in (PRICE *et al.* 2010), who proposed the Variable-Threshold (VT) approach, in which test statistics for all possible MAF_T_ are computed and the optimal MAF_T_ is adapted from the data. The method uses permutation testing to adjust for the large number of tested hypotheses within a bin; it is therefore computationally more intense. In (DRICHEL *et al.* 2014), the VT method was extended to the collapsing and the CMAT tests (ZAWISTOWSKI *et al.* 2010), whereas the method became computationally impractical for regression models.

However, even if the problem of the unknown “rareness” can be alleviated, the problem of the choice of analysis regions remains, which has been acknowledged before (FIER *et al.* 2012; FREEDMAN 2003; TIMPSON *et al.* 2018; WU *et al.* 2010). The present work can be regarded as the extension of the VT method to binning of analysis regions (“variable binning”, VB).

Here, we suggest to perform an exhaustive scan for phenotypic association using a simple RV test (collapsing method, COLL) as the test statistic (LI and LEAL 2008). COLL dichotomizes samples by their carrier status, i.e. whether the corresponding individual is carrying at least one rare allele in the analysis region. In a case-control study design, a 1-df χ^2^-test can be applied to the resulting 2×2 contingency table. Interestingly, despite its relatively simple disease model, the power of COLL is comparable to more sophisticated methods for a wide range of disease models (ZAWISTOWSKI *et al.* 2010). However, COLL is inherently limited in that it can only be applied to binary phenotypes only, does not account for covariates, and has limited power if the associated RVs in the region have different effect directions. A large number of more advanced tests have been developed, see (BASU and PAN 2011; DRICHEL *et* al. 2014; LEE *et al.* 2014) for categorizations. A notable example is the sequence kernel association test (SKAT) (WU *et al.* 2011), which is a variance-component test and sensitive to mixed effect directions in a region, allows for inclusion of covariates and can be used with binary and quantitative phenotypes.

Here, we propose the use of exhaustive scans to all possible contiguous subsequences and to perform multiple-testing correction by obtaining the distribution of extreme p-values from replicates of the data simulated under the null hypothesis by repeatedly permuting case-control status. We introduce this approach, in an exemplary way, for a specific application, namely the genomic exhaustive collapsing scan (GECS) approach for COLL, and present a computationally efficient implementation of GECS. We show that, although the number of possible contiguous bins for all RVs at a single chromosome is very large, namely *n(n+1)/2* with *n* variants, the number of *distinct* bins dramatically reduces by about 3 to 4 orders of magnitude, rendering GECS feasible and scalable even for whole-genome sequence data in large sample sets. Furthermore, this acceleration allows control of the family-wise error rate (FWER) via repeated case-control status permutation which provides optimal power to detect association (MEINSHAUSEN *et al.* 2011). Based on simulations, we derive empirical thresholds for genome-wide significance in case-control WGS studies for different sample sizes and minor allele frequency thresholds, in an approach similar to (PULIT *et al.* 2017). In applying GECS to two real-world data sets, we show that our approach is feasible and scalable with large, modern association studies and provides a fine-grained, base-pair resolution of associated regions contained in the data (Figure 1), which will enable a deeper understanding of the effect of RVs on the etiology of complex diseases.

**Figure 1:**
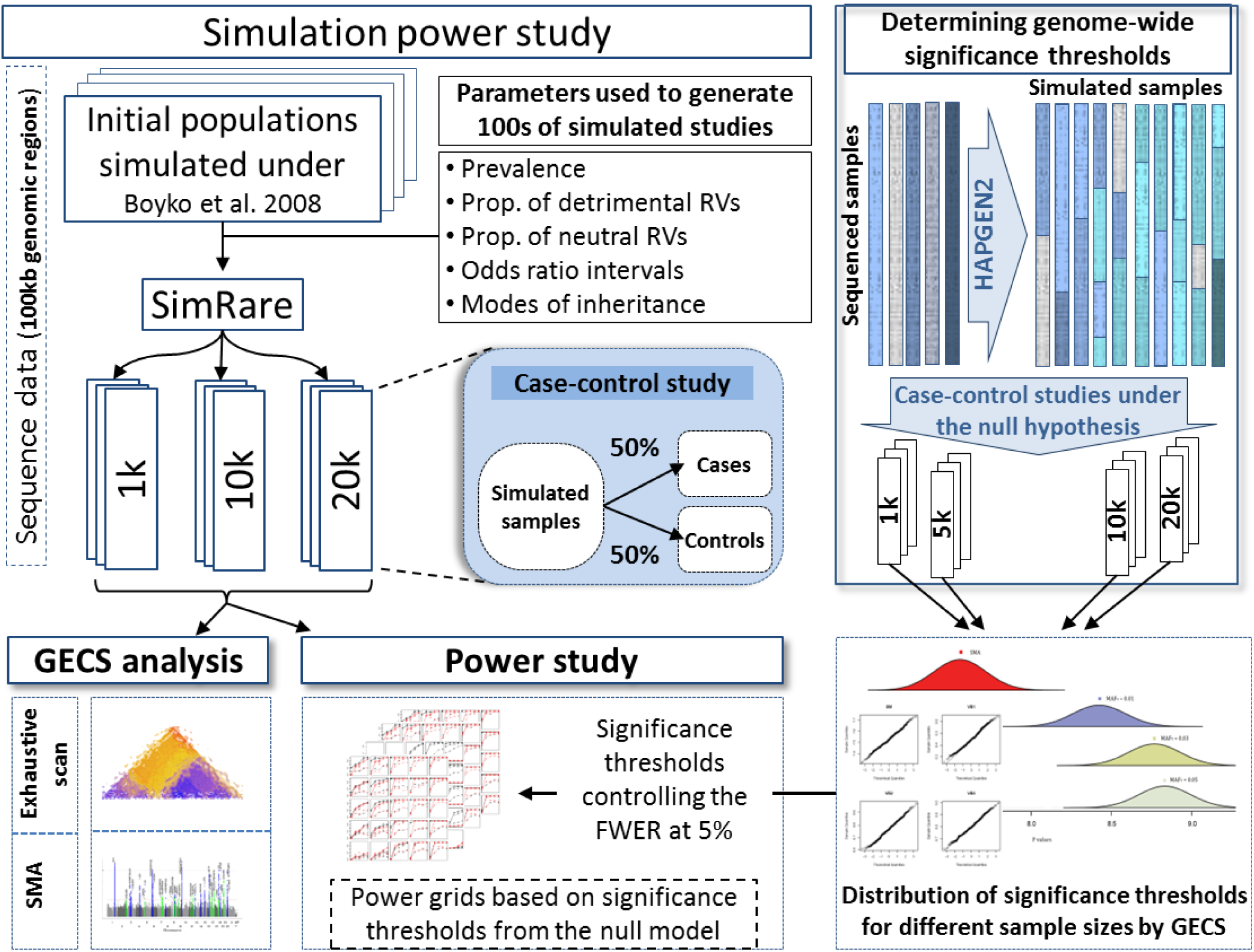
Workflow chart of the entire study illustrating the major steps and procedures. See main text for details.

## Results

### Simulation studies

#### Global significance thresholds

As expected, the GECS-specific thresholds for genome-wide significance were always more stringent than those for the single-marker analysis (SMA) and were inversely correlated with sample size and MAF_T_ (Figure S3-4).

The combined (adjusted for testing at the different MAF_T_) significance thresholds ranged between 7.35×10^-10^ and 2.59×10^-10^ for N=1,000 and N=20,000, respectively.

In real-world data, the combined significance thresholds were 1.87×10^-8^ for the exome dataset (SCZD) and 1.43×10^-9^ for the imputed data (AAMD). The much higher, and therefore less stringent, thresholds were expected due to the incomplete genomic coverage of the available real data. It is expected that in whole-genome, deeply sequenced studies the significance thresholds will be closer to the results obtained in the simulated data (Table 1).

**Table 1:** Empirical, sample-size dependent significance thresholds (α, with control of the FWER at 5%) for simulated genome-wide studies.

| Sample size | Number of replications | SMA | GECS,<br>3 $\text{MAF}_T$<br>combined | GECS,<br>$\text{MAF}_T=0.01$ | GECS,<br>$\text{MAF}_T=0.03$ | GECS,<br>$\text{MAF}_T=0.05$ |
| --- | --- | --- | --- | --- | --- | --- |
| 1,000 | 1,000 | $2.95 \times 10^{-8}$ | $7.35 \times 10^{-10}$ | $3.61 \times 10^{-9}$ | $1.73 \times 10^{-9}$ | $1.60 \times 10^{-9}$ |
| 5,000 | 1,000 | $1.86 \times 10^{-8}$ | $3.31 \times 10^{-10}$ | $1.26 \times 10^{-9}$ | $8.92 \times 10^{-10}$ | $8.49 \times 10^{-10}$ |
| 10,000 | 1,000 | $1.27 \times 10^{-8}$ | $2.81 \times 10^{-10}$ | $1.05 \times 10^{-9}$ | $7.13 \times 10^{-10}$ | $6.91 \times 10^{-10}$ |
| 20,000 | 500 | $1.15 \times 10^{-8}$ | $2.59 \times 10^{-10}$ | $9.28 \times 10^{-10}$ | $6.36 \times 10^{-10}$ | $6.01 \times 10^{-10}$ |

We evaluated the statistical power of GECS, also in comparison to SMA, by simulating realistic case-control sequence data of 2.5-100kb regions under a variety of disease etiology models and sample sizes (Table S1). The exhaustive scan was applied to the simulated region and the significance of the strongest signal was determined using the global significance thresholds (Table 1). Here, we focus on results for PNV=0.3 since results for the different proportions of considered neutral variants (PNV) were similar.

#### Power in models of rare diseases

In small (N=1,000) case-control association studies of diseases with low prevalence (*K*=0.01), GECS performed substantially better than SMA across all considered inheritance modes in the presence of moderate to large proportions of detrimental rare variants (PDV) (Figure 2). In particular, the power ranged between 80% and 99% for high ORs (15≤OR≤ 25) with PDV≥0.3, whereas SMA’s power reached at most 40%. Even in the presence of only small OR values (i.e. 1<OR≤3), as would be expected in complex diseases (BOMBA *et al.* 2017), the power of GECS ranged between 40% (PDV=0.5) and 95% (PDV=0.9). In contrast, the power of SMA exceeded 10% only for PDV>0.3 for all considered OR. Not unexpectedly, small PDV (PDV≤0.3) weakened the performance of GECS in comparison to SMA, where only few rare variants are considered in the study. With PDV=0.1, the power of GECS did not exceed 20% even for the largest OR interval [15, 25] where the power of SMA reached 80%.

**Figure 2:**
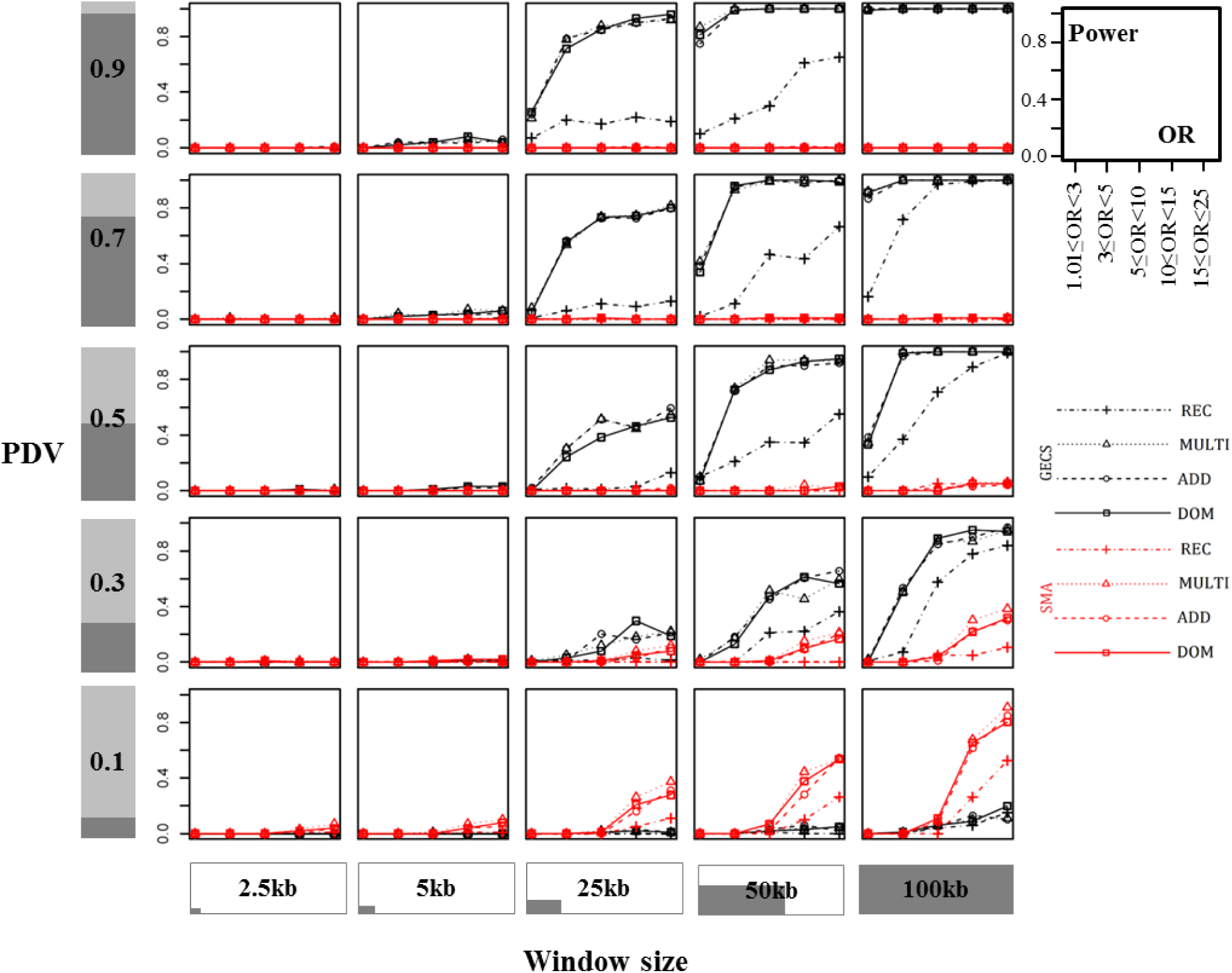
Comparative power analysis for a rare disease (prevalence K=0.01) and small sample size (N=1,000). Results are given for studies with proportion of neutral rare variants (PNV) = 0.3, different simulated window sizes (x-axis), and different proportions of detrimental rare variants (PDV) (y-axis). Black lines: GECS; red lines: SMA. In each grid cell, the power is presented on the y-axis and OR intervals on the x-axis. For an overview see table S20.

For moderate sample sizes (N=10,000), both methods provided comparable power, although of the performance of GECS was clearly superior for small to medium OR and PDV>0.1 (Figure 3). For PDV=0.1, the SMA kept a slight advantage over GECS, although it was much less pronounced than in small-sample studies (N=1,000). A similar observation was made in studies with large sample size (N=20,000): power of both methods were higher for small OR values, but GECS remained much more powerful than SMA with moderate to large PDV values; only low proportions of detrimental rare variants favored the SMA (Figure S5).Increasing PDV always resulted in a power increase for GECS, while SMA remained underpowered (<50% for N=1,000 and 10,000) for small OR values even for the highest proportion of detrimental rare variants (PDV=0.9).

**Figure 3:**
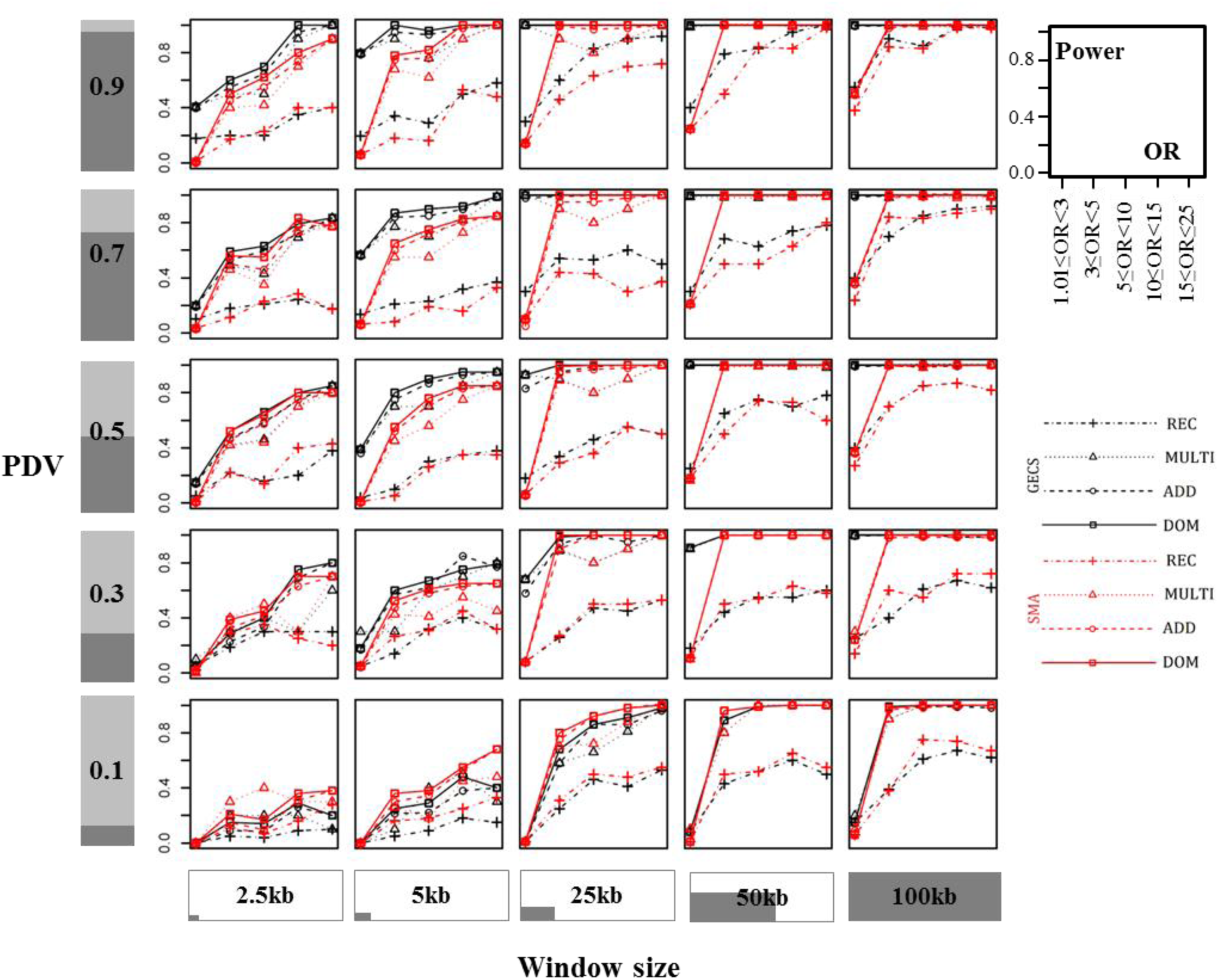
Comparative power analysis for a rare disease (prevalence K=0.01) and moderate sample size (N=10,000). Results are given for studies with proportion of neutral rare variants (PNV) = 0.3, different simulated window sizes (x-axis), and different proportions of detrimental rare variants (PDV) (y-axis). Black lines: GECS; red lines: SMA. In each grid cell, the power is presented on the y-axis and OR intervals on the x-axis. For an overview see table S20.

The impact of the assumed inheritance mode was broadly similar for the dominance (DOM), additive (ADD) and multiplicative (MULTI) genotypic inheritance modes, whereas the recessive mode (REC) resulted in the lowest power values for both GECS and SMA. The difference reached up to 40% for studies with small OR values (< 3). This is not surprising since homozygous carriers of rare variants are expected to be very rare. Extreme PDV values resulted in highly similar power values of about 10% (PDV=0.1) and about 100% (PDV=0.9) for all four modes, while intermediate values, ranging between 0.3 and 0.7, resulted in more pronounced differences between the recessive mode and the other three modes.

#### Power in models of common diseases

Applying GECS and SMA to diseases of higher prevalence (*K*=0.1) yielded some similarities to the results for rare diseases (*K*=0.01), but also some marked differences. In general, the power of both approaches slightly decreased.Differences between the two prevalence classes where most pronounced for small sample sizes (N=1,000) and a recessive inheritance mode (detailed results are shown in Supplementary Note (e)).

### Real-data analysis

#### Advanced age-related macular degeneration (AAMD)

We applied GECS to the whole-genome imputed case-control data of the subset of samples with European ancestry and cases with AAMD from the international AMD genomics consortium (Table S2-3). The analysis was conducted for three MAF thresholds, and the genome-wide significance threshold in the combined study equaled 1.43×10^-09^ (Table 2, Figures S13-16). Previously, strong signals were found in genetic regions on chromosomes 1, 3, 4, 6, 19 and 22, mostly from common variants (Table S7, Figure S13). Recently, 16 other regions containing significant association signals with rare variants were reported (FRITSCHE *et al.* 2016) (Table S8). GECS identified more than 100 genomic regions on chromosomes 1, 3, 4, 5, 6, 10 and 19, where bins of rare variants were found to be significantly associated with AAMD (locally validated by SKAT) (Tables 3, S9). The strongest signals were detected in bins overlapping with protein-coding genes, including *HLA-B*, *HLA-DRA*, and *MICB* in chromosome 6, *FYB* in chromosome 5, *CFD*, and *NRTN* in chromosome 9, and *PLEKHA1* in chromosome 10 (Table S10). These genes, among others, are involved in the regulation of the immune system process and innate immune response). The set of genes overlapping significant bins were enriched in the activation of immune response pathway, in particular, the positive regulation of immune response (7.36-fold enrichment, Bonferroni-corrected p-value of 4.4×10^-4^; a complete list of gene ontology results are shown in Table S11). Additionally, GECS re-identified and re-find most of the previously reported rare variant associations with AAMD (e.g. *CFI*, *C3*, *SKIV2L*, *SYN3*, and *C9*) (FRITSCHE *et al.* 2016; GEERLINGS *et al.* 2018) (Table S12). Odds ratios of identified bins ranged between 0.5 and 3.45, indicating that carrier status can be both positively and negatively correlated with AAMD. However, significant bins with OR>1 were overrepresented on chromosome 6, with OR values ranging between 1.1 and 1.4 and bin sizes ranging between 2 and 26 rare variants.

**Table 2:** Significance thresholds (α, with control of the FWER at 5%) for the whole-genome, imputed AAMD data set, and the whole-exome SCZD data set.

| **Data set** | **SMA** | **GECS,**<br>**3 MAF<sub>T</sub>**<br>**combined** | **GECS,**<br>**MAF<sub>T</sub>=0.01** | **GECS,**<br>**MAF<sub>T</sub>=0.03** | **GECS,**<br>**MAF<sub>T</sub>=0.05** |
| --- | --- | --- | --- | --- | --- |
| AAMD | $1.81 \times 10^{-8}$ | $1.43 \times 10^{-9}$ | $7.42 \times 10^{-9}$ | $2.80 \times 10^{-9}$ | $2.54 \times 10^{-9}$ |
| SCZD | $8.32 \times 10^{-7}$ | $1.87 \times 10^{-8}$ | $4.35 \times 10^{-8}$ | $3.59 \times 10^{-8}$ | $2.84 \times 10^{-8}$ |

**Table 3:** Bins with the locally most significant association signals in AAMD and SCZD data sets, detected by GECS and verified by SKAT. Each bin is the most significant signal in the block of all overlapping significant bins detected by GECS. These bins are verified by SKAT, adjusted for sex, age, 10 principal components, and common variants in physical proximity, if available (p’-values). For verification with SKAT, we set the threshold at 5×10^-8^ for AAMD and 2×10^-6^ for SCZD.

| Data set | Chr. | Bin position (hg19) |  | Locus name (gene) | MAF<sub>T</sub> | Number of RVs | OR | p-value | p'-value |
| --- | --- | --- | --- | --- | --- | --- | --- | --- | --- |
| AAMD | 6 | 31,935,392 | 31,937,762 | *DXO*,<br>*SKIV2L* | 0.03 | 28 | 0.55 | $6.24 \times 10^{-76}$ | $4.74 \times 10^{-81}$ |
| | 6 | 31,878,006 | 31,878,721 | *C2* | 0.05 | 5 | 0.53 | $3.78 \times 10^{-80}$ | $1.23 \times 10^{-70}$ |
| | 10 | 124,226,492 | 124,249,185 | *HTRA1* | 0.05 | 64 | 0.62 | $2.09 \times 10^{-84}$ | $2.96 \times 10^{-30}$ |
| | 19 | 6,718,146 | 6,718,155 | *C3* | 0.03 | 2 | 2.98 | $7.87 \times 10^{-27}$ | $6.30 \times 10^{-28}$ |
| | 6 | 31,473,707 | 31,474,883 | *MICB* | 0.05 | 6 | 1.27 | $1.71 \times 10^{-10}$ | $2.08 \times 10^{-13}$ |
| | 6 | 31,323,455 | 31,323,745 | *HLA-B* | 0.05 | 12 | 1.18 | $3.48 \times 10^{-10}$ | $2.76 \times 10^{-11}$ |
| | 4 | 110,685,721 | 110,685,820 | *CFI* | 0.01 | 5 | 3.42 | $2.15 \times 10^{-11}$ | $7.03 \times 10^{-10}$ |
| | 6 | 31,373,445 | 31,373,957 | *MICA* | 0.05 | 9 | 1.3 | $5.74 \times 10^{-12}$ | $1.03 \times 10^{-09}$ |
| | 5 | 39,199,134 | 39,199,134 | *FYB* | 0.03 | 1 | 1.75 | $2.40 \times 10^{-10}$ | $1.70 \times 10^{-10}$ |
| | 5 | 39,327,884 | 39,327,888 | *C9* | 0.03 | 2 | 1.74 | $4.58 \times 10^{-12}$ | $1.28 \times 10^{-11}$ |
| SCZD | 15 | 73,044,829 | 73,044,833 | *ADPGK* | 0.03 | 7 | 0.5 | $3.75 \times 10^{-20}$ | $4.30 \times 10^{-20}$ |
| | 15 | 73,029,268 | 73,044,833 | *ADPGK* -<br>*BBS4* | 0.03 | 5 | 0.5 | $4.88 \times 10^{-20}$ | $6.73 \times 10^{-19}$ |
| | 17 | 49,239,143 | 49,239,143 | *NME1* | 0.01 | 1 | 0.3 | $2.72 \times 10^{-09}$ | $5.58 \times 10^{-11}$ |
| | 19 | 8,999,386 | 9,028,410 | *MUC16* | 0.03 | 62 | 1.3 | $2.59 \times 10^{-09}$ | $3.10 \times 10^{-10}$ |
| | 22 | 17,687,954 | 17,688,129 | *CERC1* | 0.01 | 9 | 0.3 | $3.79 \times 10^{-13}$ | $6.77 \times 10^{-14}$ |

Notably, bin 6.I (chr6:31,323,455-31,323,745bp, hg19) of 12 rare variants (MAF≤0.05) was found to be significant with a p-value of 3.48×10^-11^, p’-value of 2.76×10^-10^ and OR of 1.2. This bin overlaps with the protein coding human leukocyte antigen B (*HLA-B*), which plays a very important role in the immune system (Figure S17). Interestingly, a previous study found a positive correlation between the *HLA-B* allele HLA-B27 with AAMD (VILLEGAS BECERRIL *et al.* 2009). Also, bin 6.II (chr6:31,473,707–31,474,883bp) overlapped with the *MICB* gene and comprised 6 rare variants (MAF≤0.05). This bin was found to be significantly associated with AAMD with p-value 1.71×10^-10^, p’-value of 2.08×10^-13^ and OR=1.3 (Figure S18). An example for a bin with OR<1 is 10.I (chr10: 124,226,492–124,249,185bp), which comprised 64 rare variants (MAF≤0.05), was found to be associated with AAMD with p-value of 2.09×10^-84^, p’-value of 2.96×10^-30^ with OR=0.6. Notably, this bin, with an apparently protective effect of rare alleles overlaps with *HTRA1,* which has been functionally studied in the context of AMD (NG *et al.* 2012). The association signal was independent from multiple common variants found to be associated with AAMD in this gene (LIANG *et al.* 2012; MCKIBBIN *et al.* 2012). Another noteworthy finding was bin 6.IV (chr6: 31,878,006 - 31,878,721bp) with 5 rare variants in the *C2* gene (MAF≤0.05) was found to be significantly associated with AMD with p-value 3.78×10^-80^, p’-value 1.23×10^-70^, but with OR=0.5 (Figure S19). Our finding is in line with the known role of some protective haplotypes in the *C2*-*AS1* region were found to be significantly reducing the risk of AMD (GOLD *et al.* 2006). For more results see Supplementary Note (f).

#### Schizophrenia

Gene-disruptive and putatively protein-damaging rare variants have been found to be enriched in individuals with schizophrenia (GENOVESE *et al.* 2016; PURCELL *et al.* 2014). We applied GECS to the WES variant data from a population-based schizophrenia Swedish case-control cohort (Table S5). The analysis was conducted for three MAF thresholds, and the genome-wide significance threshold in the combined study comprised 1.87×10^-08^ (Table 2, Figures S21-24). Most of the alleles identified to be significantly associated to schizophrenia had OR<1, so that the carrier status appeared to be protective (Tables 3, S15). For example, bins like 15.I (chr15:73,044,829-73,044,833bp), 17.I (chr17:49,239,143-49,239,143bp) and 22.I (chr22:17,687,954-17,688,129bp) overlapped with genes on chromosome 15 (*ADPGK*), 17 (*NME1*, *NME2*) and 22 (*CERC1*) (Table S16). These genes are involved in the purine nucleoside triphosphate biosynthetic process, which has previously been demonstrated as being strongly linked to the development of schizophrenia (FUMAGALLI *et al.* 2017). On the other hand, bin 19.I (chr9:8,999,386-9,028,410bp), comprising 62 rare SNPs (MAF≤0.03), was found to be significant, with p-value 2.59×10^-09^, p’-value 3.11×10^-10^ and OR=1.3, covering exonic regions of the *MUC16* gene (Figure S25). Although some rare alleles in *MUC16* were reported in association to schizophrenia, none of the 62 rare alleles in this bin were reported before. Moreover, genes covered by bins 15.I, 17.I, 19.I, and 22.I were found to have a function in the small molecule metabolic processes. Interestingly, gene *PRSS3* was covered by bin 9.I (9:33,796,672-33,798,630) comprising 20 rare variants (MAF≤0.05), p’-value of 3.89×10^-11^ and OR=1.4. This gene was not previously reported to be related to schizophrenia. The relatively small sizes of the detected significant bins in the WES data of schizophrenia indicate that the availability of large whole-genome sequencing studies will enable a considerable power gain for our method (Table S17).

#### Benchmarking of GECS for real-world data sets

GECS was found to be feasible for large data sets (Table 18). Analyzing the imputed whole-genome data of AMD, comprising 27,259 samples and round 900,000 variants, took less than 4 hours when analyzing each chromosome in parallel. On average, GECS required fewer computational resources than SMA, with memory usage ranging between 1-5 GB for GECS and about 14 GB for SMA (Table S19). The analysis of the schizophrenia WES data (∼10,000 samples, ∼300,000 SNPs) took at maximum 14h for GECS and 6 h for SMA (Table S19) (for more details see Supplementary Note (g)).

## Discussion

While genome-wide scans with heuristically predetermined analysis regions are an established approach, they are limited in their scope, resolution and power by requiring a prior choice of the analysis regions. In the context of selection analysis, Akey fittingly compared the scan to a hatchet and called for more refined scalpel-like approaches (AKEY 2009). We argue that in Akey’s analogy, the exhaustive scan is an electron microscope, as it allows for base-pair level analysis of genomic regions, with genome-wide, non-conservative, optimally powerful correction for multiple testing using replicates of the data generated under the null hypothesis.

GECS is scalable to large association studies of imputed and sequenced variant data, as demonstrated by our simulation of the null model. The efficiency of our implementation allowed us to estimate significance thresholds for rare-variant analysis in whole-genome sequenced data for association studies comprising up to 20,000 individuals. As a by-product, the analysis offered another opportunity to study significance thresholds (FWER control at 5%) for single-marker analysis (SMA), which, even for small sample sizes of N=1,000 was found to be stricter (α=2.95 ×10^-8^) than the commonly used threshold of α=5.0×10^-8^. This result is consistent with previously published results (AUTON *et al.* 2015; PULIT *et al.* 2017) and highlights the need to abandon the “agreed-upon” significance threshold of 5.0×10^-8^, which is anticonservative for large-scale association studies.

As expected, we found the region-based collapsing test to have stricter significance levels than SMA, ranging between α=3.61×10^-9^ and α=1.60×10^-9^ on average for N=1,000 and MAF_T_ equaling 0.01 and 0.05, respectively, which corresponds to 13.9–31.2 million independent tested hypotheses. These estimates of α allow us to assess the absolute power of the region-based exhaustive scan in future whole-genome deeply sequenced data sets. In contrast to previous studies (ZAWISTOWSKI *et al.* 2010), the power study is free from the assumption that the simulated region and the analysis region happen to coincide. Since the exhaustive scan is guaranteed to identify the most strongly associated regions, our FWER control accounts for the multiple-testing “cost” of finding these regions, which was ignored in previous studies. Overall, the power of GECS is higher, or at least comparable to SMA for small to moderate odds ratios of associated rare variants (1.01≤OR<3), being the OR range expected to be most commonly found in complex diseases. For large sample sizes and large effect sizes, GECS, in general, offers no advantage to detect association. This result reflects the expectation that given a large sample size, enough rare alleles will be present to detect associated variants with sufficient power in single-variant tests (AUER and LEAL 2017).

We applied GECS to real-world data sets, namely of AAMD (imputed microarray data) and of schizophrenia (WES), and performed very stringent quality control of both sets to avoid possible type I errors. Application of GECS to AAMD confirmed a multitude of previously reported rare associated SNPs, for which SMA was underpowered to pick up many signals due to the low MAFs. We confirmed that exhaustively scanning for association through all possible combinations of contiguous rare variants from different MAF thresholds alleviates the limitations posed by previous fixed-bin strategies. The in-depth follow-up analysis showed high enrichment of genes covered by identified bins in pathways with key roles in the development and function of immune system. This is consistent with previous findings, as a putative role for the immune system in the pathogenesis of AMD has been suggested since the 1980s (NUSSENBLATT *et al.* 2014). Furthermore, a locally exhaustive analysis with SKAT uncovered some bins with more extreme p-values than those detected by GECS, such as in bin 4.I. This is expected, since GECS considers the carrier status only and therefore has a low “resolution” in the space of allele counts. SKAT, however, is sensitive to the allele counts, so that bins that have identical contingency tables in GECS test can have differing association testing results with SKAT. On the other hand, a truly exhaustive SKAT analysis is computationally not feasible, so that GECS is required for identification of regions that can be followed up with SKAT locally.

Our approach was also successfully applied to the schizophrenia data set, uncovering previously unreported associations, notably with the *PRSS3* gene. However, judging by the limited spatial extent of the resulting bins, the approach might be underpowered due to limited coverage of the genome in WES studies and will probably improve with availability of WGS data.

In summary, GECS is a powerful approach for detecting phenotypic association of genomic regions harboring rare variants and for refining our understanding of their contribution to predisposition for complex diseases. We conclude that our approach is well-suited for whole-genome and whole-exome association analyses. However, GECS utilizes the simple allele counting function of COLL to achieve perfect, essentially base pair-level spatial resolution. As COLL is only able do dichotomize individuals by the carrier status, the test is not able to distinguish between carriers of one or more minor alleles. We alleviated the limitations of COLL by performing follow-up analysis of candidate regions with locally exhaustive scans using SKAT. However, enabling the exhaustive scan with more sophisticated tests that take more sources of information into account, like allele counts and covariates, might reveal further associated candidate regions. The challenge of extending the exhaustive scan approach to more complex association tests is purely computational in nature. Our algorithm does not generalize to other published association tests in a straightforward manner, so that new solutions will be required to generalizing the exhaustive association scan beyond the collapsing method.

Application of exhaustive scans is not limited to association testing and could be useful in further applications, in particular for studying methylation and evolutionary selection. In fact, our preliminary results show that the exhaustive scan is feasible for the study of selection when used with site frequency spectrum (SFS) based tests such as Tajima’s D (data not shown). This is due to the fact that the computational complexity of SFS-based tests is independent from the number of individuals in the study, since only allele count data is required. As a consequence, the quadratic space of all contingent regions can be computed by brute force, even for very large data sets. Moreover, modern, efficient coalescent simulators such as msprime (KELLEHER *et al.* 2016) and fastsimcoal2 (EXCOFFIER *et al.* 2013) can be used to simulate the null model under neutral evolution under realistic demographic histories (GAZAVE *et al.* 2014) which can be used for FWER-controlled p-values.

In summary, we developed a method that allows for an exhaustive scan of all possible contiguous genomic regions with the collapsing test and eliminates the choice of candidate bins. Instead, the space of all possible bins is tested. This eliminates binning as a source of type II error and is expected to improve power. Furthermore, the speed-up by several orders of magnitude allows for computation of non-conservative genome-wide significance thresholds by permutation, leading to improved power when compared to conservative correction methods such as Bonferroni’s. We show that GECS indeed improves statistical power in both simulated and empirical data sets.

## Methods

### Genomic Exhaustive Collapsing Scan (GECS) algorithm

The collapsing test COLL dichotomizes individuals upon their carrier status of at least one rare allele and applies a simple χ^2^ test to the resulting 2×2 contingency table in a case-control study design (LI and LEAL 2008) (Supplementary Note (a)).The test only considers those variants whose minor allele frequency is below a pre-determined threshold (MAF_T_).We propose an exhaustive region-agnostic whole-genome scan approach that avoids pre-determination (and probable misspecification) of bins by considering all possible contiguous regions at a chromosome. The computational burden, usually prohibitive with human and other large genomes, is solved for COLL by identifying overlapping bins that are identical with respect to their variant carrier status in cases and controls, and skipping redundant computations. More specifically, consider a study with *n* variants with MAF less or equal to a fixed MAF_T_ being present at a chromosome. When conducting COLL, we can parameterize the data set using binary arrays *v*_1_*, v*_2_, … *, v*_*n*_, each of length *N*, where *N* denotes number of samples and the elements *v*_*i,l*_ indicate the carrier status of the *l*-th individual (1=carrier, 0=non-carrier) at the i*-th* variant. Since the analyzed variants are rare, the arrays *v*_*i*_ are sparse (most entries equal 0). We note that, although MAF_T_ is usually used to define “rareness”, in the setting of COLL, we use the number of carriers 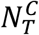, as COLL does not distinguish between homozygous and heterozygous rare-allele carries. By using the cutoff 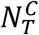, we define a fixed maximum count of 1’s in the arrays *v*_*i*_. For a given MAF_T_, the corresponding 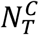 can be estimated from the Hardy-Weinberg proportions as

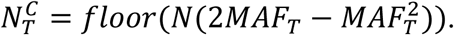

For a bin comprising multiple variants, the logical OR operator yields the array with the carrier status of the individuals with respect to this bin from the constituent variant vectors:

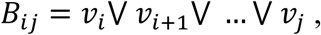

where *i* denotes the index of the start, or left, variant and *j* is the index of the end, or right, variant of the bin. Using the known affection status of the individuals in the study, contingency tables and test statistics *T*_*ij*_ *= T*(*B*_*ij*_) can be obtained from binary arrays *B*_*ij*_ in a computationally efficient way. The space of all combinations of bin start and end positions on a linear chromosome can be considered as a matrix *B* with elements *B*_*ij*_ (Figure S1). Note that diagonal elements represent single-variants, *B*_*ii*_ *= v*_*i*_, and *B*_1*n*_ contains the carrier status with respect to all rare variants on the chromosome. Since *B* is symmetric (*B*_*ij*_ *= B*_*ji*_), only the upper triangular part of *B* with *n(n + 1)/2* elements requires consideration.

Despite binary encoding and utilization of computationally efficient Boolean operators, computing *B* and performing association testing by brute force is not feasible for large data sets. However, the computational space can be considerably reduced, typically by several orders of magnitude. Our algorithm utilizes the fact that a given element *B*_*ij*_ does not change by adding a variant that does not contribute new carriers, i.e. *B*_*ij*_ *= B*_*i*(*j*+1)_. Likewise, removing the first variant *B*_*ij*_ *= B*_(*i*+1)*j*_ often results in an identical set of carriers and, subsequently, invariant contingency table and test statistic. Extrapolating this observation, we find that classes of overlapping bins can be found that are identical with respect to the collapsing test under variation of start and end coordinates. For such clusters of locally non-distinct bins (with respect to the binary representation), the test statistic needs to be computed only once. Therefore, the problem of computing the whole matrix *B* can be reduced to computing all locally distinct bins *B*_*ij*_, in which a locally distinct bin represents a whole cluster of overlapping bins with identical contingency tables. Moreover, for practical purposes we do not need to keep track of the whole range of physical positions under which a bin remains invariant, it is sufficient to keep track of the coordinates of an arbitrary region representing the whole cluster. The set of locally distinct elements of *B* can be found very efficiently by systematically traversing the matrix (w.l.o.g. row-wise) and exploiting properties of the logical OR operation that allow to perform early abandoning of a row while being sure that no distinct bin will be missed in the non-computed elements of the matrix. The crucial observation is that early abandoning can be performed if a bin *B*_*ij*_ is equal to *B*_(*i*+1)*j*_, i.e. the exact element below in the matrix. The condition *B*_*ij*_ *= B*_(*i*+1)*j*_ ensures that *B*_*i*(*j*+*k*)_ *= B*_(*i*+1)(*j*+*k*)_ for all *k ≥ 0*. In other words, if the condition is encountered once, it will be satisfied for the remainder of the row, which is a direct consequence of OR operator’s truth table. Intuitively, if the minor allele of the start variant *v*_*i*_ is not carried by any individual that does not already carry at least one minor allele in *B*_(*i*+1)*j*_, then *v*_*i*_ will not contribute any new carriers to bins with an incremented end position (*j+k)*. Therefore, all distinct bins not yet encountered in row *i* will be encountered in the next row, and may be in the following rows. This observation allows eliminating redundant computations inductively. Early abandoning can also be performed if a bin is found in which all individuals are carriers, i.e. the encoding binary array consists of only 1’s, *B_ij_* = 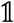, since the array remains “fully collapsed” for the remainder of the row. In pseudocode, the algorithm can be summarized as follows (see Supplementary Note (b) for comprehensive example of the algorithm and its implementation).

~~~
*for* (*i=0; i<n; i++*){
    for (j=i, j<n, j++){
      if (***B*_*ij*_==*B*_(*i*+1)*j*_**|| ***B*_*ij*_==1**) break;
      else if (***B*_*ij*_==*B*_*i*(*j*+1)_**) continue;
      else compute ***T*(*B*_*ij*_)** // locally distinct bin identified
     }
}
~~~

We implemented this algorithm in our publicly available analysis software GECS (https://github.com/ddrichel/GECS) using C++.

Finally, we note that although we perform our analysis under either one or multiple fixed thresholds 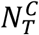, it is possible to generalize the algorithm to consider all possible thresholds 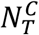, i.e. to simultaneously perform a variable threshold (VT) analysis (see Supplementary Note (c)). However, the algorithm of GECS combined with VT approach is in general not efficient enough to handle large-scale association studies. We therefore resort to applying GECS with three fixed 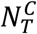. Control of the FWER can be performed by combining the results from the three separate permutation runs under different MAF_T_s (see Supplementary Note (d)).Following (KULLDORFF 1997), we aim to perform at least 9999 permutation replicates for significance testing at alpha=0.05, however for large data sets we sometimes only use 999 permutations.

### Genome-wide significance thresholds for rare-variant analysis

The analysis of large genomic data sets involves simultaneous testing of many hypotheses, whose dependency structure may be unknown. Population history is a strong determining factor for this structure, as it induces specific patterns of genetic variation and allelic correlation. A genome-wide significance level α for analyzing rare variants in genomic data to control type I error will depend on the studied data set and the used statistical test. Given that the number of rare variants is high and that the number of possible bins grows quadratically with the number or rare variants, it can be expected that an exhaustive analysis of all possible contiguous subsequences will require smaller α than with tests of single common variants. For single-marker analysis, the problem of determining significance thresholds that control the FWER has been approached analytically (DICKHAUS and STANGE 2013) and by using Monte-Carlo (MC) simulations (PULIT *et al.* 2017). In realistic scenarios, the analytical approach has to rely on approximations, possibly resulting in a conservative bias. On the other hand, the MC approach results in a quickly converging approximation of the exact thresholds, under the condition that the computational burden can be overcome (ATANASSOV and DIMOV 2008). In this study, we used the maxT/minP approach by (WESTFALL P. H. 1993) in order to control the FWER at the 5% level, performing 999 permutations with respect to the case-control labels. When conducting an analysis with multiple MAF_T_, maxT/minP is obtained across MAF_T_ levels in each permutation, subsequently determining the combined significance level.

This approach can be used also if the analysis is conducted in different chromosomes separately (for more details see Supplementary Note (d)).

### Estimation of genome-wide significance levels

In order to explore the distribution of the test statistic under the null hypothesis of no phenotypic association in a case-control study design, we simulated genome-wide WGS variant data sets representing populations of European (EUR) ancestry using HAPGEN2 software (SU *et al.* 2011) and the 99 CEU (Utah residents with Northwestern European ancestry) individuals from the 1000 Genomes Project Phase 3 as reference (AUTON *et al.* 2015) (ftp://ftp.1000genomes.ebi.ac.uk/vol1/ftp/release/20130502/; accessed October 2017).

HAPGEN2 maintains the frequency distribution and linkage disequilibrium (LD) structure in the simulated data sets similar to that observed in the reference (PULIT *et al.* 2017). We repeatedly simulated data sets comprising 1,000 (“1k”), 5,000 (“5k”), 10,000 (“10k”) and 20,000 (20k”) individuals, respectively, totaling 500 replications for 20k and 1,000 replications for all other each sample sizes. Each data set was partitioned into “cases” and “controls” of equal size by random assignment. We then derived empirical thresholds for genome-wide significance by applying GECS under a threshold 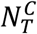 corresponding to minor allele frequencies of MAF_T_ = 0.01, 0.03 and 0.05, respectively. For comparison, COLL was applied to single-marker analysis (SMA) without imposing a frequency threshold and results were compared to those of GECS, delivering a base-line comparison for the performance of our method. Note that under COLL, the SMA analysis is equivalent to the genotypic χ^2^ test under the dominant model. The global 5% significance threshold for each combination of 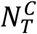 and sample size was estimated by the average 95% quantile for the test statistic across all replications.

### GECS power and performance study

#### Power assessment

In order to assess the power of GECS, we simulated autosomal local DNA sequences (2.5-100kb) under different disease models in a case-control study design.Simulating locally is justified by the expectation that true association signals will be small compared to whole chromosomes. Since GECS is guaranteed to find all distinct subsequences, we do not need to simulate association signals within whole genomes. Instead, we applied the exhaustive scan for association on the simulated subsequences and used the sample-size and variant-frequency specific genome-wide significance thresholds. Again, we also performed single-marker association (SMA) tests for comparison. More specifically, we repeatedly generated genomic regions of 100kb in forward-time simulated populations using SimRare (LI *et al.* 2012), while considering a range of disease models with multiple causal variants in these regions. First, we obtained 25 replications of variant pools from the forward-time simulations following a European simple bottleneck model (BOYKO *et al.* 2008) as demographic history with an ancestral effective population size of N_anc_=7,895, a subsequent bottleneck with N_bot_=5,699 and a current effective population size of N_exp_=30,030, with an exponential growth duration of Gen_exp_=7,703 generations, a mutation rate of *μ*=2×10^-8^ and no recombination. In the second step, we repeatedly and randomly picked a variant pool and simulated a sample set comprising equal numbers of cases and controls under different disease models. Common variants (MAF≥0.05) in the simulations were given an OR of 1.Sample sets were parameterized in terms of: (i) proportion of functional detrimental rare variants (PDV), with values 0.1, 0.3, 0.5, 0.7, 0.9; (ii) proportion of considered neutral rare variants (PNV), with values 0.1, 0.3; (iii) intervals of odds ratios (OR) of detrimental rare variants as a function of their minor allele frequency (MAF), namely [1, 3], [3, 5], [5,10], [10,15], [15,25]; (iv) mode of inheritance, namely, dominant (DOM), recessive (REC), additive (ADD), and multiplicative (MULTI); and (v) prevalence values *K* of 0.01 and 0.1 for the modeled trait (Table S1). Furthermore, we considered sample sizes of 1000, 10,000, and 20,000 individuals. We restricted our analysis to variants with MAF_T_ ≤ 0.01, corresponding to 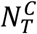 equaling 19, 199 and 398 for the three sample sizes, respectively. For most combination of parameters and sample sizes, 100 simulations were performed and GECS as well as a SMA test were applied to test for phenotypic association (see Table S1 for more details). Obtained nominal p-values were compared against simulation-based sample-size and frequency-specific global significance thresholds.

#### Feasibility of GECS

We benchmarked GECS with respect to (i) the average reduction rate, defined as ratio of the numbers of actually calculated bins to those theoretically existing with a given set of variants, and (ii) the computational memory and time requirements. Since the number of variants included in the analyses depended on the applied MAF_T_ (MAF_T_=0.5 for SMA; MAF_T_=0.01, 0.03, 0.05, respectively, for GECS), it is useful to consider the number of performed tests for association in each study. The computation was conducted on Cheops, a high-performance cluster of University of Cologne, Germany (https://rrzk.uni-koeln.de/cheops.html). Nodes used for the calculations possess 24 GB RAM and CPUs with 2.66 GHz.

### Real-world data set analysis

We applied GECS to two previously analyzed data sets using three different MAF thresholds and corrected for multiple testing using the combined significance thresholds. For each corrected p-value, in the results of the analysis of AAMD and SCZD data, the upper and lower limits of Wilson score binomial interval were calculated for 95% confidence interval. Subsequently we performed an in-depth analysis of the significant regions (Figure S2). The in-depth analysis was performed locally using SKAT (WU *et al.* 2011), as implemented in the SKAT R package (v 1.3.2.1), in two stages, the first of which was a locally exhaustive SKAT (“eSKAT”) with adjustment for age, sex and the first 10 principal components (p*-values).SKAT uses single-SNP score test statistics and a default weighting scheme based on the family of *Beta(x, α, β) ∼ x^α-1^(1-x)^β-1^* distributions which assigns significantly higher weights to variants with low MAFs. In addition, candidate regions with p*-values that indicated possible association were re-analyzed with SKAT and common variants in physical proximity as additional covariates (p’-values). The last step was performed to assure that the association in the candidate region could not be explained simply due to linkage disequilibrium with an associated, common variant.

#### Advanced age-related macular degeneration GWAS

Advanced age-related macular degeneration (AAMD) is a neurodegenerative disease of the retina and the leading cause of blindness among people aged 55 years and older in the western world (SMITH *et al.* 2001). The disease is characterized by reduced retinal pigment epithelium (RPE) function and photoreceptor loss in the macula (FRITSCHE *et al.* 2016). AAMD (late stage AMD) includes two morphological sub-types: endovascular AMD (“wet AMD”) and geographic atrophy (“dry AMD”) and typically preceded by clinically asymptomatic earlier stages of AMD (CHAKRAVARTHY *et al.* 2010). A multitude of genes are reported to harbor common genetic variations associated with the disease (GWAS catalogue: https://www.ebi.ac.uk/gwas). Furthermore, the International AMD Genomics Consortium already examined the contribution of rare and common genetic variation and identified 52 independently associated common and rare variants distributed across 34 loci (FRITSCHE *et al.* 2014; FRITSCHE *et al.* 2016). Besides these single-variant signals, gene-based enrichment of very rare coding variants (frequency < 0.1%) was observed in cases, implicating causal roles for *CFH, CFI,* and *TIMP3* in three of the known AMD risk loci. We obtained whole-genome imputed data from a case-control cohort study, where 14,566 patients with AAMD and 12,693 controls were included (FRITSCHE *et al.* 2014) (data deposited in dbGaP database under dbGaP accession: phs001039.v1.p1). Besides more than 12 million variants, the study included 163,714 directly genotyped, mostly rare, protein-altering variants and 8,200 variants in known AAMD-associated loci (FRITSCHE *et al.* 2016). We included only the 27,259 European samples from this study and restricted the phenotype to AAMD cases (Table S2-3). GECS was applied with three MAF thresholds, namely 0.01, 0.03 and 0.05, for defining rare variants. The final analysis was conducted by combining the results from all three thresholds and applying correction for multiple testing (see Methods). In a post-GECS analysis, we categorized all significant bins in blocks constituted by all overlapped bins in a region. Then we determined bins with the lowest p-values in each block, each of them being defined by their chromosomal positions and threshold MAF_T_. Subsequently, these bins were subject to further locally exhaustive association analysis.

#### Schizophrenia Exome Study

Schizophrenia is a chronic debilitating mental disorder with a lifetime risk of about 0.7% and a heritability of 60–80% conferred by common and rare alleles at many loci (LICHTENSTEIN *et* al. 2009; MCGRATH *et al.* 2008). Alleles associated with an extremely high risk are expected to be prevented from reaching even modest allele frequencies due to purifying selection (ZUK *et al.* 2014). Schizophrenia affects approximately 1% of the population worldwide and is characterized by thought abnormalities, hallucinations, delusions, and bizarre behaviors (FREEDMAN 2003). The genetic basis of this complex disorder was subject to numerous genome-wide association studies which continue to uncover common SNPs at novel loci (GWAS catalogue: https://www.ebi.ac.uk/gwas). One notable outcome of these large-scale genome-wide investigations is the degree of polygenicity, with thousands of genes and non-coding loci harboring risk alleles (PURCELL *et al.* 2014). Recently, a Swedish study aimed to investigate the disruptive role of extremely rare variants in schizophrenia by sequencing a very large number of individuals from the same population. They analyzed the exome sequences of about 12,300 unrelated individuals from Sweden and informed this analysis with a much larger set of exome sequencing data from 45,376 individuals from multiple non-psychiatric cohorts ascertained by the Exome Aggregation Consortium (GENOVESE *ET AL.* 2016).

We analyzed WES variant data of 12,380 samples from the schizophrenia and bipolar disorder Swedish case-control cohort (data deposited in dbGaP database under dbGaP accession: phs000473.v2.p2). We included only those 10,898 samples that passed our quality control regarding population stratification (Table S5). Individuals with schizophrenia comprised 4,795 samples of the cases. Four 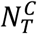 values were considered for GECS analysis, namely 216, 642, 1059, and 3,910, corresponding to MAF_T_ of 0.01, 0.03, and 0.05, respectively. The post-GECS analysis was conducted analogously to AAMD.

## Authors’ contributions

DD conceived the study and demonstrated the proof of concept. GK performed the simulations and the data analysis. DD and GK wrote the analysis software, with contributions by TB. DD supervised the study, with some contributions by TB and MN. GK, DD, MN, and TB wrote the manuscript. All authors read, revised and approved the final manuscript.

## Acknowledgements

The work was supported by the German Research Foundation grant BE 38/28/9-1. The funding organization did not have any influence on the design, conduct or conclusions of the study. We would like to thank Dr. Gabrielle Thorn for her organizing support.

