## Supplementary file for "The exhaustive genomic scan approach, with an application to rare-variant association analysis"

### Tables:

**Table S1: Values of parameters which were used in the power study. Each possible combination of parameters represented a scenario in the study, except for simulations with N=20,000, which were performed for PNV=0.3 and the dominant inheritance mode only**

| **Parameters** | **Values** |
|---|---|
| Prevalence (K) | 0.01, 0.1 |
| PNV | 0.1, 0.3 |
| Sample size | 1,000, 10,000, 20,000 |
| Inheritance mode | dominant (DOM), additive (ADD), multiplicative (MULTI), recessive (REC) |
| PDV | 0.1, 0.3, 0.5, 0.7, 0.9 |
| OR intervals | 1.01-3, 3-5, 5-10, 10-15, 15-25 |
| Size of analysis window | 2.5kb, 5kb, 25kb, 50kb, 100kb |

**Table S2: Description of the AMD data set (build hg19)**

| | All samples | European ancestry | African ancestry | Asian ancestry | Unknown ancestry |
|---|---|---|---|---|---|
| **Number of samples** | 35,358 | 32,637 | 208 | 1,742 | 771 |
| cases | 19,112 | 18,047 | 76 | 649 | 340 |
| controls | 16,246 | 16,106 | 132 | 1093 | 431 |
| women | 20,214 | 18,829 | 890 | 85 | 410 |
| men | 15,144 | 13,808 | 852 | 123 | 361 |

**Table S3: Description of the European AMD data set, used for further analysis**

| | **All data set** | **European** | **European after QC*** |
|---|---|---|---|
| **Sample size** | 35,358 | 32,637 | 27,259 |
| **Controls** | 16,246 | 14,590 | 14,566 |
| **Advanced AMD (AAMD) cases** | 13,379 | 12,711 | 12,693 |
| **Intermediate AMD cases** | 5,733 | 5,336 | - |
| **All AMD cases** | 19,112 | 18,047 | 12,693 |
| **Average age** | ~75 | ~77 | ~72 |
| **Number of SNPs** | 11,722,963 | 11,712,893 | 11,712,830 ** |

* In the quality control stage, intermediate AMD cases and 42 PCA-outliers (see table S4) were excluded.
** Only 428,357 variants were directly genotyped in the data set.

**Table S4: Excluded PCA-outliers in the AMD data set.**

| Nr. | Sample-ID |
|---|---|
| 1 | IAMDGC00769 |
| 2 | IAMDGC05427 |
| 3 | IAMDGC06555 |
| 4 | IAMDGC06574 |
| 5 | IAMDGC08194 |
| 6 | IAMDGC08279 |
| 7 | IAMDGC09965 |
| 8 | IAMDGC11776 |
| 9 | IAMDGC13070 |
| 10 | IAMDGC14221 |
| 11 | IAMDGC14509 |
| 12 | IAMDGC14534 |
| 13 | IAMDGC14967 |
| 14 | IAMDGC15779 |
| 15 | IAMDGC16012 |
| 16 | IAMDGC16119 |
| 17 | IAMDGC16223 |
| 18 | IAMDGC16753 |
| 19 | IAMDGC17010 |
| 20 | IAMDGC18277 |
| 21 | IAMDGC18598 |
| 22 | IAMDGC18648 |
| 23 | IAMDGC19367 |
| 24 | IAMDGC22398 |
| 25 | IAMDGC24181 |
| 26 | IAMDGC25735 |
| 27 | IAMDGC25908 |
| 28 | IAMDGC26258 |
| 29 | IAMDGC26643 |
| 30 | IAMDGC26652 |
| 31 | IAMDGC26702 |
| 32 | IAMDGC28886 |
| 33 | IAMDGC29369 |
| 34 | IAMDGC31174 |
| 35 | IAMDGC32617 |
| 36 | IAMDGC32793 |
| 37 | IAMDGC32830 |
| 38 | IAMDGC33953 |
| 39 | IAMDGC34220 |
| 40 | IAMDGC34288 |
| 41 | IAMDGC34749 |
| 42 | IAMDGC35085 |

**Table S5: Description of the SCZD data set (build hg19).**

| Applied QC-level | Cases | Controls | Female | Male | Total | Total number of variants |
|---|---|---|---|---|---|---|
| None | 6,135 | 6,245 | 5,780 | 6,600 | 12,380 | 1,811,204 |
| SCZD only | 4,969 | 6,245 | 5,069 | 6,145 | 11,214 | 1,650,674 |
| QC* | 4,696 | 6,067 | 4,856 | 5,907 | 10,763 | 308,456 |

* Quality control required non-monomorphic bi-allelic SNPs, a genotype call rate of 0.05 for variants and of 0.04 for individuals, no relatives (expected identity-by-descent sharing ≤0.35), in addition to exclusion of PCA-outliers (see table S6).

**Table S6: Excluded PCA-outliers in the SCZD data set.**

| Nr. | Sample-ID |
|---|---|
| 1 | 28304 |
| 2 | 28309 |
| 3 | 28391 |
| 4 | 28449 |
| 5 | 45265 |
| 6 | 45366 |
| 7 | 45438 |
| 8 | 60620 |
| 9 | 71502 |
| 10 | 75693 |
| 11 | 75695 |
| 12 | 107730 |
| 13 | 141934 |
| 14 | 238481 |
| 15 | 313392 |
| 16 | 355922 |
| 17 | 376954 |
| 18 | 377199 |
| 19 | 443738 |
| 20 | 447150 |
| 21 | 447359 |
| 22 | 447545 |
| 23 | 455200 |
| 24 | 481887 |
| 25 | 481890 |
| 26 | 482599 |
| 27 | 482613 |
| 28 | 513409 |
| 29 | 514030 |
| 30 | 573331 |
| 31 | 595814 |
| 32 | 595854 |
| 33 | 654662 |
| 34 | 654900 |

**Table S7: List of previously reported genes in association to AAMD (FRITSCHE *et al.* 2016).**

| Chr. | Gene | Lead SNP |
|---|---|---|
| 1 | *CFH* | rs10922109 |
| 2 | *COL4A3* | rs11884770 |
| 3 | *ADAMTS9/AS2* | rs62247658 |
| 3 | *COL8A1* | rs140647181 |
| 4 | *CFI* | rs10033900 |
| 5 | *C9* | rs62358361 |
| 5 | *PRLR/SPEF2* | rs114092250 |
| 6 | *C2/CFB/SKIV2L* | rs116503776 |
| 6 | *VEGFA* | rs943080 |
| 7 | *KMT2E/SRPK2* | rs1142 |
| 7 | *PILRB/PILRA* | rs7803454 |
| 8 | *TNRSF10A* | rs79037040 |
| 9 | *TRPM3* | rs71507014 |
| 9 | *MIR6130/RORB* | rs10781182 |
| 9 | *TGFBR1* | rs1626340 |
| 9 | *ABCA1* | rs2740488 |
| 10 | *ARHGAP21* | rs12357257 |
| 10 | *ARMS2/HTRA1* | rs3750846 |
| 12 | *RDH5/CD63* | rs3138141 |
| 12 | *ACAD10* | rs61941274 |
| 13 | *B3GALTL* | rs9564692 |
| 14 | *RAD51B* | rs61985136 |
| 15 | *LIPC* | rs2043085 |
| 16 | *CETP* | rs5817082 |
| 16 | *CTRB2* | rs72802342 |
| 17 | *TMEM97/VTN* | rs11080055 |
| 17 | *NPLOC4/TSPAN10* | rs6565597 |
| 19 | *C3* | rs2230199 |
| 19 | *CNN2* | rs67538026 |
| 19 | *APOE* | rs429358 |
| 20 | *MMP9* | rs142450006 |
| 20 | *C20orf85* | rs201459901 |
| 22 | *SYN3/TIMP3* | rs5754227 |
| 22 | *SLC16A8* | rs8135665 |

**Table S8: List of previously reported rare variants in association with AAMD.**

| Chr. | Position | Locus | rs-ID | Function** | MAF | p-value | Reference* |
|---|---|---|---|---|---|---|---|
| 1 | 196,747,245 | *CFH* | rs121913059 | 1 | 0.0001 | $9.00\times10^{-24}$ | 1 |
| 1 | 196,644,043 | *KCNT2-CFH* | rs148553336 | 2 | 0.003 | $9.00\times10^{-26}$ | 1 |
| 1 | 196,411,028 | *KCNT2* | rs187328863 | 3 | 0.01 | $1.00\times10^{-68}$ | 1 |
| 1 | 196,737,512 | *CFH* | rs35292876 | 4 | 0.01 | $8.00\times10^{-37}$ | 1 |
| 1 | 196,958,651 | *CFH* | rs191281603 | 3 | 0.009 | $7.00\times10^{-07}$ | 1 |
| 3 | 99,461,824 | *LOC105374007* | rs140647181 | 2 | 0.012 | $1.00\times10^{-11}$ | 1 |
| 4 | 109,764,664 | *CFI* | rs141853578 | 1 | 0.0005 | $6.00\times10^{-10}$ | 1 |
| 5 | 39,327,786 | *C9* | rs62358361 | 3 | 0.002 | $1.00\times10^{-14}$ | 1 |
| 5 | 35,494,346 | *PRLR-SPEF2* | rs114092250 | 5 | 0.006 | $2.00\times10^{-08}$ | 1 |
| 6 | 32,107,027 | *TNXB* | rs12153855 | 3 | 0.011 | $1.00\times10^{-09}$ | 3 |
| 6 | 31,946,792 | *STK19* | rs2746394 | 3 | 0.008 | $1.00\times10^{-32}$ | 1 |
| 6 | 31,938,233 | *C2-AS1, C2* | rs9380272 | 3 | 0.012 | $2.00\times10^{-08}$ | 3 |
| 6 | 31,979,250 | *STK19* | rs181705462 | 3 | 0.01 | $3.00\times10^{-10}$ | 1 |
| 12 | 111,694,806 | *ACAD10* | rs61941274 | 3 | 0.01 | $1.00\times10^{-09}$ | 1 |
| 15 | 63,782,903 | *HERC1* | rs74320127 | 3 | 0.05 | $5.00\times10^{-08}$ | 2 |
| 19 | 6,718,135 | *C3* | rs147859257 | 1 | 0.003 | $3.00\times10^{-28}$ | 1 |

*1 Fritsche et al 2016 (FRITSCHE *et al.* 2016); 2: Yan Q 2018 (YAN *et al.* 2018); 3: Cipriani V 2012 (CIPRIANI *et al.* 2012).

** Function abbreviations are: 1: missense variant, 2: intergenic variant, 3: intron variant, 4: synonymous variant, and 5: regulatory region variant.

**Table S9: Bins with the locally most significant association signals in AAMD data set, detected by GECS and verified by SKAT.** Each bin is the most significant signal in the block of all overlapping significant bins detected by GECS. These bins are verified by SKAT, adjusted for sex, age, 10 principal components, and common variants in physical proximity, if available (p'-values). For verification with SKAT, we set the threshold at $5\times10^{-8}$.

| NCT | Chr. | Bin position (gh19) | | Bin size in bp | Number of RVs | p-value | Corrected p-value (p_corr) | Wilson lower CI 95% | Wilson upper CI 95% | OR | p'-value | Overlapping genes |
|---|---|---|---|---|---|---|---|---|---|---|---|---|
| 542 | 4 | 110,685,721 | 110,685,820 | 100 | 5 | $2.15\times10^{-11}$ | 0.002 | $5.49\times10^{-04}$ | $7.27\times10^{-03}$ | 3.43 | $7.03\times10^{-10}$ | *CFI* |
| 1,611 | 1 | 197,814,623 | 197,814,623 | 1 | 1 | $8.08\times10^{-10}$ | 0.002 | $5.49\times10^{-04}$ | $7.27\times10^{-03}$ | 1.453 | $1.20\times10^{-09}$ | |
| 1,611 | 1 | 197,815,382 | 197,816,763 | 1,382 | 2 | $4.30\times10^{-17}$ | 0.001 | $1.77\times10^{-04}$ | $5.65\times10^{-03}$ | 0.553 | $8.17\times10^{-15}$ | |
| 1,611 | 1 | 197,829,433 | 197,832,376 | 2,944 | 14 | $1.65\times10^{-12}$ | 0.001 | $1.77\times10^{-04}$ | $5.65\times10^{-03}$ | 0.573 | $2.09\times10^{-09}$ | |
| 1,611 | 1 | 197,876,319 | 197,880,175 | 3,857 | 10 | $3.69\times10^{-22}$ | 0.001 | $1.77\times10^{-04}$ | $5.65\times10^{-03}$ | 0.733 | $6.93\times10^{-13}$ | *C1orf53* |
| 1,611 | 1 | 197,997,953 | 198,003,207 | 5,255 | 8 | $1.97\times10^{-11}$ | 0.001 | $1.77\times10^{-04}$ | $5.65\times10^{-03}$ | 0.79 | $1.20\times10^{-08}$ | |
| 1,611 | 1 | 198,094,634 | 198,101,964 | 7,331 | 5 | $1.91\times10^{-10}$ | 0.001 | $1.77\times10^{-04}$ | $5.65\times10^{-03}$ | 0.78 | $3.10\times10^{-08}$ | |
| 1,611 | 5 | 39,199,134 | 39,199,134 | 1 | 1 | $2.40\times10^{-10}$ | 0.001 | $1.77\times10^{-04}$ | $5.65\times10^{-03}$ | 1.74 | $1.70\times10^{-10}$ | *FYB* |
| 1,611 | 5 | 39,327,884 | 39,327,888 | 5 | 2 | $4.58\times10^{-12}$ | 0.001 | $1.77\times10^{-04}$ | $5.65\times10^{-03}$ | 1.74 | $1.28\times10^{-11}$ | *C9* |
| 1,611 | 5 | 39,331,894 | 39,331,894 | 1 | 1 | $7.49\times10^{-12}$ | 0.001 | $1.77\times10^{-04}$ | $5.65\times10^{-03}$ | 1.73 | $1.47\times10^{-11}$ | *C9* |
| 1,611 | 6 | 31,024,244 | 31,024,244 | 1 | 1 | $1.42\times10^{-09}$ | 0.002 | $5.49\times10^{-04}$ | $7.27\times10^{-03}$ | 0.71 | $3.56\times10^{-09}$ | *HCG22* |
| 1,611 | 6 | 31,421,170 | 31,421,514 | 345 | 4 | $1.14\times10^{-12}$ | 0.001 | $1.77\times10^{-04}$ | $5.65\times10^{-03}$ | 1.31 | $6.37\times10^{-10}$ | *HCP5* |
| 1,611 | 6 | 31,437,566 | 31,437,566 | 1 | 1 | $1.11\times10^{-09}$ | 0.001 | $1.77\times10^{-04}$ | $5.65\times10^{-03}$ | 0.65 | $4.63\times10^{-10}$ | *HCP5* |
| 1,611 | 6 | 31,440,505 | 31,440,641 | 137 | 2 | $2.09\times10^{-10}$ | 0.001 | $1.77\times10^{-04}$ | $5.65\times10^{-03}$ | 0.63 | $6.85\times10^{-11}$ | *HCP5* |
| 1,611 | 6 | 31,444,187 | 31,445,396 | 1,21 | 7 | $4.65\times10^{-12}$ | 0.001 | $1.77\times10^{-04}$ | $5.65\times10^{-03}$ | 0.779 | $1.43\times10^{-15}$ | *HCP5* |
| 1,611 | 6 | 31,458,400 | 31,460,161 | 1,762 | 9 | $2.99\times10^{-15}$ | 0.001 | $1.77\times10^{-04}$ | $5.65\times10^{-03}$ | 1.27 | $2.13\times10^{-09}$ | |
| 1,611 | 6 | 31,485,262 | 31,487,066 | 1,805 | 7 | $4.81\times10^{-12}$ | 0.001 | $1.77\times10^{-04}$ | $5.65\times10^{-03}$ | 0.72 | $3.48\times10^{-12}$ | |
| 1,611 | 6 | 31,518,764 | 31,519,389 | 626 | 2 | $2.32\times10^{-12}$ | 0.001 | $1.77\times10^{-04}$ | $5.65\times10^{-03}$ | 0.70 | $5.17\times10^{-13}$ | *NFKBIL1* |
| 1,611 | 6 | 31,522,174 | 31,522,518 | 345 | 2 | $1.80\times10^{-12}$ | 0.001 | $1.77\times10^{-04}$ | $5.65\times10^{-03}$ | 0.65 | $2.03\times10^{-12}$ | *NFKBIL1* |

| | | | | | | | | | | | | |
|---|---|---|---|---|---|---|---|---|---|---|---|---|
| 1,611 | 6 | 31,586,761 | 31,592,082 | 5,322 | 23 | $6.95\times10^{-26}$ | 0.001 | $1.77\times10^{-04}$ | $5.65\times10^{-03}$ | 0.72 | $1.14\times10^{-23}$ | *BTNL2* |
| 1,611 | 6 | 31,787,120 | 31,787,539 | 420 | 2 | $5.52\times10^{-14}$ | 0.001 | $1.77\times10^{-04}$ | $5.65\times10^{-03}$ | 0.57 | $9.08\times10^{-16}$ | |
| 1,611 | 6 | 31,799,353 | 31,800,492 | 1,14 | 4 | $2.60\times10^{-14}$ | 0.001 | $1.77\times10^{-04}$ | $5.65\times10^{-03}$ | 1.30 | $4.80\times10^{-13}$ | |
| 1,611 | 6 | 31,935,392 | 31,937,762 | 2,371 | 28 | $6.24\times10^{-76}$ | 0.001 | $1.77\times10^{-04}$ | $5.65\times10^{-03}$ | 0.55 | $4.74\times10^{-81}$ | *DXO SKIV2L* |
| 1,611 | 6 | 32,202,920 | 32,202,920 | 1 | 1 | $2.62\times10^{-15}$ | 0.001 | $1.77\times10^{-04}$ | $5.65\times10^{-03}$ | 0.61 | $3.46\times10^{-15}$ | |
| 1,611 | 6 | 32,216,480 | 32,216,895 | 416 | 3 | $1.03\times10^{-19}$ | 0.001 | $1.77\times10^{-04}$ | $5.65\times10^{-03}$ | 0.69 | $6.96\times10^{-21}$ | |
| 1,611 | 6 | 32,241,969 | 32,242,178 | 210 | 3 | $1.13\times10^{-15}$ | 0.001 | $1.77\times10^{-04}$ | $5.65\times10^{-03}$ | 0.64 | $1.12\times10^{-15}$ | |
| 1,611 | 6 | 32,251,835 | 32,252,170 | 336 | 4 | $3.25\times10^{-13}$ | 0.001 | $1.77\times10^{-04}$ | $5.65\times10^{-03}$ | 0.68 | $7.48\times10^{-13}$ | |
| 1,611 | 6 | 32,270,925 | 32,271,088 | 164 | 2 | $2.12\times10^{-14}$ | 0.001 | $1.77\times10^{-04}$ | $5.65\times10^{-03}$ | 0.59 | $1.47\times10^{-13}$ | *C6orf10* |
| 1,611 | 6 | 32,300,471 | 32,300,824 | 354 | 4 | $1.65\times10^{-13}$ | 0.001 | $1.77\times10^{-04}$ | $5.65\times10^{-03}$ | 0.68 | $3.52\times10^{-16}$ | *C6orf10* |
| 1,611 | 6 | 32,302,546 | 32,302,546 | 1 | 1 | $4.80\times10^{-14}$ | 0.001 | $1.77\times10^{-04}$ | $5.65\times10^{-03}$ | 0.61 | $2.70\times10^{-14}$ | *C6orf10* |
| 1,611 | 6 | 32,304,466 | 32,304,466 | 1 | 1 | $2.45\times10^{-13}$ | 0.001 | $1.77\times10^{-04}$ | $5.65\times10^{-03}$ | 0.62 | $1.59\times10^{-13}$ | *C6orf10* |
| 1,611 | 6 | 32,304,863 | 32,305,102 | 240 | 2 | $3.11\times10^{-13}$ | 0.001 | $1.77\times10^{-04}$ | $5.65\times10^{-03}$ | 0.628 | $2.84\times10^{-13}$ | *C6orf10* |
| 1,611 | 6 | 32,319,173 | 32,319,189 | 17 | 2 | $1.69\times10^{-16}$ | 0.001 | $1.77\times10^{-04}$ | $5.65\times10^{-03}$ | 0.63 | $3.77\times10^{-16}$ | *C6orf10* |
| 1,611 | 6 | 32,340,374 | 32,340,593 | 220 | 2 | $1.61\times10^{-13}$ | 0.001 | $1.77\times10^{-04}$ | $5.65\times10^{-03}$ | 0.68 | $1.52\times10^{-13}$ | |
| 1,611 | 6 | 32,343,086 | 32,343,588 | 503 | 2 | $1.16\times10^{-15}$ | 0.001 | $1.77\times10^{-04}$ | $5.65\times10^{-03}$ | 0.64 | $1.72\times10^{-14}$ | |
| 1,611 | 6 | 32,346,903 | 32,346,903 | 1 | 1 | $2.53\times10^{-13}$ | 0.001 | $1.77\times10^{-04}$ | $5.65\times10^{-03}$ | 0.64 | $6.97\times10^{-13}$ | |
| 1,611 | 6 | 32,351,217 | 32,351,217 | 1 | 1 | $1.76\times10^{-13}$ | 0.001 | $1.77\times10^{-04}$ | $5.65\times10^{-03}$ | 0.64 | $3.30\times10^{-13}$ | |
| 1,611 | 6 | 32,356,684 | 32,356,684 | 1 | 1 | $1.02\times10^{-13}$ | 0.001 | $1.77\times10^{-04}$ | $5.65\times10^{-03}$ | 0.64 | $1.33\times10^{-13}$ | |
| 1,611 | 6 | 32,369,601 | 32,369,601 | 1 | 1 | $5.03\times10^{-14}$ | 0.001 | $1.77\times10^{-04}$ | $5.65\times10^{-03}$ | 0.60 | $6.47\times10^{-15}$ | *BTNL2* |
| 1,611 | 6 | 32,395,997 | 32,396,078 | 82 | 2 | $3.74\times10^{-12}$ | 0.001 | $1.77\times10^{-04}$ | $5.65\times10^{-03}$ | 0.66 | $2.84\times10^{-12}$ | |
| 1,611 | 6 | 32,402,968 | 32,404,276 | 1,309 | 4 | $1.43\times10^{-11}$ | 0.001 | $1.77\times10^{-04}$ | $5.65\times10^{-03}$ | 0.75 | $8.68\times10^{-14}$ | |
| 1,611 | 10 | 124,156,944 | 124,241,923 | 84,98 | 273 | $1.04\times10^{-69}$ | 0.001 | $1.77\times10^{-04}$ | $5.65\times10^{-03}$ | 0.63 | $2.37\times10^{-22}$ | *ARMS2 HTRA1 MIR3941 PLEKHA1* |
| 1,611 | 19 | 6,718,146 | 6,718,155 | 10 | 2 | $7.87\times10^{-27}$ | 0.001 | $1.77\times10^{-04}$ | $5.65\times10^{-03}$ | 2. 99 | $6.30\times10^{-28}$ | *C3* |
| 2,657 | 1 | 197,920,216 | 197,924,770 | 4,555 | 4 | $5.74\times10^{-22}$ | 0.001 | $1.77\times10^{-04}$ | $5.65\times10^{-03}$ | 0.70 | $1.36\times10^{-18}$ | |
| 2,657 | 1 | 197,931,243 | 197,939,490 | 8,248 | 7 | $4.32\times10^{-21}$ | 0.001 | $1.77\times10^{-04}$ | $5.65\times10^{-03}$ | 0.75 | $1.86\times10^{-14}$ | |
| 2,657 | 1 | 198,151,111 | 198,151,688 | 578 | 3 | $5.49\times10^{-13}$ | 0.001 | $1.77\times10^{-04}$ | $5.65\times10^{-03}$ | 0.63 | $7.02\times10^{-10}$ | *NEK7* |

| | | | | | | | | | | | | |
|---|---|---|---|---|---|---|---|---|---|---|---|---|
| 2,657 | 1 | 198,210,986 | 198,216,293 | 5,308 | 18 | $8.50\times10^{-10}$ | 0.002 | $5.49\times10^{-04}$ | $7.27\times10^{-03}$ | 0.81 | $1.62\times10^{-09}$ | *NEK7* |
| 2,657 | 6 | 30,075,877 | 30,076,828 | 952 | 10 | $9.81\times10^{-10}$ | 0.002 | $5.49\times10^{-04}$ | 7.27×10-03 | 0.78 | $1.45\times10^{-11}$ | *TRIM31 TRIM31-AS1* |
| 2,657 | 6 | 31,274,584 | 31,274,734 | 151 | 3 | $7.15\times10^{-14}$ | 0.001 | $1.77\times10^{-04}$ | $5.65\times10^{-03}$ | 0.79 | $1.21\times10^{-11}$ | |
| 2,657 | 6 | 31,322,108 | 31,322,611 | 504 | 6 | $9.80\times10^{-13}$ | 0.001 | $1.77\times10^{-04}$ | $5.65\times10^{-03}$ | 1.28 | $1.62\times10^{-11}$ | *HLA-B* |
| 2,657 | 6 | 31,323,455 | 31,323,745 | 291 | 12 | $3.48\times10^{-10}$ | 0.001 | $1.77\times10^{-04}$ | $5.65\times10^{-03}$ | 1.18 | $2.76\times10^{-11}$ | *HLA-B* |
| 2,657 | 6 | 31,329,415 | 31,331,621 | 2,207 | 16 | $5.77\times10^{-13}$ | 0.001 | $1.77\times10^{-04}$ | $5.65\times10^{-03}$ | 1.23 | $1.57\times10^{-11}$ | |
| 2,657 | 6 | 31,335,791 | 31,335,983 | 193 | 3 | $1.69\times10^{-12}$ | 0.001 | $1.77\times10^{-04}$ | $5.65\times10^{-03}$ | 1.31 | $2.09\times10^{-08}$ | |
| 2,657 | 6 | 31,341,327 | 31,342,419 | 1,093 | 9 | $4.10\times10^{-12}$ | 0.001 | $1.77\times10^{-04}$ | $5.65\times10^{-03}$ | 1.22 | $3.09\times10^{-10}$ | *AL671883.1* |
| 2,657 | 6 | 31,346,655 | 31,347,314 | 660 | 7 | $1.84\times10^{-11}$ | 0.001 | $1.77\times10^{-04}$ | $5.65\times10^{-03}$ | 1.25 | $1.47\times10^{-13}$ | |
| 2,657 | 6 | 31,373,445 | 31,373,957 | 513 | 9 | $5.74\times10^{-12}$ | 0.001 | $1.77\times10^{-04}$ | $5.65\times10^{-03}$ | 1.29 | $1.03\times10^{-09}$ | *HCP5 MICA* |
| 2,657 | 6 | 31,377,662 | 31,378,089 | 428 | 3 | $3.52\times10^{-13}$ | 0.001 | $1.77\times10^{-04}$ | $5.65\times10^{-03}$ | 1.28 | $1.98\times10^{-09}$ | *HCP5 MICA* |
| 2,657 | 6 | 31,378,864 | 31,379,016 | 153 | 5 | $1.76\times10^{-13}$ | 0.001 | $1.77\times10^{-04}$ | $5.65\times10^{-03}$ | 1.30 | $1.07\times10^{-09}$ | *HCP5 MICA* |
| 2,657 | 6 | 31,380,001 | 31,380,601 | 601 | 8 | $2.16\times10^{-11}$ | 0.001 | $1.77\times10^{-04}$ | $5.65\times10^{-03}$ | 1.21 | $1.49\times10^{-09}$ | *HCP5 MICA* |
| 2,657 | 6 | 31,390,874 | 31,391,175 | 302 | 6 | $1.20\times10^{-14}$ | 0.001 | $1.77\times10^{-04}$ | $5.65\times10^{-03}$ | 1.31 | $1.45\times10^{-11}$ | *HCP5* |
| 2,657 | 6 | 31,421,170 | 31,421,547 | 378 | 5 | $4.20\times10^{-20}$ | 0.001 | $1.77\times10^{-04}$ | $5.65\times10^{-03}$ | 1.33 | $1.04\times10^{-14}$ | *HCP5* |
| 2,657 | 6 | 31,444,174 | 31,445,396 | 1,223 | 8 | $2.83\times10^{-15}$ | 0.001 | $1.77\times10^{-04}$ | $5.65\times10^{-03}$ | 0.78 | $1.22\times10^{-17}$ | *HCP5* |
| 2,657 | 6 | 31,473,707 | 31,474,883 | 1,177 | 6 | $1.71\times10^{-10}$ | 0.001 | $1.77\times10^{-04}$ | $5.65\times10^{-03}$ | 1.28 | $2.08\times10^{-13}$ | *MICB* |
| 2,657 | 6 | 31,480,212 | 31,481,149 | 938 | 10 | $4.19\times10^{-15}$ | 0.001 | $1.77\times10^{-04}$ | $5.65\times10^{-03}$ | 1.25 | $1.03\times10^{-12}$ | |
| 2,657 | 6 | 31,483,892 | 31,485,240 | 1,349 | 6 | $7.72\times10^{-11}$ | 0.001 | $1.77\times10^{-04}$ | $5.65\times10^{-03}$ | 1.28 | $1.70\times10^{-13}$ | *XXbac-BPG16N22.5* |
| 2,657 | 6 | 31,554,382 | 31,554,382 | 1 | 1 | $1.53\times10^{-10}$ | 0.001 | $1.77\times10^{-04}$ | $5.65\times10^{-03}$ | 0.64 | $2.93\times10^{-11}$ | *LST1* |
| 2,657 | 6 | 31,560,910 | 31,561,913 | 1,004 | 5 | $4.03\times10^{-16}$ | 0.001 | $1.77\times10^{-04}$ | $5.65\times10^{-03}$ | 0.73 | $1.55\times10^{-17}$ | |
| 2,657 | 6 | 31,587,134 | 31,592,082 | 4,949 | 20 | $5.35\times10^{-35}$ | 0.001 | $1.77\times10^{-04}$ | $5.65\times10^{-03}$ | 0.70 | $7.44\times10^{-37}$ | *PRRC2A SNORA38* |
| 2,657 | 6 | 31,600,906 | 31,602,654 | 1,749 | 10 | $7.76\times10^{-14}$ | 0.001 | $1.77\times10^{-04}$ | $5.65\times10^{-03}$ | 0.73 | $3.74\times10^{-10}$ | *PRRC2A* |
| 2,657 | 6 | 31,606,813 | 31,606,977 | 165 | 3 | $5.87\times10^{-12}$ | 0.001 | $1.77\times10^{-04}$ | $5.65\times10^{-03}$ | 0.60 | $1.77\times10^{-13}$ | *BAG6* |
| 2,657 | 6 | 31,607,952 | 31,608,827 | 876 | 6 | $4.39\times10^{-13}$ | 0.001 | $1.77\times10^{-04}$ | $5.65\times10^{-03}$ | 0.71 | $1.92\times10^{-10}$ | *BAG6* |
| 2,657 | 6 | 31,633,482 | 31,633,552 | 71 | 3 | $3.59\times10^{-16}$ | 0.001 | $1.77\times10^{-04}$ | $5.65\times10^{-03}$ | 1.31 | $2.35\times10^{-14}$ | *CSNK2B GPANK1* |

| | | | | | | | | | | | | |
|---|---|---|---|---|---|---|---|---|---|---|---|---|
| 2,657 | 6 | 31,647,027 | 31,651,194 | 4,168 | 12 | $3.79\times10^{-14}$ | 0.001 | $1.77\times10^{-04}$ | $5.65\times10^{-03}$ | 1.24 | $1.71\times10^{-13}$ | *LY6G5C* |
| 2,657 | 6 | 31,656,565 | 31,656,565 | 1 | 1 | $6.35\times10^{-12}$ | 0.001 | $1.77\times10^{-04}$ | $5.65\times10^{-03}$ | 0.59 | $1.10\times10^{-13}$ | *ABHD16A* |
| 2,657 | 6 | 31,713,785 | 31,714,220 | 436 | 4 | $1.19\times10^{-25}$ | 0.001 | $1.77\times10^{-04}$ | $5.65\times10^{-03}$ | 1.37 | $6.25\times10^{-21}$ | *MSH5 MSH5-SAPCD1* |
| 2,657 | 6 | 31,729,958 | 31,730,568 | 611 | 3 | $3.30\times10^{-14}$ | 0.001 | $1.77\times10^{-04}$ | $5.65\times10^{-03}$ | 0.73 | $1.12\times10^{-12}$ | *MSH5 MSH5-SAPCD1* |
| 2,657 | 6 | 31,733,224 | 31,737,279 | 4,056 | 26 | $5.69\times10^{-21}$ | 0.001 | $1.77\times10^{-04}$ | $5.65\times10^{-03}$ | 0.76 | $1.15\times10^{-14}$ | *SAPCD1-AS1 VWA7* |
| 2,657 | 6 | 31,761,291 | 31,762,141 | 851 | 3 | $4.02\times10^{-17}$ | 0.001 | $1.77\times10^{-04}$ | $5.65\times10^{-03}$ | 0.56 | $1.06\times10^{-16}$ | *VARS* |
| 2,657 | 6 | 31,770,823 | 31,770,823 | 1 | 1 | $4.81\times10^{-17}$ | 0.001 | $1.77\times10^{-04}$ | $5.65\times10^{-03}$ | 0.62 | $1.10\times10^{-16}$ | *LSM2* |
| 2,657 | 6 | 31,788,479 | 31,792,174 | 3,696 | 9 | $4.35\times10^{-23}$ | 0.001 | $1.77\times10^{-04}$ | $5.65\times10^{-03}$ | 0.73 | $4.43\times10^{-16}$ | |
| 2,657 | 6 | 31,798,707 | 31,799,223 | 517 | 4 | $5.39\times10^{-15}$ | 0.001 | $1.77\times10^{-04}$ | $5.65\times10^{-03}$ | 0.73 | $1.11\times10^{-09}$ | |
| 2,657 | 6 | 31,799,353 | 31,800,434 | 1,082 | 5 | $7.47\times10^{-10}$ | 0.002 | $5.49\times10^{-04}$ | 7.27×10-03 | 1.21 | $1.05\times10^{-09}$ | |
| 2,657 | 6 | 31,807,148 | 31,807,652 | 505 | 7 | $1.22\times10^{-31}$ | 0.001 | $1.77\times10^{-04}$ | $5.65\times10^{-03}$ | 0.64 | $1.01\times10^{-27}$ | *C6orf48* |
| 2,657 | 6 | 31,844,593 | 31,847,180 | 2,588 | 15 | $1.83\times10^{-49}$ | 0.001 | $1.77\times10^{-04}$ | $5.65\times10^{-03}$ | 0.65 | $1.31\times10^{-56}$ | *SLC44A4* |
| 2,657 | 6 | 31,878,006 | 31,878,721 | 716 | 5 | $3.78\times10^{-80}$ | 0.001 | $1.77\times10^{-04}$ | $5.65\times10^{-03}$ | 0.53 | $1.23\times10^{-70}$ | *C2* |
| 2,657 | 6 | 32,006,004 | 32,006,453 | 450 | 4 | $6.80\times10^{-10}$ | 0.001 | $1.77\times10^{-04}$ | $5.65\times10^{-03}$ | 1.36 | $3.32\times10^{-08}$ | *CYP21A2* |
| 2,657 | 6 | 32,017,545 | 32,018,345 | 801 | 3 | $6.61\times10^{-54}$ | 0.001 | $1.77\times10^{-04}$ | $5.65\times10^{-03}$ | 0.56 | $6.61\times10^{-51}$ | *TNXB* |
| 2,657 | 6 | 32,070,602 | 32,070,838 | 237 | 3 | $4.18\times10^{-48}$ | 0.001 | $1.77\times10^{-04}$ | $5.65\times10^{-03}$ | 0.56 | $1.02\times10^{-42}$ | *ATF6B FKBPL* |
| 2,657 | 6 | 32,093,363 | 32,097,421 | 4,059 | 17 | $3.49\times10^{-54}$ | 0.001 | $1.77\times10^{-04}$ | $5.65\times10^{-03}$ | 0.59 | $1.14\times10^{-56}$ | *ATF6B FKBPL* |
| 2,657 | 6 | 32,188,713 | 32,188,853 | 141 | 4 | $2.46\times10^{-38}$ | 0.001 | $1.77\times10^{-04}$ | $5.65\times10^{-03}$ | 0.60 | $1.15\times10^{-37}$ | *NOTCH4* |
| 2,657 | 6 | 32,192,330 | 32,192,560 | 231 | 5 | $5.44\times10^{-10}$ | 0.001 | $1.77\times10^{-04}$ | $5.65\times10^{-03}$ | 0.77 | $5.89\times10^{-11}$ | |
| 2,657 | 6 | 32,196,249 | 32,196,389 | 141 | 3 | $3.33\times10^{-15}$ | 0.001 | $1.77\times10^{-04}$ | $5.65\times10^{-03}$ | 0.70 | $1.36\times10^{-13}$ | |
| 2,657 | 6 | 32,198,847 | 32,202,201 | 3,355 | 18 | $2.57\times10^{-16}$ | 0.001 | $1.77\times10^{-04}$ | $5.65\times10^{-03}$ | 1.23 | $1.06\times10^{-17}$ | |
| 2,657 | 6 | 32,210,714 | 32,211,077 | 364 | 4 | $1.32\times10^{-13}$ | 0.001 | $1.77\times10^{-04}$ | $5.65\times10^{-03}$ | 1.39 | $3.58\times10^{-16}$ | |
| 2,657 | 6 | 32,231,102 | 32,232,988 | 1,887 | 11 | $2.91\times10^{-19}$ | 0.001 | $1.77\times10^{-04}$ | $5.65\times10^{-03}$ | 0.672 | $4.18\times10^{-13}$ | *XXbac-BPG154L12.4* |
| 2,657 | 6 | 32,256,782 | 32,257,200 | 419 | 4 | $1.33\times10^{-11}$ | 0.001 | $1.77\times10^{-04}$ | $5.65\times10^{-03}$ | 1.34 | $1.01\times10^{-11}$ | *C6orf10* |
| 2,657 | 6 | 32,303,245 | 32,303,474 | 230 | 3 | $3.66\times10^{-16}$ | 0.001 | $1.77\times10^{-04}$ | $5.65\times10^{-03}$ | 1.38 | $1.19\times10^{-17}$ | *C6orf10* |

| | | | | | | | | | | | | |
|---|---|---|---|---|---|---|---|---|---|---|---|---|
| 2,657 | 6 | 32,306,922 | 32,308,729 | 1,808 | 8 | $3.62\times10^{-16}$ | 0.001 | $1.77\times10^{-04}$ | $5.65\times10^{-03}$ | 1.28 | $2.98\times10^{-22}$ | *C6orf10* |
| 2,657 | 6 | 32,350,328 | 32,350,622 | 295 | 4 | $7.88\times10^{-14}$ | 0.001 | $1.77\times10^{-04}$ | $5.65\times10^{-03}$ | 1.38 | $4.54\times10^{-16}$ | |
| 2,657 | 6 | 32,407,537 | 32,407,773 | 237 | 2 | $3.73\times10^{-12}$ | 0.001 | $1.77\times10^{-04}$ | $5.65\times10^{-03}$ | 0.66 | $1.02\times10^{-12}$ | *HLA-DRA* |
| 2,657 | 10 | 124,226,492 | 124,249,185 | 22,694 | 64 | $2.09\times10^{-84}$ | 0.001 | $1.77\times10^{-04}$ | $5.65\times10^{-03}$ | 0.62 | $2.96\times10^{-30}$ | *HTRA1* |
| 2,657 | 19 | 5,827,765 | 5,828,064 | 300 | 2 | $2.49\times10^{-11}$ | 0.001 | $1.77\times10^{-04}$ | $5.65\times10^{-03}$ | 0.74 | $1.04\times10^{-12}$ | *NRTN* |
| 2,657 | 19 | 5,832,268 | 5,832,773 | 506 | 3 | $1.55\times10^{-10}$ | 0.001 | $1.77\times10^{-04}$ | $5.65\times10^{-03}$ | 0.74 | $4.84\times10^{-12}$ | *FUT6* |
| 2,657 | 19 | 5,835,677 | 5,835,677 | 1 | 1 | $8.06\times10^{-12}$ | 0.001 | $1.77\times10^{-04}$ | $5.65\times10^{-03}$ | 0.74 | $1.22\times10^{-13}$ | *FUT6* |
| 2,657 | 22 | 33,074,351 | 33,091,173 | 16,823 | 11 | $1.02\times10^{-16}$ | 0.001 | $1.77\times10^{-04}$ | $5.65\times10^{-03}$ | 0.69 | $5.71\times10^{-11}$ | *SYN3* |
| 2,657 | 22 | 33,097,419 | 33,101,049 | 3,631 | 11 | $1.78\times10^{-16}$ | 0.001 | $1.77\times10^{-04}$ | $5.65\times10^{-03}$ | 0.70 | $1.27\times10^{-11}$ | *SYN3* |

**Table S10: Genes overlapping with significant bins were detected by GECS and verified by SKAT in the AAMD data set. Mapped gene ID refers to availability of the ID in gene ontology.**

| Chr. | Locus name | Mapped gene ID | Previously reported genes |
|---|---|---|---|
| 6 | *ABHD16A* | TRUE | NO |
| 6 | *AL671883.1* | FALSE | NO |
| 10 | *ARMS2* | TRUE | YES |
| 6 | *ATF6B* | TRUE | YES |
| 6 | *BAG6* | TRUE | NO |
| 6 | *BTNL2* | TRUE | NO |
| 6 | *C1orf53* | TRUE | NO |
| 6 | *C2* | TRUE | YES |
| 19 | *C3* | TRUE | YES |
| 6 | *C6orf10* | TRUE | NO |
| 6 | *C6orf48* | TRUE | NO |
| 5 | *C9* | TRUE | YES |
| 4 | *CFI* | TRUE | YES |
| 6 | *CSNK2B* | TRUE | NO |
| 6 | *CYP21A2* | TRUE | NO |
| 6 | *DXO* | TRUE | YES |
| 6 | *FKBPL* | TRUE | NO |
| 19 | *FUT6* | TRUE | YES |
| 5 | *FYB* | TRUE | NO |
| 6 | *GPANK1* | TRUE | NO |
| 6 | *HCG22* | TRUE | NO |
| 6 | *HCP5* | TRUE | NO |
| 6 | *HLA-B* | TRUE | NO |
| 6 | *HLA-DRA* | TRUE | NO |
| 10 | *HTRA1* | TRUE | YES |
| 6 | *LSM2* | TRUE | NO |
| 6 | *LST1* | TRUE | NO |
| 6 | *LY6G5C* | TRUE | NO |
| 6 | *MICA* | TRUE | NO |
| 6 | *MICB* | TRUE | NO |
| 6 | *MIR3941* | FALSE | NO |
| 6 | *MSH5* | TRUE | NO |
| 6 | *MSH5-SAPCD1* | FALSE | NO |
| 1 | *NEK7* | TRUE | NO |
| 6 | *NFKBIL1* | TRUE | NO |
| 6 | *NOTCH4* | TRUE | YES |
| 19 | *NRTN* | TRUE | YES |
| 10 | *PLEKHA1* | TRUE | YES |
| 6 | *PRRC2A* | TRUE | NO |

| | | | |
|---|---|---|---|
| 6 | *SKIV2L* | TRUE | YES |
| 6 | *SLC44A4* | TRUE | NO |
| 6 | *SNORA38* | FALSE | NO |
| 22 | *SYN3* | TRUE | YES |
| 6 | *TNXB* | TRUE | YES |
| 6 | *TRIM31* | TRUE | NO |
| 6 | *TRIM31-AS1* | FALSE | NO |
| 6 | *VARS* | TRUE | NO |
| 6 | *VWA7* | TRUE | NO |
| 6 | *XXbac-BPG154L12.4* | FALSE | NO |
| 6 | *XXbac-BPG16N22.5* | FALSE | NO |
| 6 | *XXbac-BPG181B23.4* | FALSE | NO |
| 6 | *XXbac-BPG32J3.20* | FALSE | NO |

**Table S11: A list of gene ontology* results for GECS findings in AAMD data set (significant results in grey).**

| GO biological process | Homo sapiens - REFLIST (20996) | upload_1 (44) | upload_1 (expected) | upload_1 (over/under) | upload_1 (fold Enrichment) | upload_1 (p-value) |
|---|---|---|---|---|---|---|
| positive regulation of immune response (GO:0050778) | 778 | 12 | 1.63 | + | 7.36 | $4,40\times10^{-04}$ |
| regulation of immune response (GO:0050776) | 1,056 | 13 | 2.21 | + | 5.87 | $1,45\times10^{-03}$ |
| activation of immune response (GO:0002253) | 560 | 10 | 1.17 | + | 8.52 | $1,86\times10^{-03}$ |
| regulation of immune system process (GO:0002682) | 1,557 | 15 | 3.26 | + | 4.60 | $2,98\times10^{-03}$ |
| positive regulation of immune system process (GO:0002684) | 1,064 | 12 | 2.23 | + | 5.38 | $1,21\times10^{-02}$ |
| regulation of immune effector process (GO:0002697) | 451 | 8 | .95 | + | 8.46 | $3,75\times10^{-02}$ |
| lymphocyte mediated immunity (GO:0002449) | 258 | 6 | 0.54 | + | 11.10 | $1,52\times10^{-01}$ |
| innate immune response (GO:0045087) | 741 | 9 | 1.55 | + | 5.80 | $1,79\times10^{-01}$ |
| adaptive immune response based on somatic recombination of immune receptors built from immunoglobulin superfamily domains (GO:0002460) | 269 | 6 | 0.56 | + | 10.64 | $1,92\times10^{-01}$ |
| negative regulation of immune system process (GO:0002683) | 429 | 7 | 0.90 | + | 7.79 | $2,70\times10^{-01}$ |
| cytolysis (GO:0019835) | 27 | 3 | 0.06 | + | 53.02 | $2,91\times10^{-01}$ |
| immune system process (GO:0002376) | 2,687 | 16 | 5.63 | + | 2.84 | $5,46\times10^{-01}$ |
| positive regulation of lymphocyte mediated immunity (GO:0002708) | 104 | 4 | 0.22 | + | 18.35 | $6,64\times10^{-0}1$ |
| regulation of complement activation (GO:0030449) | 113 | 4 | 0.24 | + | 16.89 | $9,06\times10^{-01}$ |

* Analysis type: PANTHER overrepresentation Test (Released 20181113). Annotation version and release date: GO Ontology, database released 2018-12-01. Analyzed list: upload_1 (Homo sapiens). Reference list: Homo sapiens (all genes in database). Test type: FISHER. Correction: BONFERRONI. Bonferroni count: 8723.

**Table S12: Previously reported rare-variant association signals in AAMD which were detected as significant by GECS (nominal p-values shown, see Methods for more information).**

| Chr. | Locus | rs-ID | Bin position (hg19) | | Number of RV's | p-value |
|---|---|---|---|---|---|---|
| 1 | *CFH* | rs121913059 | 196,734,777 | 19,6748,844 | 22 | $4.68\times10^{-25}$ |
| 1 | *KCNT2 - CFH* | rs148553336 | 196,642,858 | 196,644,117 | 11 | $2.94\times10^{-35}$ |
| 1 | *KCNT2* | rs187328863 | 196,383,386 | 96,412,490 | 88 | $8.77\times10^{-29}$ |
| 1 | *CFH* | rs35292876 | 196,730,754 | 196,738,102 | 19 | $8.89\times10^{-31}$ |
| 1 | *CFH* | rs191281603 | 196,958,365 | 196,961,953 | 37 | $1.94\times10^{-31}$ |
| 3 | *LOC105374007* | rs140647181 | | | | |
| 4 | *CFI* | rs141853578 | | | | |
| 5 | *C9* | rs62358361 | 39,326,270 | 39,327,888 | 12 | $4.43\times10^{-10}$ |
| 5 | *PRLR - SPEF2* | rs114092250 | 35,494,324 | 35,494,448 | 3 | $3.55\times10^{-07}$ |
| 6 | *TNXB* | rs12153855 | 32,105,360 | 32,109,828 | 14 | $5.86\times10^{-19}$ |
| 6 | *STK19* | rs2746394 | 31,946,792 | 31,947,261 | 5 | $6.12\times10^{-14}$ |
| 6 | *C2-AS1, C2* | rs9380272 | 31,938,120 | 31,938,515 | 7 | $7.72\times10^{-15}$ |
| 6 | *STK19* | rs181705462 | 31,933,248 | 31,980,000 | 125 | $1.99\times10^{-12}$ |
| 12 | *ACAD10* | rs61941274 | | | | |
| 15 | *HERC1* | rs74320127 | | | | |
| 19 | *C3* | rs147859257 | | | | |

**Table S13: Huge bins (number of rare variants ≥ 1,000) that were detected as significant by GECS in the AAMD data set.**

| NCT | Chr. | Bin position | | Number of rare variants | p_value |
|---|---|---|---|---|---|
| 542 | 1 | 194,548,913 | 197,626,814 | 6,351 | $8.27\times10^{-10}$ |
| 542 | 1 | 196,133,060 | 198,510,045 | 6,168 | $8.61\times10^{-10}$ |
| 542 | 1 | 196,150,322 | 198,524,866 | 6,189 | $6.33\times10^{-10}$ |
| 542 | 10 | 123,863,539 | 124,449,418 | 1,192 | $9.45\times10^{-10}$ |
| 542 | 10 | 123,878,617 | 124,449,418 | 1,185 | $2.88\times10^{-10}$ |
| 542 | 19 | 281,468 | 2,557,330 | 1,082 | $3.91\times10^{-10}$ |
| 542 | 19 | 281,468 | 2,558,709 | 1,083 | $5.27\times10^{-10}$ |
| 1,611 | 1 | 196,311,284 | 197,201,223 | 3,729 | $6.44\times10^{-10}$ |
| 1,611 | 1 | 196,311,578 | 197,201,223 | 3,728 | $6.85\times10^{-10}$ |
| 1,611 | 1 | 196,380,240 | 197,236,152 | 3,730 | $4.37\times10^{-10}$ |
| 1,611 | 10 | 123,988,786 | 124,425,237 | 1,420 | $3.84\times10^{-10}$ |
| 1,611 | 10 | 123,993,600 | 124,435,927 | 1,444 | $1.77\times10^{-10}$ |
| 1,611 | 17 | 78,865,313 | 80,206,871 | 1,785 | $7.43\times10^{-10}$ |
| 1,611 | 17 | 78,982,299 | 80,369,949 | 1,781 | $8.79\times10^{-10}$ |
| 1,611 | 17 | 78,991,475 | 80,372,759 | 1,781 | $7.50\times10^{-10}$ |
| 1,611 | 19 | 1,116,932 | 2,481,236 | 1,205 | $9.09\times10^{-13}$ |
| 1,611 | 19 | 1,116,932 | 2,484,530 | 1,207 | $5.18\times10^{-13}$ |
| 2,657 | 1 | 196,681,376 | 197,026,151 | 1,567 | $3.55\times10^{-11}$ |
| 2,657 | 1 | 196,693,159 | 197,037,552 | 1,546 | $3.39\times10^{-11}$ |
| 2,657 | 1 | 196,693,761 | 197,037,552 | 1,544 | $2.63\times10^{-11}$ |
| 2,657 | 19 | 1,140,069 | 2,301,128 | 1,275 | $3.02\times10^{-11}$ |
| 2,657 | 19 | 1,140,069 | 2,305,170 | 1,280 | $2.88\times10^{-11}$ |
| 2,657 | 19 | 1,140,069 | 2,308,505 | 1,283 | $2.47\times10^{-11}$ |

**Table S14: Counts of significant signals from GECS and SMA in the AAMD data set.**

| Chr. | SMA | GECS | | |
|---|---|---|---|---|
| | | $MAF_T$ = 0.01 | $MAF_T$ = 0.03 | $MAF_T$ = 0.05 |
| 1 | 2,120 | 170,197 | 143,688 | 58,491 |
| 2 | 0 | 0 | 0 | 0 |
| 3 | 32 | 0 | 15 | 15 |
| 4 | 33 | 6 | 6 | 6 |
| 5 | 3 | 2 | 8 | 8 |
| 6 | 1,536 | 0 | 53,684 | 24,532 |
| 7 | 1 | 0 | 0 | 0 |
| 8 | 0 | 0 | 0 | 0 |
| 9 | 45 | 0 | 0 | 2 |
| 10 | 1,093 | 6,960 | 72,537 | 38,329 |
| 11 | 2 | 0 | 0 | 925 |
| 12 | 9 | 0 | 0 | 0 |
| 13 | 3 | 0 | 0 | 0 |
| 14 | 0 | 0 | 0 | 0 |
| 15 | 19 | 0 | 0 | 0 |
| 16 | 62 | 0 | 0 | 0 |
| 17 | 4 | 1 | 19,702 | 1,003 |
| 18 | 0 | 0 | 0 | 0 |
| 19 | 514 | 9,582 | 43,154 | 59,800 |
| 20 | 49 | 0 | 2 | 20 |
| 21 | 0 | 0 | 456 | 0 |
| 22 | 44 | 0 | 0 | 354 |
| **SUM** | **5,590** | **186,748** | **333,252** | **183,485** |

**Table S15: Bins with the locally most significant association signals in SCZD data set, detected by GECS and verified by SKAT.**

Each bin is the most significant signal in the block of all overlapping significant bins detected by GECS. These bins are verified by SKAT, adjusted for sex, age, 10 principal components, and common variants in physical proximity, if available (p'-values). For verification with SKAT, we set the threshold at $2\times10^{-6}$.

| NCT | Chr. | Bin position | | Bin size | Number of rare variants | p-value | Corrected p-value (p_corr) | Wilson lower CI 95% | Wilson upper CI 95% | OR | p'-value | Overlapping or close genes (+/-10kb) |
|---|---|---|---|---|---|---|---|---|---|---|---|---|
| 214 | 2 | 38,903,107 | 38,956,836 | 53,73 | 3 | $2.61\times10^{-10}$ | 0.001 | $1.77\times10^{-04}$ | $5.65\times10^{-03}$ | 0.20 | $2.97\times10^{-09}$ | *GALM* |
| 214 | 2 | 61,719,303 | 61,719,303 | 1 | 1 | $1.07\times10^{-11}$ | 0.001 | $1.77\times10^{-04}$ | $5.65\times10^{-03}$ | 0.32 | $8.05\times10^{-13}$ | *XPO1* |
| 214 | 3 | 187,444,543 | 187,444,543 | 1 | 1 | $1.07\times10^{-12}$ | 0.001 | $1.77\times10^{-04}$ | $5.65\times10^{-03}$ | 0.18 | $1.11\times10^{-14}$ | *BCL6* |
| 214 | 4 | 47,887,513 | 47,887,513 | 1 | 1 | $1.26\times10^{-08}$ | 0.017 | $1.77\times10^{-04}$ | $2.71\times10^{-02}$ | 0.39 | $2.42\times10^{-09}$ | *NFXL1* |
| 214 | 5 | 178,392,653 | 178,392,792 | 140 | 2 | $1.02\times10^{-08}$ | 0.014 | $8.37\times10^{-03}$ | $2.34\times10^{-02}$ | 0.19 | $3.63\times10^{-09}$ | *ZNF454* |
| 214 | 6 | 150,290,454 | 150,291,207 | 754 | 2 | $2.70\times10^{-13}$ | 0.001 | $1.77\times10^{-04}$ | $5.65\times10^{-03}$ | 0.27 | $1.90\times10^{-14}$ | *ULBP1* |
| 214 | 7 | 129,680,877 | 129,680,877 | 1 | 1 | $1.38\times10^{-08}$ | 0.019 | $1.22\times10^{-02}$ | $2.95\times10^{-02}$ | 0.13 | $1.66\times10^{-09}$ | *ZC3HC1* |
| 214 | 11 | 57,100,475 | 57,100,642 | 168 | 2 | $1.56\times10^{-12}$ | 0.001 | $1.77\times10^{-04}$ | $5.65\times10^{-03}$ | 0.04 | $1.65\times10^{-12}$ | *SSRP1* |
| 214 | 11 | 82,642,868 | 82,642,868 | 1 | 1 | $2.23\times10^{-09}$ | 0.003 | $1.02\times10^{-03}$ | $8.79\times10^{-03}$ | 1.01 | $1.54\times10^{-10}$ | *PRCP* |
| 214 | 12 | 53,455,228 | 53,455,228 | 1 | 1 | $3.63\times10^{-10}$ | 0.001 | $1.77\times10^{-04}$ | $5.65\times10^{-03}$ | 0.31 | $1.35\times10^{-10}$ | *TENC1* |
| 214 | 12 | 57,398,026 | 57,398,026 | 1 | 1 | $1.40\times10^{-11}$ | 0.001 | $1.77\times10^{-04}$ | $5.65\times10^{-03}$ | 0.20 | $2.17\times10^{-13}$ | *ZBTB39* |
| 214 | 13 | 31,897,996 | 31,897,996 | 1 | 1 | $9.72\times10^{-10}$ | 0.002 | $5.49\times10^{-04}$ | $7.27\times10^{-03}$ | 0.10 | $5.47\times10^{-12}$ | *B3GALTL* |
| 214 | 14 | 60,611,664 | 60,611,664 | 1 | 1 | $1.41\times10^{-08}$ | 0.021 | $1.38\times10^{-02}$ | $3.19\times10^{-02}$ | 0.17 | $6.09\times10^{-10}$ | *DHRS7 PCNXL4* |
| 214 | 14 | 89,078,090 | 89,088,986 | 10,897 | 2 | $1.14\times10^{-09}$ | 0.002 | $5.49\times10^{-04}$ | $7.27\times10^{-03}$ | 0.14 | $9.19\times10^{-11}$ | *EML5* |
| 214 | 17 | 49,239,143 | 49,239,143 | 1 | 1 | $2.72\times10^{-09}$ | 0.003 | $1.02\times10^{-03}$ | $8.79\times10^{-03}$ | 0.29 | $5.58\times10^{-11}$ | *NME1 NME1-NME2 NME2* |
| 214 | 22 | 17,687,954 | 17,688,129 | 176 | 2 | $3.79\times10^{-13}$ | 0.001 | $1.77\times10^{-04}$ | $5.65\times10^{-03}$ | 0.27 | $6.77\times10^{-14}$ | *CECR1* |
| 636 | 1 | 24,447,832 | 24,447,843 | 12 | 2 | $1.28\times10^{-14}$ | 0.001 | $1.77\times10^{-04}$ | $5.65\times10^{-03}$ | 0.38 | $5.86\times10^{-16}$ | *IL22RA1* |
| 636 | 3 | 138,402,579 | 138,402,579 | 1 | 1 | $1.42\times10^{-21}$ | 0.001 | $1.77\times10^{-04}$ | $5.65\times10^{-03}$ | 0.30 | $8.36\times10^{-23}$ | *PIK3CB* |
| 636 | 4 | 103,720,097 | 103,806,388 | 86,292 | 6 | $3.19\times10^{-17}$ | 0.001 | $1.77\times10^{-04}$ | $5.65\times10^{-03}$ | 0.35 | $4.07\times10^{-18}$ | *CISD2 RNU7-151P* |

| | | | | | | | | | | | | |
|---|---|---|---|---|---|---|---|---|---|---|---|---|
| | | | | | | | | | | | | *RP11-10L12.4 SLC9B1 snoU13 UBE2D3* |
| 636 | 5 | 32,415,110 | 32,415,216 | 107 | 2 | $2.12\times10^{-17}$ | 0.001 | $1.77\times10^{-04}$ | $5.65\times10^{-03}$ | 0.24 | $3.52\times10^{-18}$ | *ZFR* |
| 636 | 5 | 54,591,272 | 54,591,272 | 1 | 1 | $3.95\times10^{-12}$ | 0.001 | $1.77\times10^{-04}$ | $5.65\times10^{-03}$ | 0.38 | $1.20\times10^{-14}$ | *DHX29* |
| 636 | 7 | 128,413,777 | 128,413,777 | 1 | 1 | $1.96\times10^{-15}$ | 0.001 | $1.77\times10^{-04}$ | $5.65\times10^{-03}$ | 0.42 | $7.85\times10^{-17}$ | *OPN1SW* |
| 636 | 12 | 53,069,392 | 53,069,392 | 1 | 1 | $1.19\times10^{-09}$ | 0.003 | $1.02\times10^{-03}$ | $8.79\times10^{-03}$ | 0.58 | $8.07\times10^{-10}$ | *KRT1* |
| 636 | 14 | 67,848,325 | 67,848,325 | 1 | 1 | $3.67\times10^{-13}$ | 0.001 | $1.77\times10^{-04}$ | $5.65\times10^{-03}$ | 0.35 | $4.27\times10^{-12}$ | *EIF2S1* |
| 636 | 15 | 73,044,829 | 73,044,833 | 5 | 2 | $1.17\times10^{-19}$ | 0.001 | $1.77\times10^{-04}$ | $5.65\times10^{-03}$ | 0.43 | $4.30\times10^{-20}$ | *ADPGK* |
| 636 | 19 | 8,999,386 | 9,028,410 | 29,025 | 62 | $2.59\times10^{-09}$ | 0.004 | $1.56\times10^{-03}$ | $1.02\times10^{-02}$ | 1.29 | $3.10\times10^{-10}$ | *MUC16* |
| 1,049 | 1 | 202,724,559 | 202,729,678 | 5,12 | 3 | $3.06\times10^{-13}$ | 0.001 | $1.77\times10^{-04}$ | $5.65\times10^{-03}$ | 0.42 | $3.77\times10^{-13}$ | *KDM5B* |
| 1,049 | 9 | 33,796,672 | 33,798,630 | 1,959 | 20 | $5.07\times10^{-10}$ | 0.003 | $1.02\times10^{-03}$ | $8.79\times10^{-03}$ | 1.37 | $3.89\times10^{-11}$ | *PRSS3* |

**Table S16: Genes overlapping with significant bins were detected by GECS and verified by SKAT in the SCZD data set. Unmapped gene ID refers to availability of the ID in gene ontology.**

| Chr. | Locus name | Unmapped gene ID | Previously reported (Purcell et al 2014, Genovese et al 2016) |
|---|---|---|---|
| 1 | *IL22RA1* | NO | YES |
| 1 | *KDM5B* | NO | YES |
| 1 | *KLHL12* | NO | YES |
| 1 | *RABIF* | NO | NO |
| 1 | *HNRNPA1P59* | YES | NO |
| 1 | *PCAT6* | YES | NO |
| 1 | *RP11-480I12.10* | YES | NO |
| 1 | *RP11-480I12.2* | YES | NO |
| 1 | *RP11-480I12.4* | YES | NO |
| 1 | *RP11-480I12.5* | YES | NO |
| 1 | *RP11-480I12.7* | YES | NO |
| 1 | *RP11-480I12.9* | YES | NO |
| 1 | *SLC25A39P1* | YES | NO |
| 1 | *Y_RNA* | YES | NO |
| 2 | *XPO1* | NO | NO |
| 2 | *GALM* | NO | YES |
| 3 | *BCL6* | NO | YES |
| 3 | *PIK3CB* | NO | YES |
| 3 | *RP11-211G3* | YES | NO |
| 4 | *NFXL1* | NO | YES |
| 4 | *CISD2* | NO | No |
| 4 | *SLC9B1* | NO | YES |
| 4 | *UBE2D3* | NO | No |
| 4 | *RNU7-151P* | YES | NO |
| 4 | *RP11-10L12* | YES | NO |
| 4 | *snoU13* | YES | NO |
| 5 | *ZNF454* | NO | YES |
| 5 | *DHX29* | NO | YES |
| 5 | *ZFR* | NO | NO |
| 6 | *ULBP1* | NO | NO |
| 7 | *ZC3HC1* | NO | YES |
| 7 | *OPN1SW* | NO | YES |
| 9 | *PRSS3* | NO | NO |

| | | | |
|---|---|---|---|
| 9 | *RP11-133O22.6* | YES | NO |
| 11 | *SSRP1* | NO | NO |
| 11 | *PRCP* | NO | YES |
| 11 | *C11orf82* | NO | YES |
| 12 | *ZBTB39* | NO | YES |
| 12 | *KRT1* | NO | YES |
| 12 | *TENC1* | YES | YES |
| 13 | *B3GALTL* | NO | YES |
| 14 | *DHRS7* | NO | NO |
| 14 | *EML5* | NO | YES |
| 14 | *ZC3H14* | NO | NO |
| 14 | *PCNXL4* | NO | YES |
| 14 | *EIF2S1* | NO | NO |
| 15 | *ADPGK* | NO | NO |
| 17 | *NME1* | NO | YES |
| 17 | *NME2* | NO | NO |
| 19 | *MUC16* | NO | YES |
| 22 | *CECR1* | NO | YES |

**Table S17: Largest bins (number of rare variants ≥ 50) that were detected as significant by GECS in the SCZD data set.**

| NCT | Chr. | Bin position | | Number of rare variants | p-value |
|---|---|---|---|---|---|
| 214 | 1 | 16,525,790 | 16,918,465 | 77 | $6.54\times10^{-09}$ |
| 214 | 1 | 16,528,955 | 16,918,473 | 77 | $2.93\times10^{-09}$ |
| 214 | 1 | 16,528,955 | 17,030,399 | 80 | $3.33\times10^{-09}$ |
| 214 | 1 | 16,531,261 | 16,918,523 | 77 | $3.92\times10^{-09}$ |
| 636 | 7 | 128,049,496 | 128,471,084 | 56 | $6.06\times10^{-09}$ |
| 636 | 7 | 128,049,590 | 128,471,084 | 54 | $4.70\times10^{-09}$ |
| 636 | 16 | 89,168,971 | 89,347,730 | 89 | $1.92\times10^{-08}$ |
| 636 | 19 | 8,994,484 | 9,026,247 | 66 | $1.03\times10^{-08}$ |
| 636 | 19 | 8,997,119 | 9,026,247 | 65 | $1.25\times10^{-08}$ |
| 636 | 19 | 8,997,472 | 9,026,247 | 64 | $8.74\times10^{-09}$ |
| 636 | 19 | 8,998,663 | 9,026,247 | 63 | $7.79\times10^{-09}$ |

**Table S18: Performance of GECS in simulated data sets under the null model.**

| Method | $N_T^C$ | $MAF_T$ | Number of variants (millions) | Sample size | Average time* (h) | Average memory in gb | Average number of all bins* (millions) | Average number of distinct bins* (millions) | Average reduction rate |
|---|---|---|---|---|---|---|---|---|---|
| **GECS** | 19 | 0.01 | 3 | 1,000 | 6 | 3 | 295,000 | 276 | 99.91% |
| | 59 | 0.03 | 5 | 1,000 | 6 | 4 | 795,000 | 285 | 99.96% |
| | 97 | 0.05 | 6 | 1,000 | 6 | 4 | 1,064,000 | 259 | 99.98% |
| **SMA** | | | 12 | 1,000 | 0.2 | 7 | | | |
| **GECS** | 99 | 0.01 | 4 | 5,000 | 42 | 11 | 386,000 | 730 | 99.81% |
| | 295 | 0.03 | 6 | 5,000 | 38 | 13 | 902,000 | 667 | 99.93% |
| | 487 | 0.05 | 6 | 5,000 | 34 | 13 | 1,186,000 | 599 | 99.95% |
| **SMA** | | | 12 | 5,000 | 3 | 19 | | | |
| **GECS** | 199 | 0.01 | 4 | 10,000 | 129 | 20 | 411,000 | 1,083 | 99.74% |
| | 591 | 0.03 | 6 | 10,000 | 110 | 22 | 1,001,000 | 931 | 99.91% |
| | 975 | 0.05 | 7 | 10,000 | 98 | 24 | 1,205,000 | 890 | 99.93% |
| **SMA** | | | 12 | 10,000 | 4 | 34 | | | |
| **GECS** | 398 | 0.01 | 4 | 20,000 | 452 | 40 | 413,000 | 1,504 | 99.64% |
| | 1,181 | 0.03 | 6 | 20,000 | 380 | 34 | 1,070,000 | 1,360 | 99.86% |
| | 1,950 | 0.05 | 7 | 20,000 | 340 | 29 | 1,213,000 | 1,230 | 99.88% |
| **SMA** | | | 13 | 20,000 | 12 | 63 | | | |
| **GECS**** | 398 | 0.01 | 0.05-0.3 | 20,000 | 6-100 | >1-3 | 1,339-57,010 | 20-130 | 98.44%-99.77% |
| | 1,181 | 0.03 | 0.07-0.5 | 20,000 | 5-85 | >1-3 | 3,468-147,655 | 50-325 | 98.67%-99.91% |
| | 1,950 | 0.05 | 0.09-0.6 | 20,000 | 4-75 | >1-3 | 3,953-168,327 | 56-360 | 98.73%-99.93% |

* Average across 1,000 simulations.
** GECS was performed in parallel for each chromosome independently; min and max values among all chromosomes were considered.

**Table S19: Performance of GECS in the imputed whole-genome AAMD data set and in the whole-exome SCZD data set.**

| | Method | $N_T^C$ | $MAF_T$ | Number of variants on chromosomes | Sample size | Time (h) | Memory in gb | Number of all bins (millions) | Number of distinct bins (millions) |
|---|---|---|---|---|---|---|---|---|---|
| **AAMD** | **GECS** | 542 | 0.01 | 34-267k | 27,259 | 3 | >1-4 | 600-38,000 | 5-35 |
| | | 1,611 | 0.03 | 54-406k | | 4 | >1-4 | 1,000-82,000 | 6-40 |
| | | 2,657 | 0.05 | 64-470k | | 3 | >1-5 | 2,000-111,000 | 5-36 |
| | **SMA** | | | 927k | | 14 | 72 | | |
| **SCZD** | **GECS** | 214 | 0.01 | 219k | 10,763 | 14 | >1 | 1,300 | 116 |
| | | 636 | 0.03 | 241k | | 10 | 1 | 1,600 | 65 |
| | | 1,049 | 0.05 | 251k | | 8 | 1 | 1,700 | 49 |
| | **SMA** | | | 308k | | 6 | 2 | | |

**Table S20: Plots visualizing the results of the power study for different parameters.**

| | Figure | | | |
|---|---|---|---|---|
| | Rare diseases, K=0.01 | | Common diseases, K=0.1 | |
| **Sample size** | **PNV=0.1** | **PNV=0.3** | **PNV=0.1** | **PNV=0.3** |
| 1,000 | Figure S9 | Figure 2 | Figure S11 | Figure S6 |
| 10,000 | Figure S10 | Figure 3 | Figure S12 | Figure S7 |
| 20,000 | - | Figure S5 | - | Figure S8 |

### Figures:

**Figure S1: Combined carrier matrix B and inductive computation of bins.** The arrays $v_1, v_2, \ldots, v_n$ are binary, each of length *N*, where *N* denotes number of samples and the elements $v_{i,l}$ indicate the carrier status of the *l*-th individual (1=carrier, 0=non-carrier) at the *i-th* variant.

$$B = \begin{pmatrix} v_1 & (v_1 \vee v_2) & (v_1 \vee v_2 \vee v_3) & \cdots & \cdots & (v_1 \vee \ldots \vee v_{n-1}) & (v_1 \vee \ldots \vee v_n) \\ 0 & v_2 & (v_2 \vee v_3) & & & \cdots & (v_2 \vee \ldots \vee v_n) \\ \vdots & 0 & v_3 & & & & \vdots \\ \vdots & & & \ddots & & & \vdots \\ & & & & \ddots & & \\ \vdots & & & & 0 & v_{n-1} & (v_{n-1} \vee v_n) \\ 0 & \cdots & & & \cdots & 0 & v_n \end{pmatrix}$$

Inductive computation of bins:

$$B_{i,i} = v_i$$

$$B_{i,j} = B_{i,j-1} \vee v_j$$

$$B = \begin{pmatrix} v_1 & B_{1,2} & B_{1,3} & \cdots & \cdots & B_{1,n-1} & B_{1,n} \\ 0 & v_2 & B_{2,3} & & & \cdots & B_{2,n} \\ \vdots & 0 & v_3 & & & & \vdots \\ \vdots & & & \ddots & & & \vdots \\ & & & & \ddots & & \\ \vdots & & & & 0 & v_{n-1} & B_{n-1,n} \\ 0 & \cdots & & & \cdots & 0 & v_n \end{pmatrix}$$

**Figure S2: In-silico post-GECS analysis**

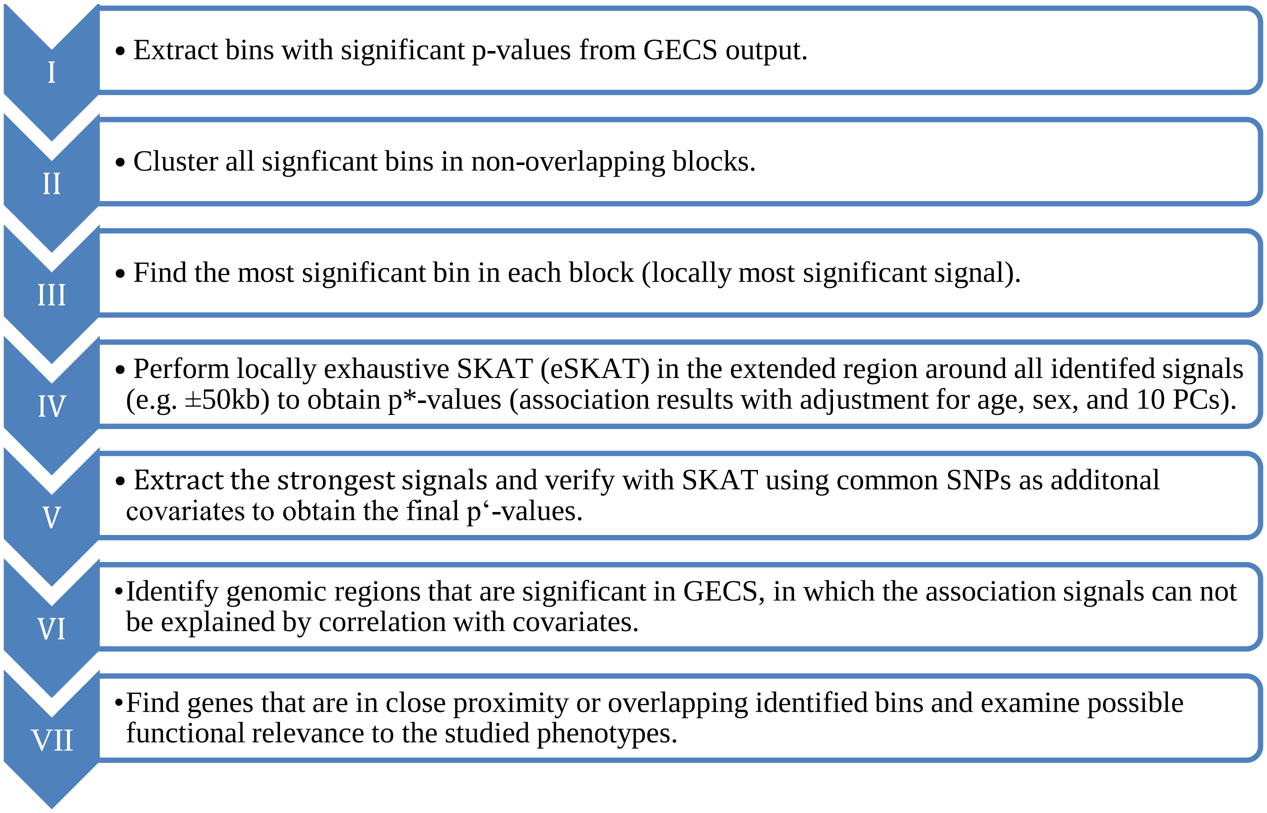

**Figure S3: P-value densities obtained from GECS in 1,000 simulated studies with 1,000 samples each, compared to corresponding results from single-marker analyses (SMA).**

$MAF_T$: minor allele frequency threshold.

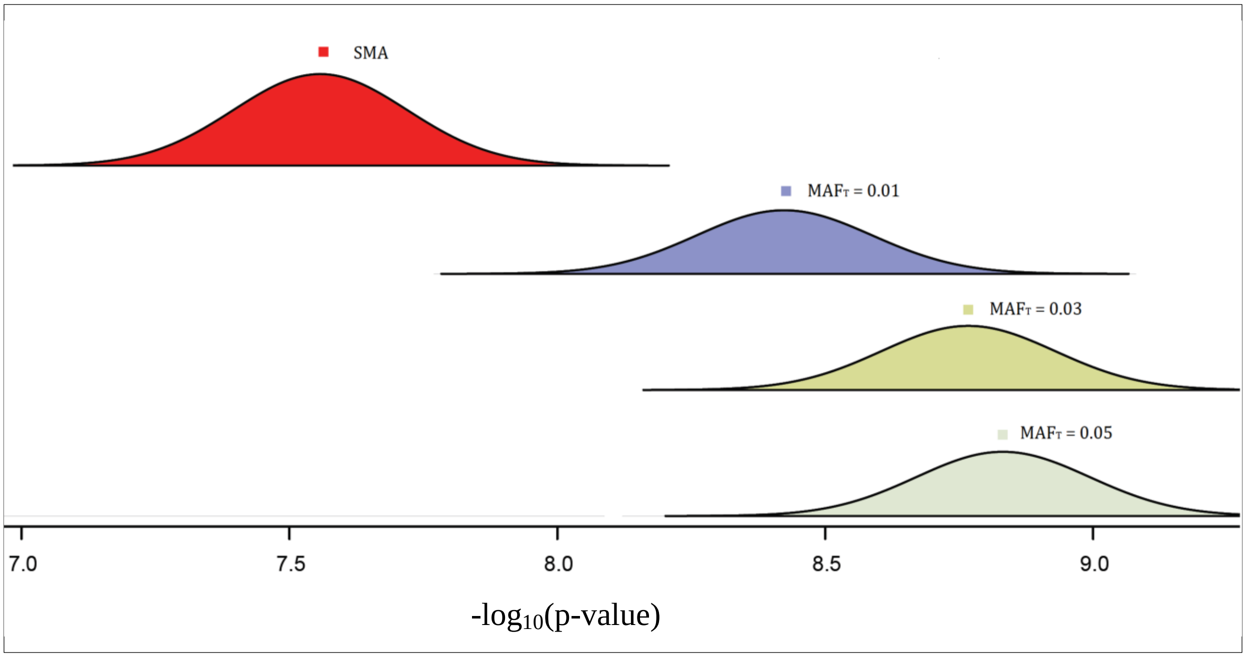

**Figure S4: P-value densities obtained from GECS in 1000 simulated studies with 5,000, 10,000, and 20,000 samples each, compared to corresponding results from single-marker analyses (SMA).** Distributions always shifted to the right with increasing sample sizes.

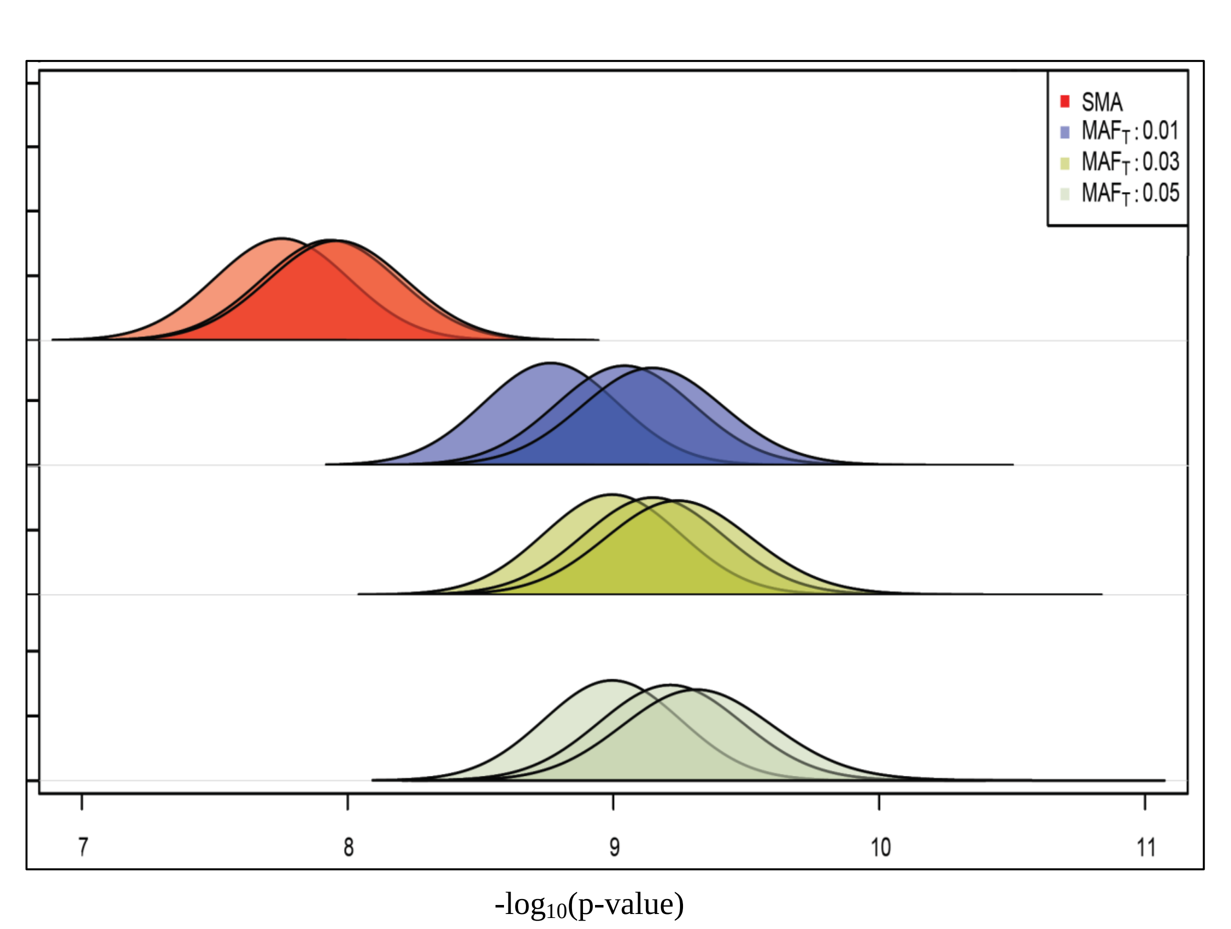

**Figure S5: Comparative power analysis for a rare disease (prevalence K=0.01) and large sample size (N=20,000).** Results are given for studies with proportion of neutral rare variants (PNV) = 0.3, different simulated window sizes (x-axis), and different proportions of detrimental rare variants (PDV) (y-axis). Black lines: GECS; red lines: SMA. In each grid cell, the power is presented on the y-axis and OR intervals on the x-axis. For an overview see table S20.

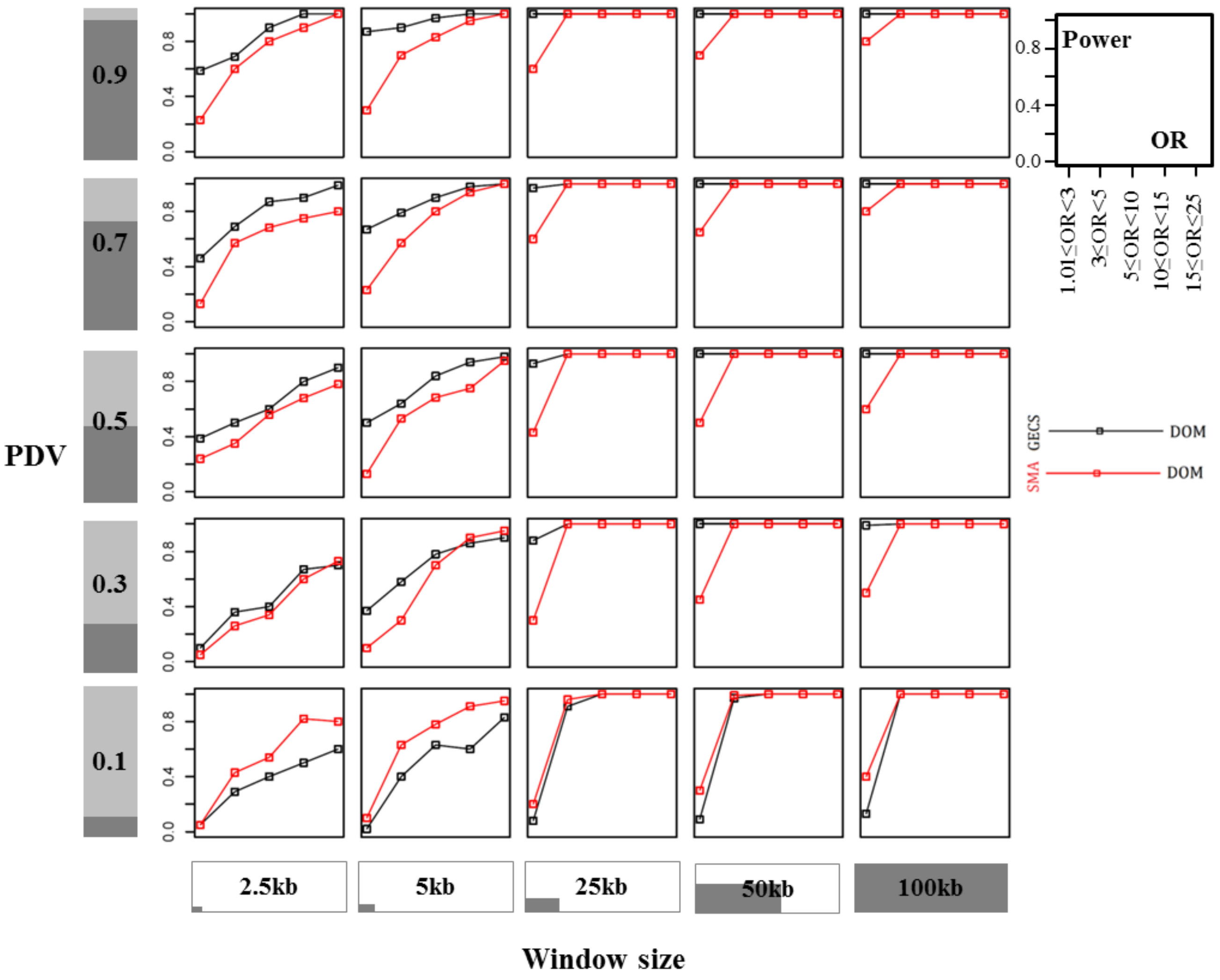

**Figure S6: Comparative power analysis for a common disease (prevalence K=0.1) and small sample size (N=1,000).** Results are given for studies with proportion of neutral rare variants (PNV) = 0.3, different simulated window sizes (x-axis), and different proportions of detrimental rare variants (PDV) (y-axis). Black lines: GECS; red lines: SMA. In each grid cell, the power is presented on the y-axis and OR intervals on the x-axis. For an overview see table S20.

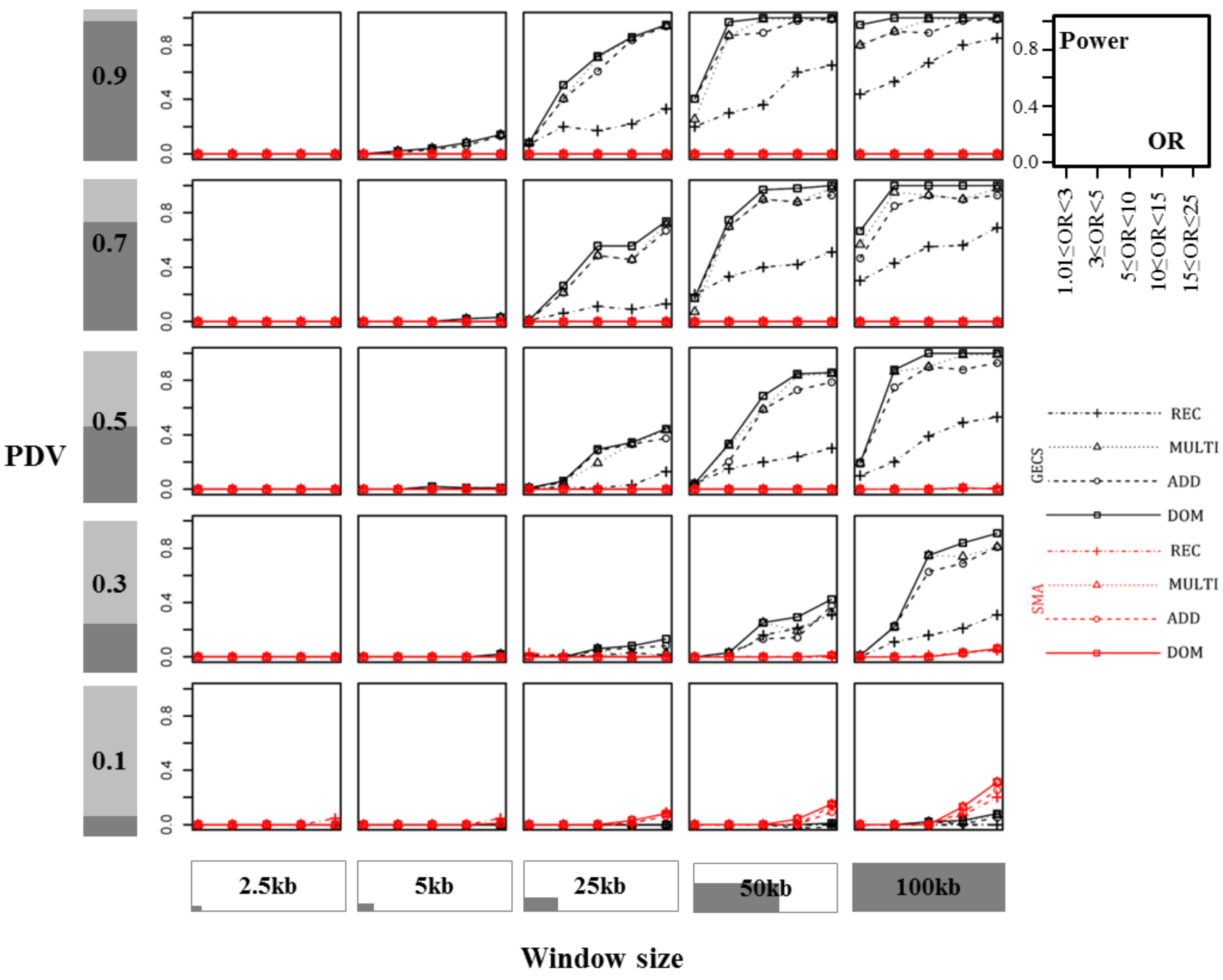

**Figure S7: Comparative power analysis for a common disease (prevalence K=0.1) and moderate sample size (N=10,000).** Results are given for studies with proportion of neutral rare variants (PNV) = 0.3, different simulated window sizes (x-axis), and different proportions of detrimental rare variants (PDV) (y-axis). Black lines: GECS; red lines: SMA. In each grid cell, the power is presented on the y-axis and OR intervals on the x-axis. For an overview see table S20.

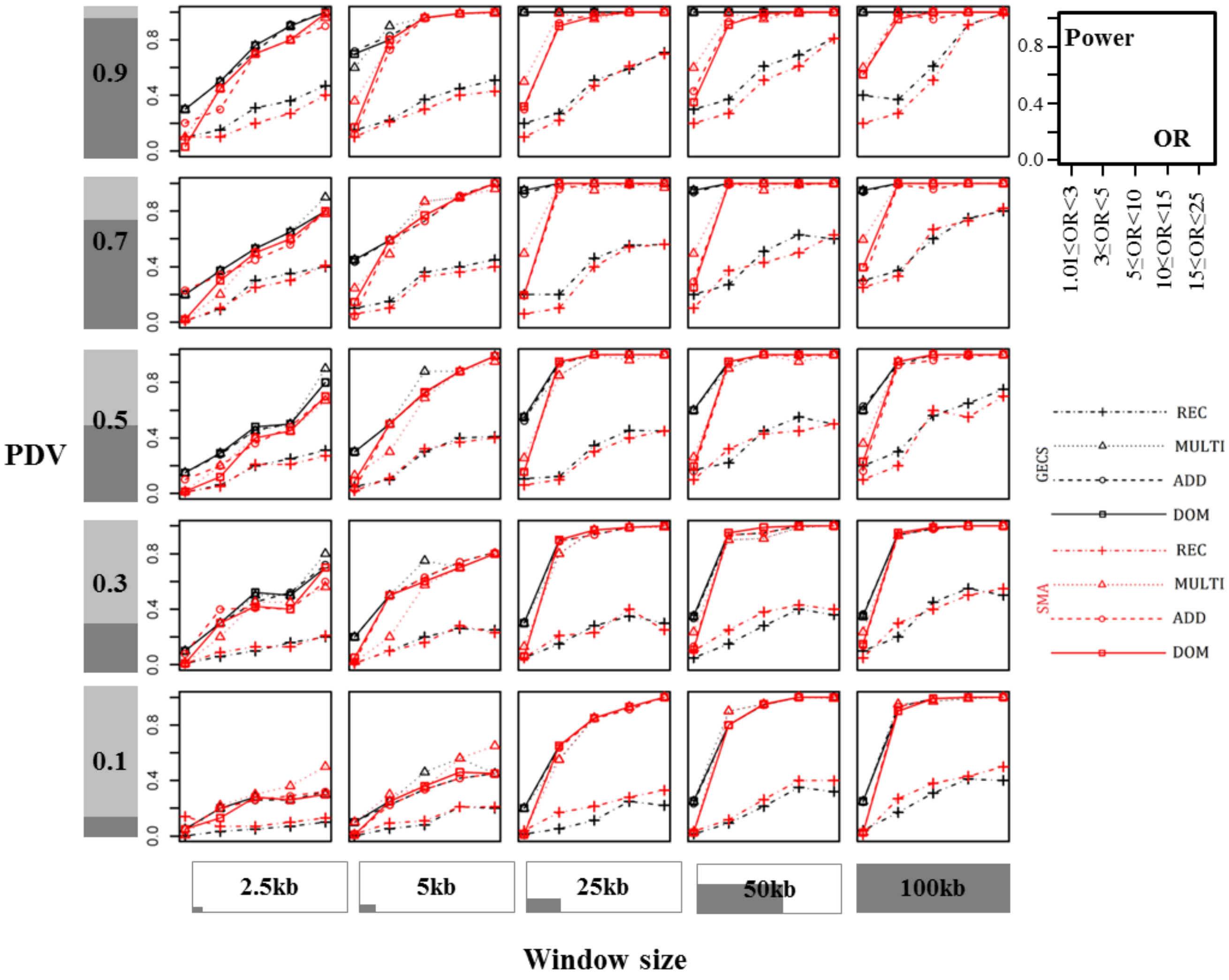

**Figure S8: Comparative power analysis for a common disease (prevalence K=0.1) and large sample size (N=20,000).** Results are given for studies with proportion of neutral rare variants (PNV) = 0.3, different simulated window sizes (x-axis), and different proportions of detrimental rare variants (PDV) (y-axis). Black lines: GECS; red lines: SMA. In each grid cell, the power is presented on the y-axis and OR intervals on the x-axis. For an overview see table S20.

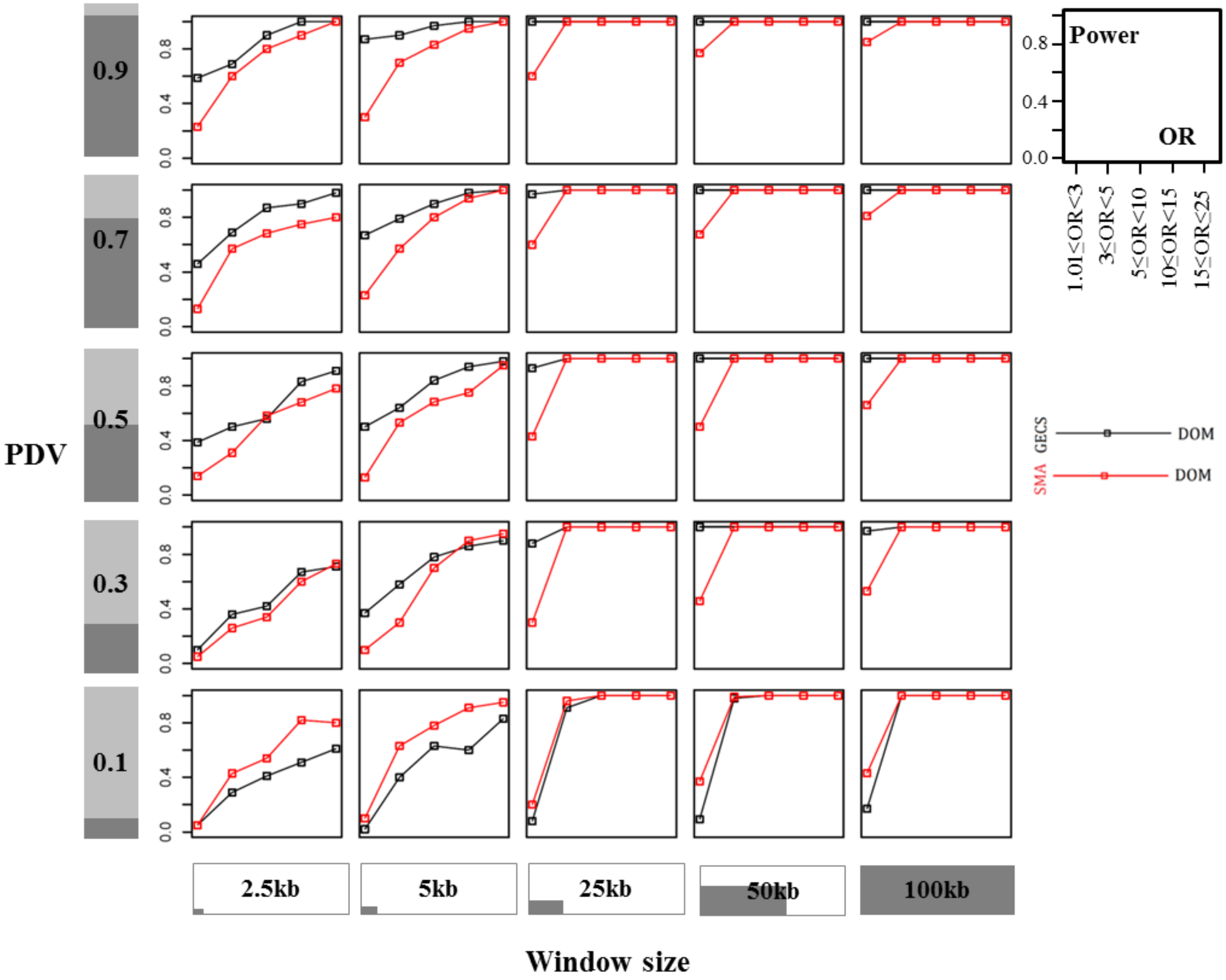

**Figure S9: Comparative power analysis for a rare disease (prevalence K=0.01) and small sample size (N=1,000).** Results are given for studies with proportion of neutral rare variants (PNV) = 0.1, different simulated window sizes (x-axis), and different proportions of detrimental rare variants (PDV) (y-axis). Black lines: GECS; red lines: SMA. In each grid cell, the power is presented on the y-axis and OR intervals on the x-axis. For an overview see table S20.

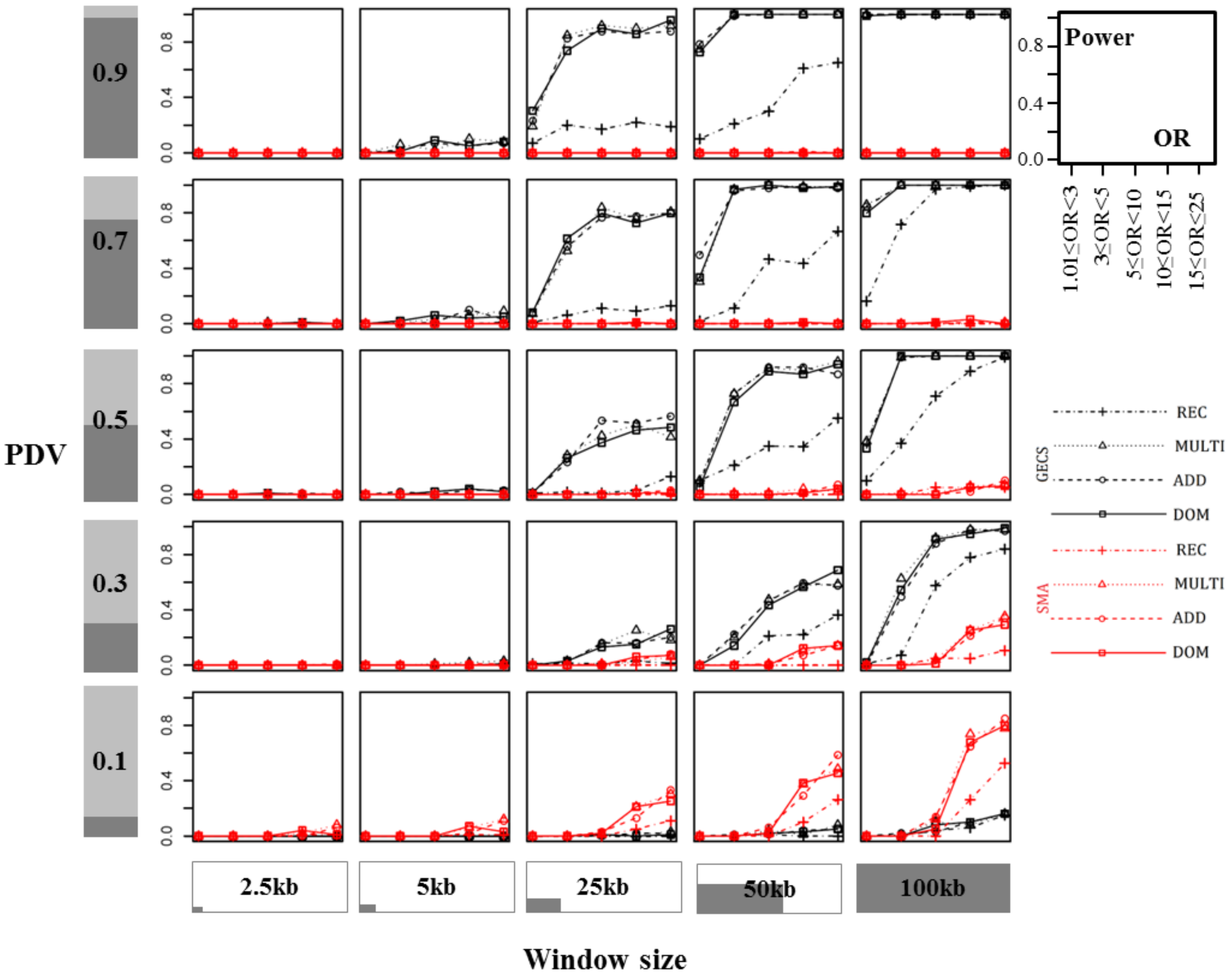

**Figure S10: Comparative power analysis for a rare disease (prevalence K=0.01) and moderate sample size (N=10,000).** Results are given for studies with proportion of neutral rare variants (PNV) = 0.1, different simulated window sizes (x-axis), and different proportions of detrimental rare variants (PDV) (y-axis). Black lines: GECS; red lines: SMA. In each grid cell, the power is presented on the y-axis and OR intervals on the x-axis. For an overview see table S20.

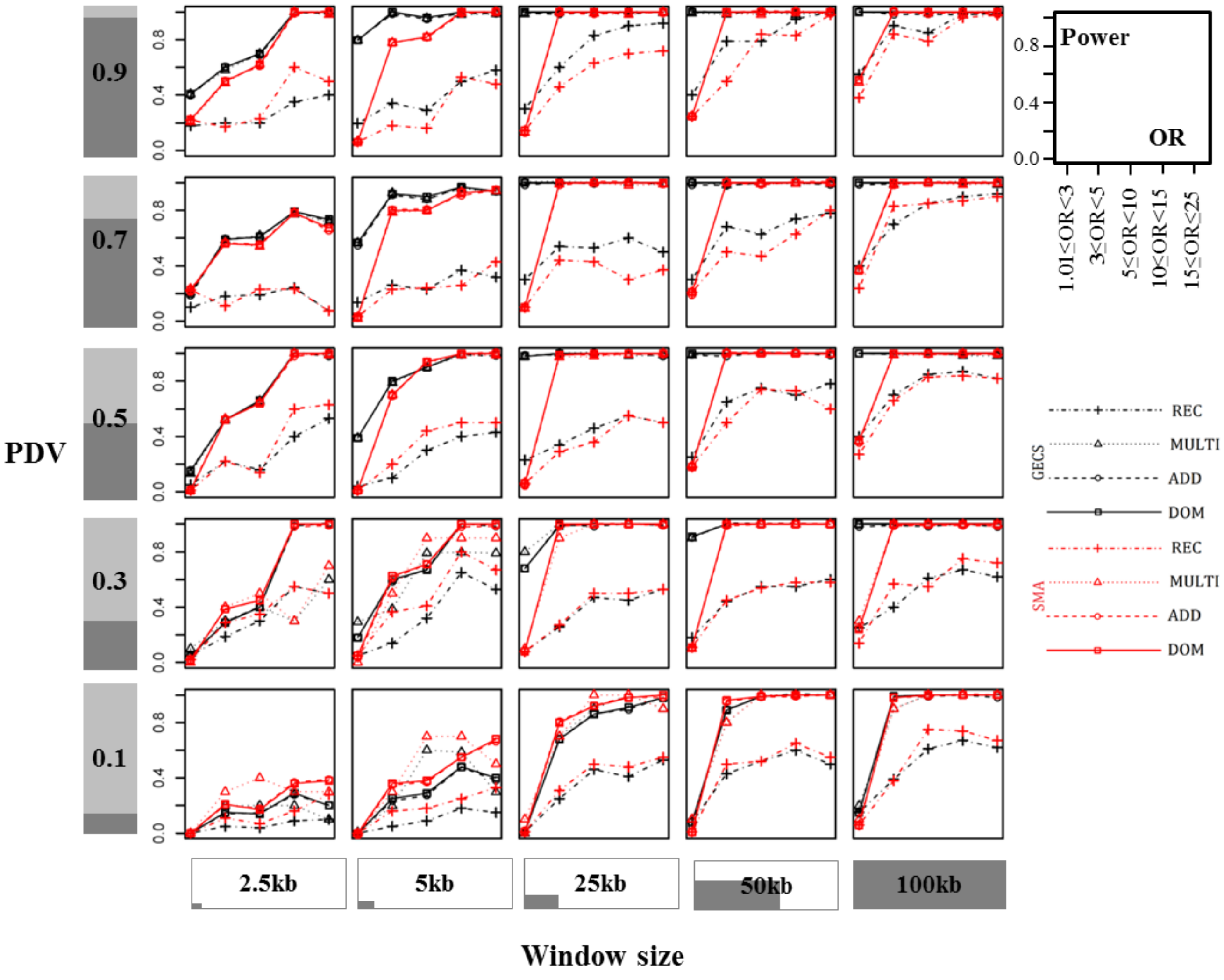

**Figure S11: Comparative power analysis for a common disease (prevalence K=0.1) and small sample size (N=1,000).** Results are given for studies with proportion of neutral rare variants (PNV) = 0.1, different simulated window sizes (x-axis), and different proportions of detrimental rare variants (PDV) (y-axis). Black lines: GECS; red lines: SMA. In each grid cell, the power is presented on the y-axis and OR intervals on the x-axis. For an overview see table S20.

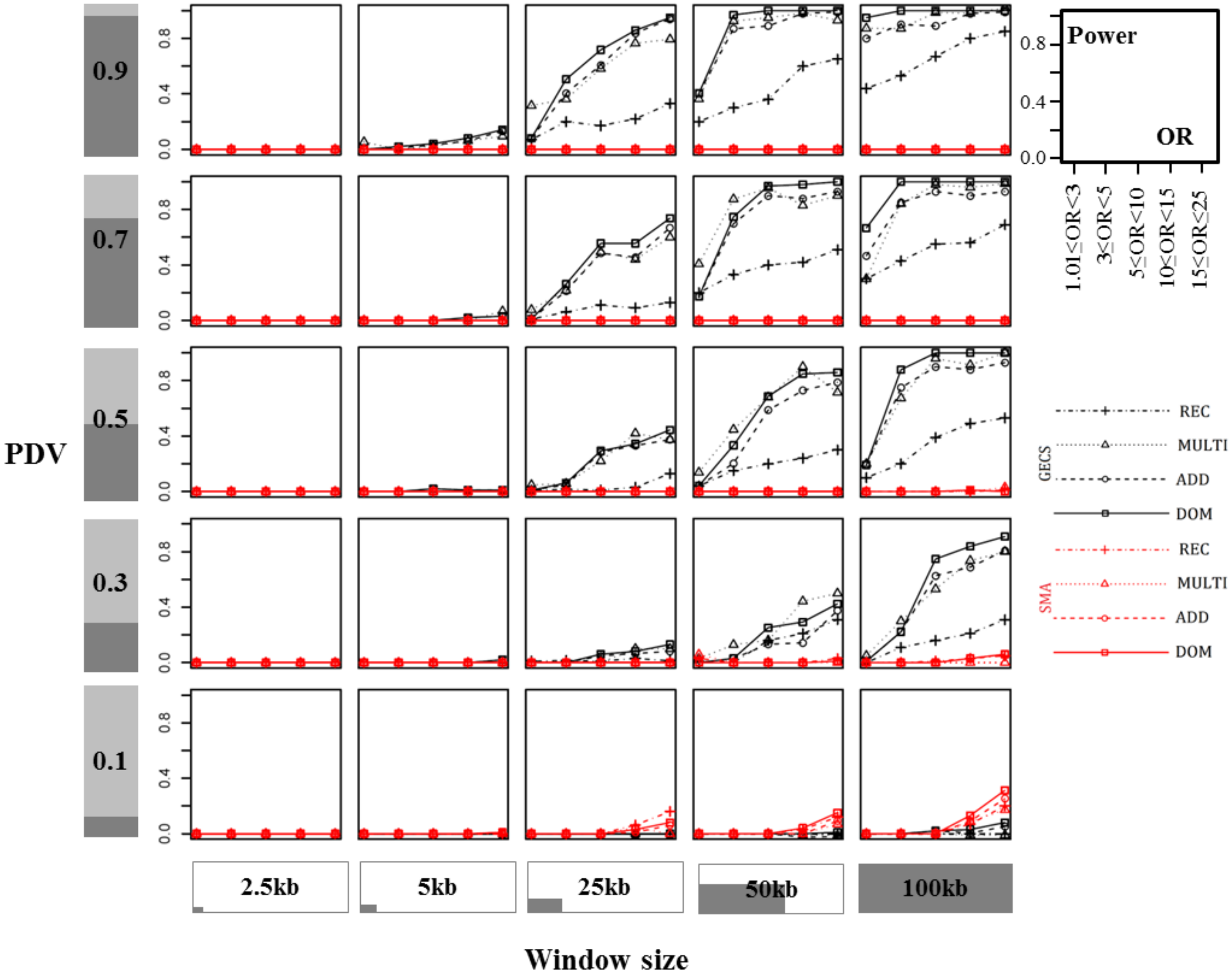

**Figure S12: Comparative power analysis for a common disease (prevalence K=0.1) and moderate sample size (N=10,000).** Results are given for studies with proportion of neutral rare variants (PNV) = 0.1, different simulated window sizes (x-axis), and different proportions of detrimental rare variants (PDV) (y-axis). Black lines: GECS; red lines: SMA. In each grid cell, the power is presented on the y-axis and OR intervals on the x-axis. For an overview see table S20.

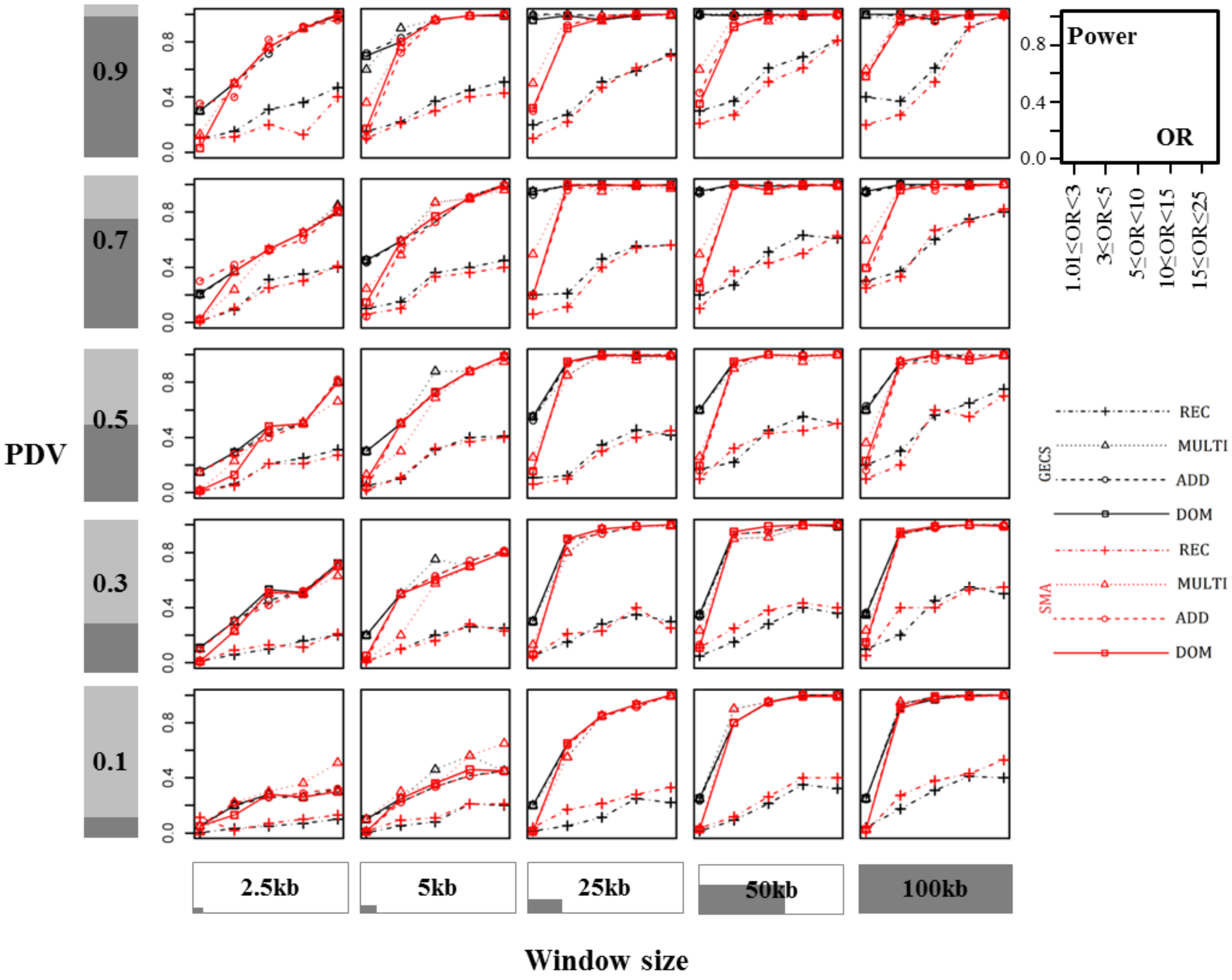

**Figure S13: Manhattan plot of single-marker analysis (SMA) for the AAMD data set. The y-axis is truncated at $-\log_{10}(p)=25$. The significance threshold (Table 2) and p-values were computed with the GECS software.**

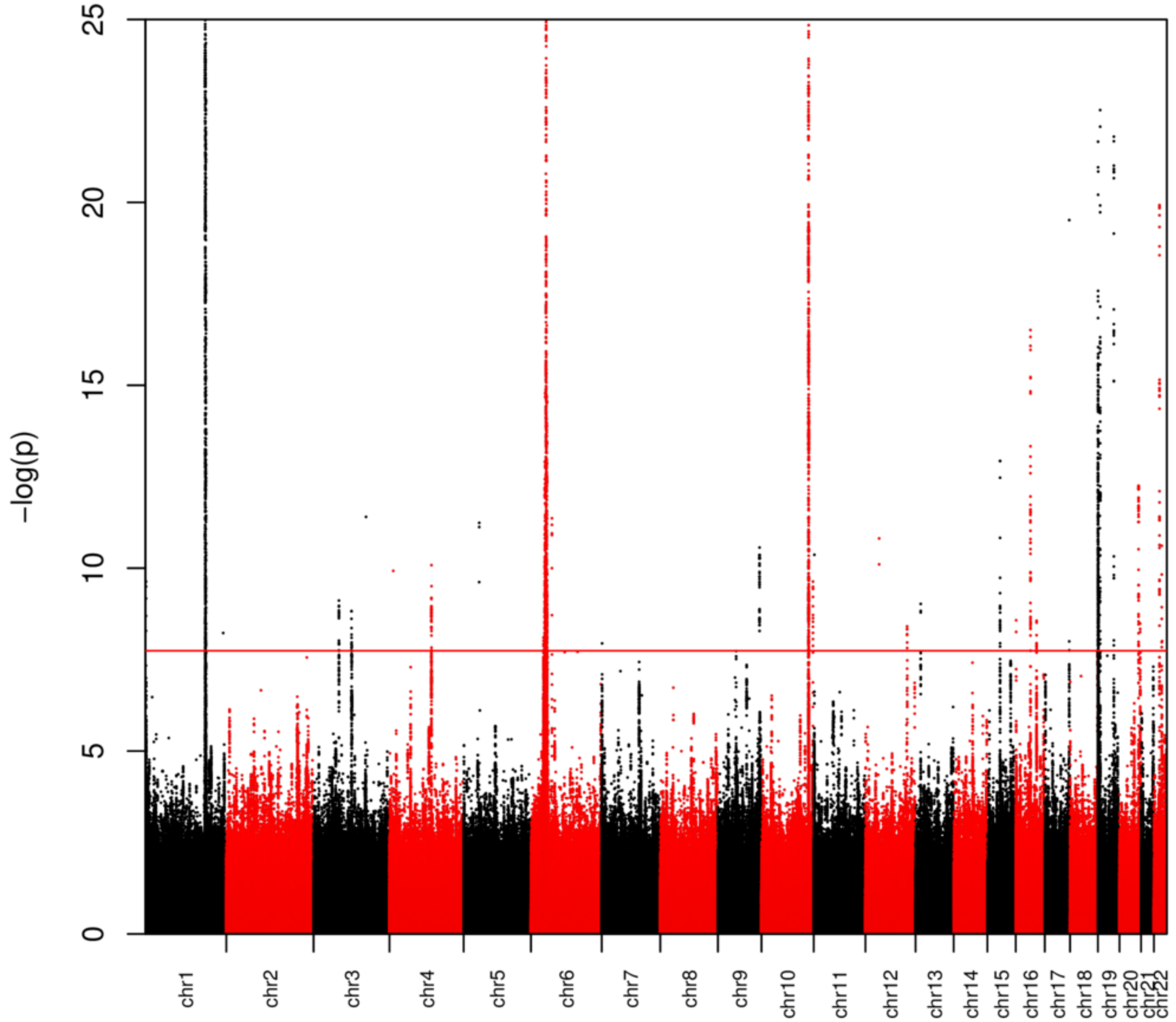

**Figure 14: Manhattan plot of bins at $MAF_T$=0.01 for the AAMD data set. Dots represent the middle position of bins (average between the start and end positions). The y-axis is truncated at $-\log_{10}(p)=25$ and values of $-\log_{10}(p)\leq 3$ are omitted. The significance threshold for the combined the three MAF thresholds (Table 2) and p-values were computed with the GECS software.**

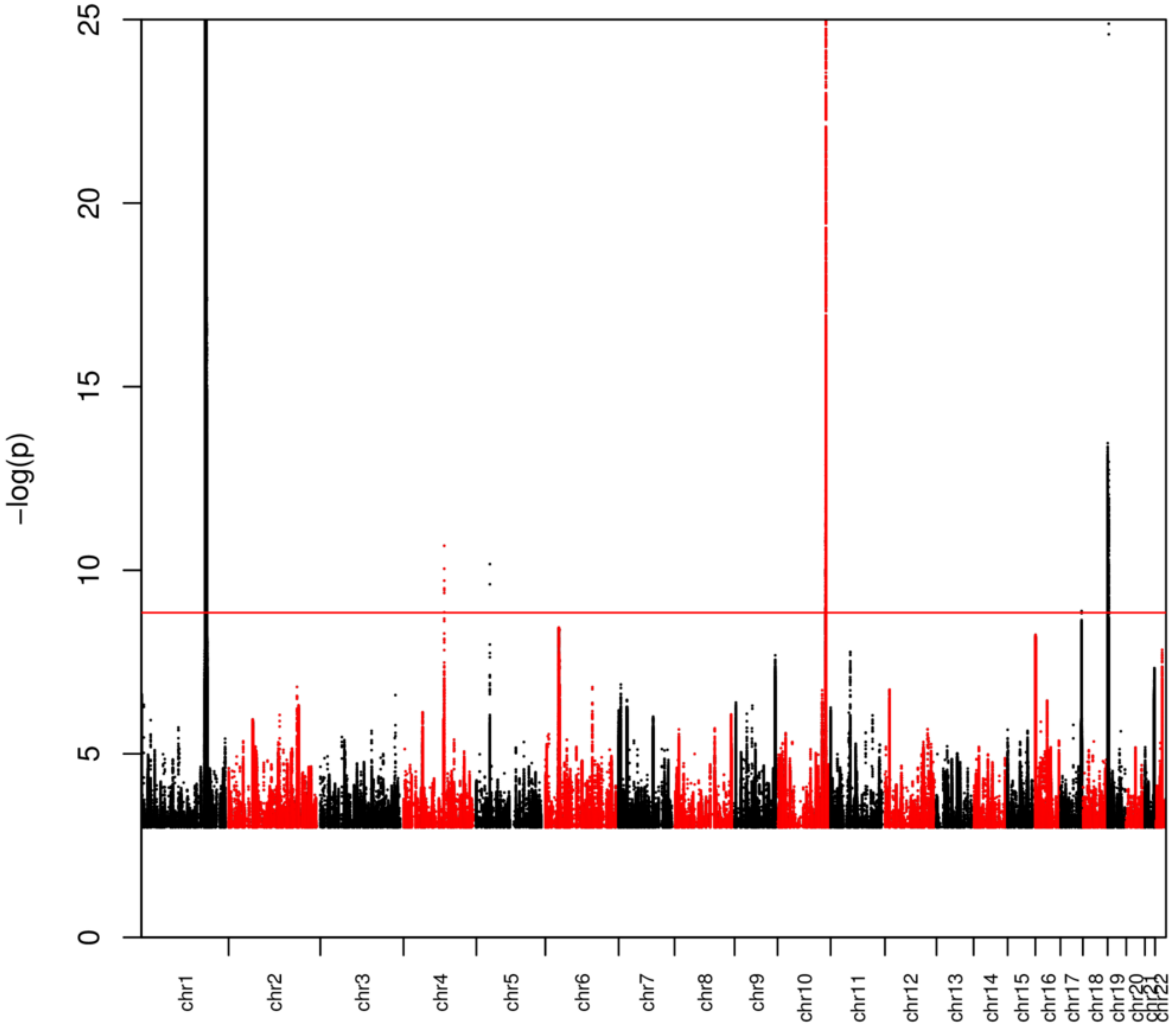

**Figure 15: Manhattan plot of bins at $MAF_T$=0.03 for the AAMD data set. Dots represent the middle position of bins (average between the start and end positions). The y-axis is truncated at $-\log_{10}(p)=25$ and values of $-\log_{10}(p)\leq 3$ are omitted. The significance threshold for the combined the three MAF thresholds (Table 2) and p-values were computed with the GECS software.**

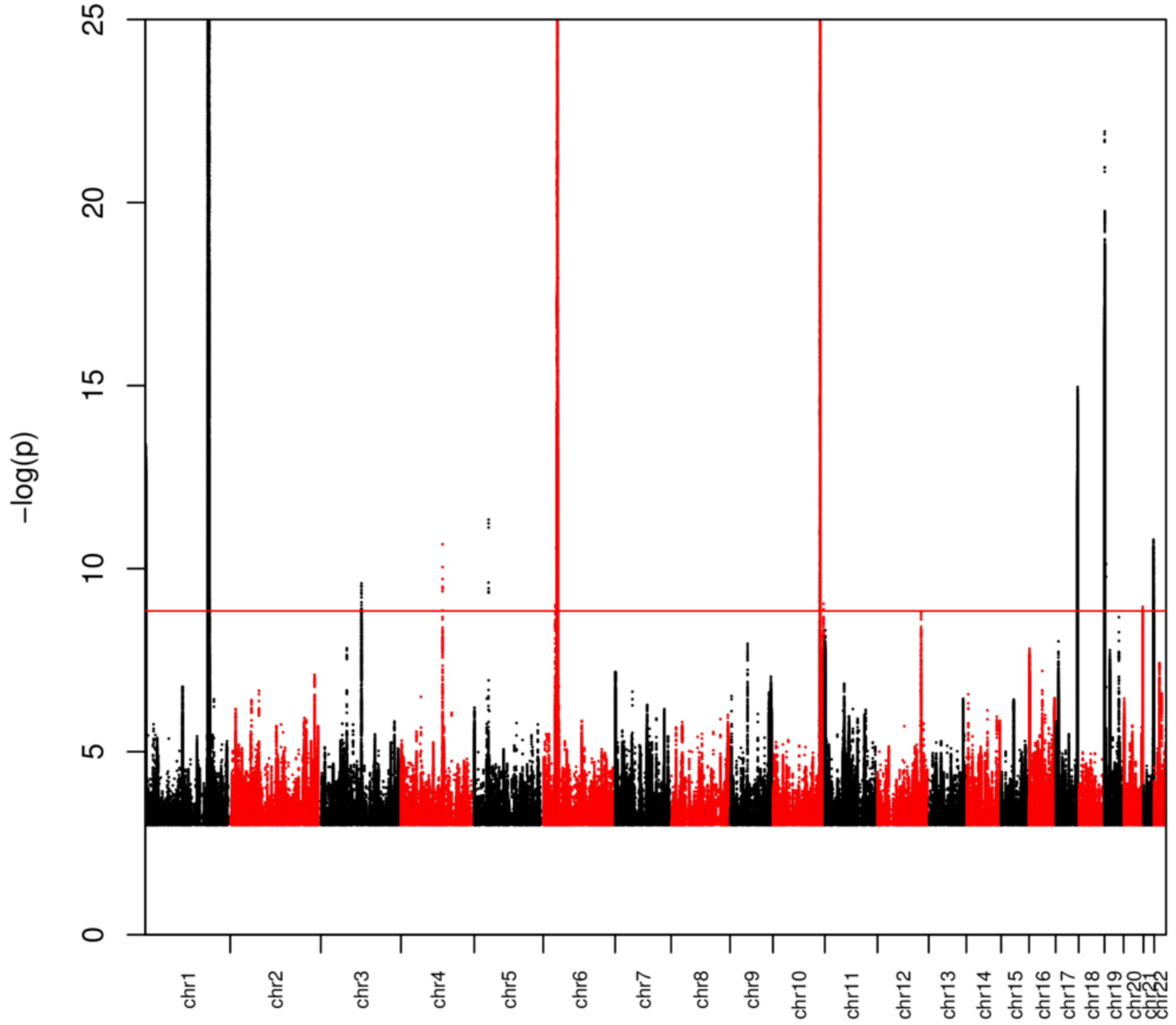

**Figure S16: Manhattan plot of bins at $MAF_T$=0.05 for the AAMD data set. Dots represent the middle position of bins (average between the start and end positions). The y-axis is truncated at $-\log_{10}(p)=25$ and values of $-\log_{10}(p)\leq 3$ are omitted. The significance threshold for the combined the three MAF thresholds (Table 2) and p-values were computed with the GECS software.**

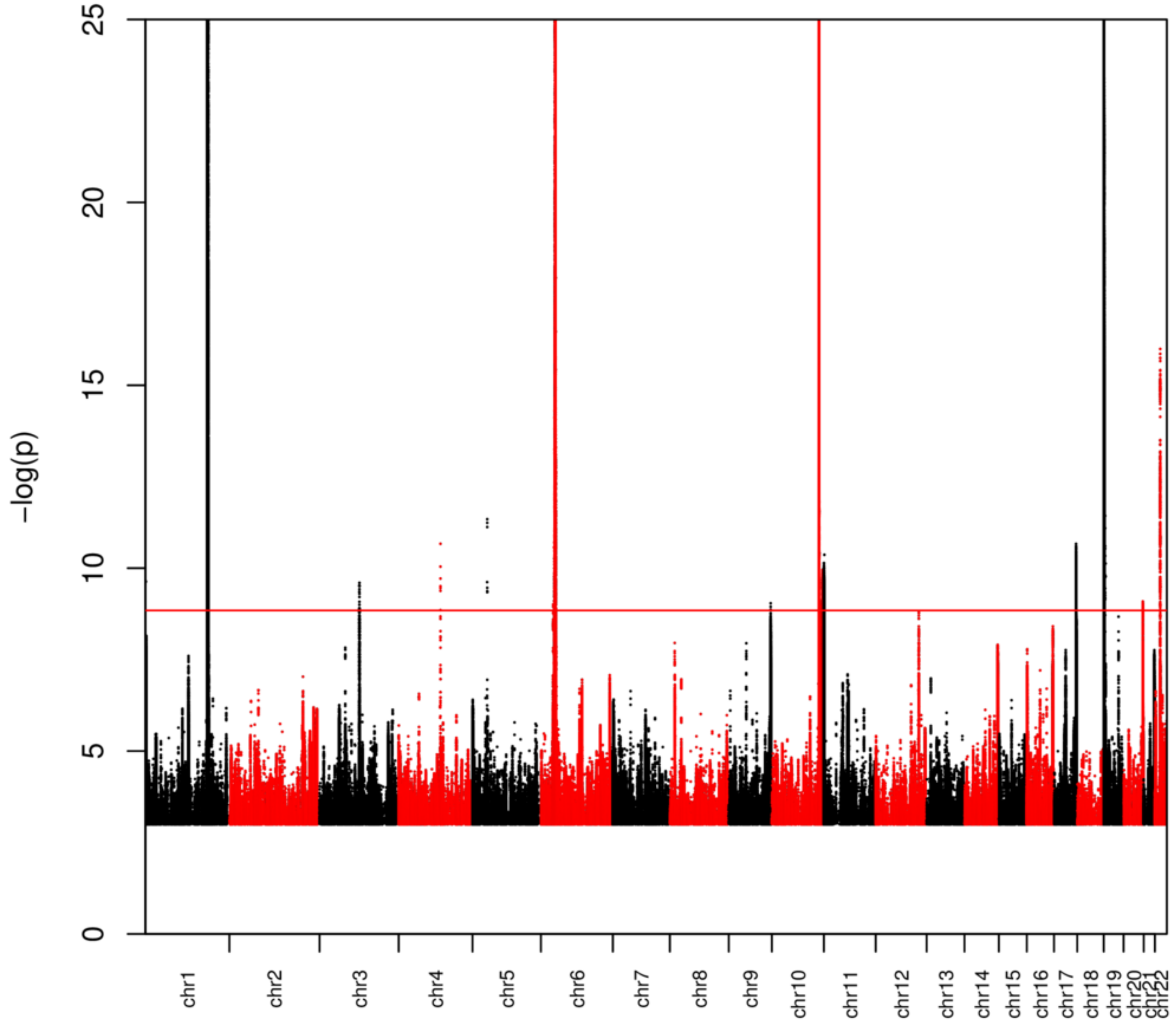

**Figure S17:**

The extended region [31323455-31323745] on chromosome 6 for NCT=2657 in the AAMD data set. Numbers in cells represent the $-\log_{10}$ of p*-values. Cells with the lowest p*-values are shown in red.

**Figure S18:**

The extended region [31473707-31474883] on chromosome 6 for NCT=2657 in the AAMD data set. Numbers in cells represent the $-\log_{10}$ of p*-values. Cells with the lowest p*-values are shown in red.

**Figure S19:**

The extended region [31878006-31878721] on chromosome 6 for NCT=2657 in the AAMD data set. Numbers in cells represent the $-\log_{10}$ of p*-values. Cells with the lowest p*-values are shown in red.

**Figure S20:**

The extended region [110685721-110685820] on chromosome 4 for NCT=1,611 in the AAMD data set. Numbers in cells represent the $-\log_{10}$ of p*-values. Cells with the lowest p*-values are shown in red.

Figure S17

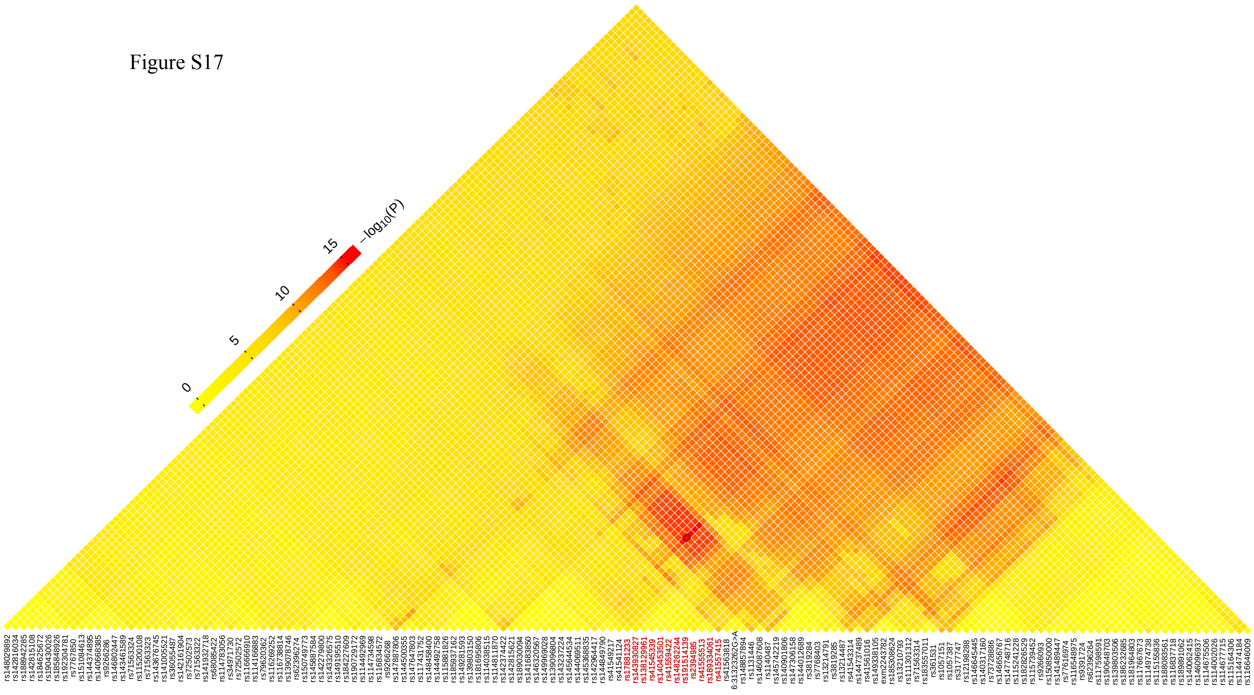

Figure S18

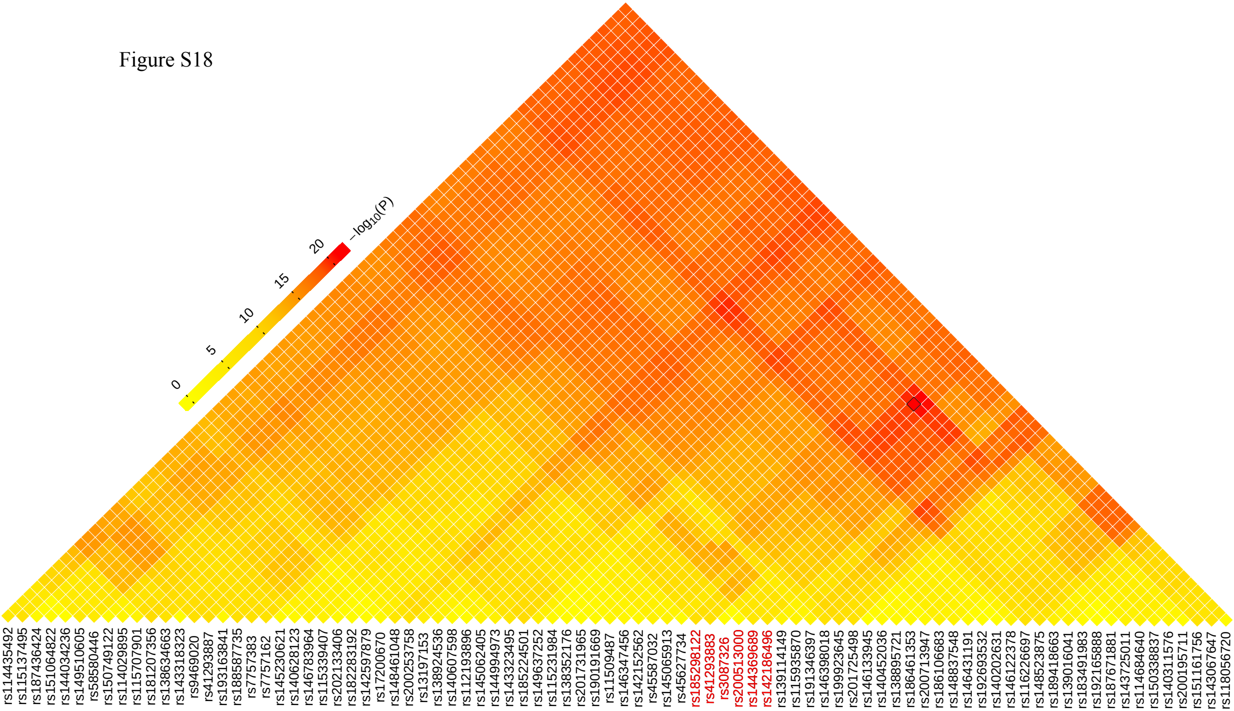
0
5
10
15
20
$-\log_{10}(P)$
rs114435492
rs115137495
rs187436424
rs151064822
rs144034236
rs149510605
rs58580446
rs150749122
rs114029895
rs115707901
rs181207356
rs138634663
rs143318323
rs9469020
rs41293887
rs193163841
rs188587735
rs7757383
rs7757162
rs145230621
rs140628123
rs146783964
rs115339407
rs202133406
rs182283192
rs142597879
rs17200670
rs148461048
rs200253758
rs13197153
rs138924536
rs140607598
rs112193896
rs145062405
rs144994973
rs143323495
rs185224501
rs149637252
rs115231984
rs138352176
rs201731965
rs190191669
rs115509487
rs146347456
rs142152562
rs45587032
rs145065913
rs45627734
rs185298122
rs41293883
rs3087326
rs200513000
rs144369689
rs142186496
rs139114149
rs115935870
rs191346397
rs146398018
rs199923645
rs201725498
rs146133945
rs140452036
rs138895721
rs186461353
rs200713947
rs186106683
rs148837548
rs146431191
rs192693532
rs140202631
rs146122378
rs116226697
rs148523875
rs189418663
rs139016041
rs183491983
rs192165888
rs187671881
rs143725011
rs114684640
rs150338837
rs140311576
rs200195711
rs151161756
rs143067647
rs118056720

Figure S19

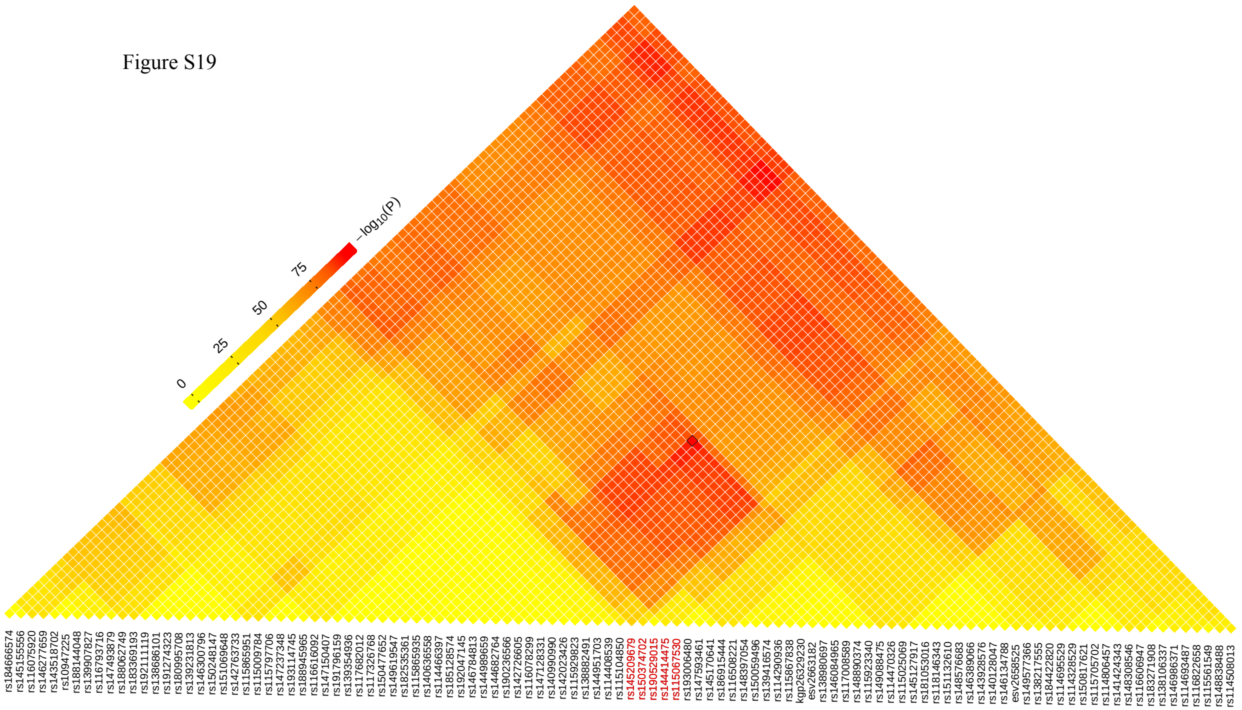
0
25
50
75
$-log_{10}(P)$
rs184666574
rs145155556
rs116075920
rs146277659
rs143518702
rs10947225
rs188144048
rs139907827
rs116793716
rs147493879
rs188062749
rs183369193
rs192111119
rs138686101
rs191274323
rs180995708
rs139231813
rs146300796
rs150248147
rs151069648
rs142763733
rs115865951
rs115009784
rs115797706
rs147237348
rs193114745
rs188945965
rs116616092
rs147150407
rs191796159
rs139354936
rs117682012
rs117326768
rs150477652
rs149619547
rs182535361
rs115865935
rs140636558
rs114466397
rs185128574
rs192047145
rs146784813
rs144989659
rs144682764
rs190236566
rs142726605
rs116078299
rs147128331
rs140990990
rs142023426
rs115929823
rs138882491
rs144951703
rs114408539
rs115104850
rs145209679
rs150374702
rs190529015
rs144414475
rs115067530
rs193006480
rs147593461
rs145170641
rs186915444
rs116508221
rs148397054
rs150059496
rs139416574
rs114290936
rs115867838
kgp2629230
esv2663182
rs138980697
rs146084965
rs117008589
rs148890374
rs115923340
rs149088475
rs114470326
rs115025069
rs145127917
rs181053016
rs118146343
rs151132610
rs148576683
rs146389066
rs143928576
rs140128047
rs146134788
esv2658525
rs149577366
rs138217555
rs184422820
rs114695529
rs114328529
rs150817621
rs115705702
rs114806457
rs141424343
rs148308546
rs116606947
rs183271908
rs138106337
rs146986371
rs114693487
rs116822658
rs115561549
rs148838488
rs114508013

Figure S20

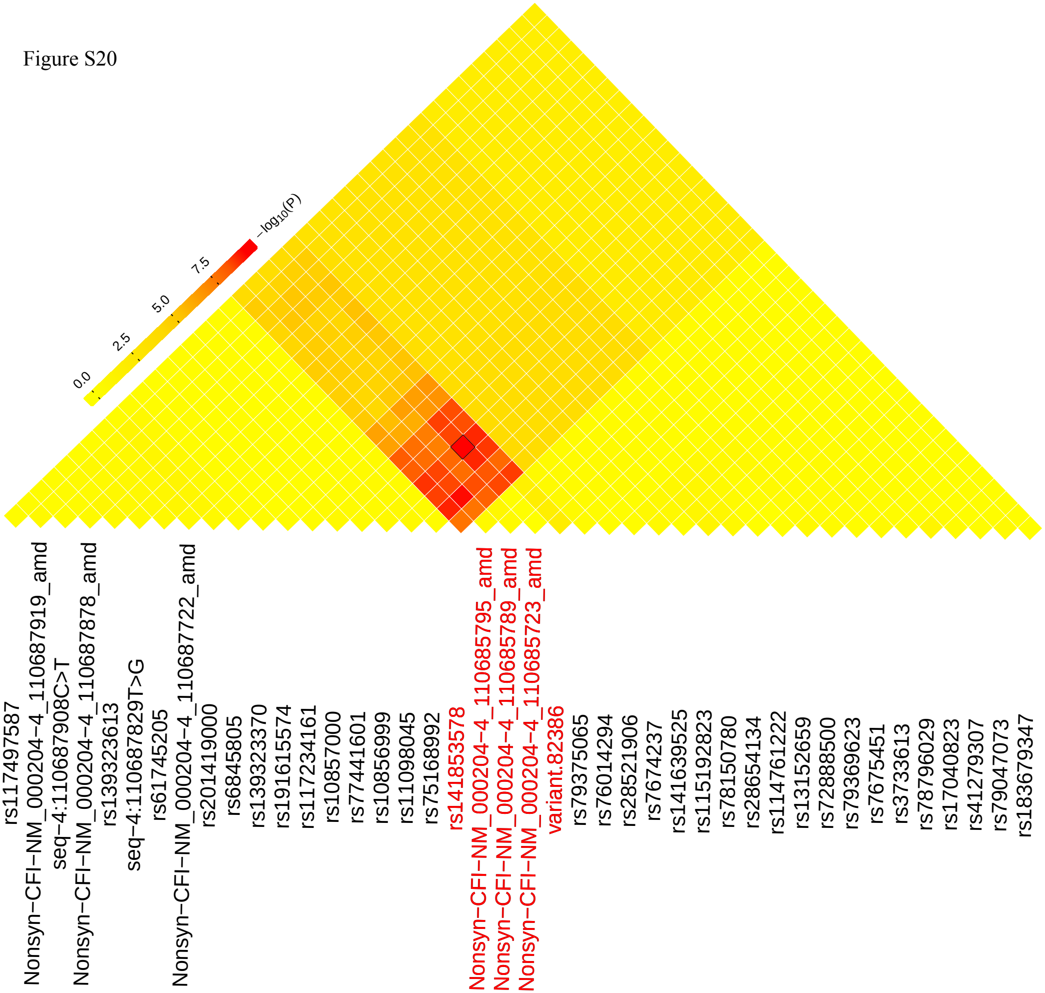
0.0
2.5
5.0
7.5
$-\log_{10}(P)$
rs117497587
Nonsyn−CFI−NM_000204−4_110687919_amd
seq−4:110687908C>T
Nonsyn−CFI−NM_000204−4_110687878_amd
rs139323613
seq−4:110687829T>G
rs61745205
Nonsyn−CFI−NM_000204−4_110687722_amd
rs201419000
rs6845805
rs139323370
rs191615574
rs117234161
rs10857000
rs77441601
rs10856999
rs11098045
rs75168992
rs141853578
Nonsyn−CFI−NM_000204−4_110685795_amd
Nonsyn−CFI−NM_000204−4_110685789_amd
Nonsyn−CFI−NM_000204−4_110685723_amd
variant.82386
rs79375065
rs76014294
rs28521906
rs7674237
rs141639525
rs115192823
rs78150780
rs28654134
rs114761222
rs13152659
rs72888500
rs79369623
rs7675451
rs3733613
rs78796029
rs17040823
rs41279307
rs79047073
rs183679347

**Figure 21: Manhattan plot of single-marker analysis (SMA) for the SCZD data set. The significance threshold (Table 2) and p-values were computed with the GECS software.**

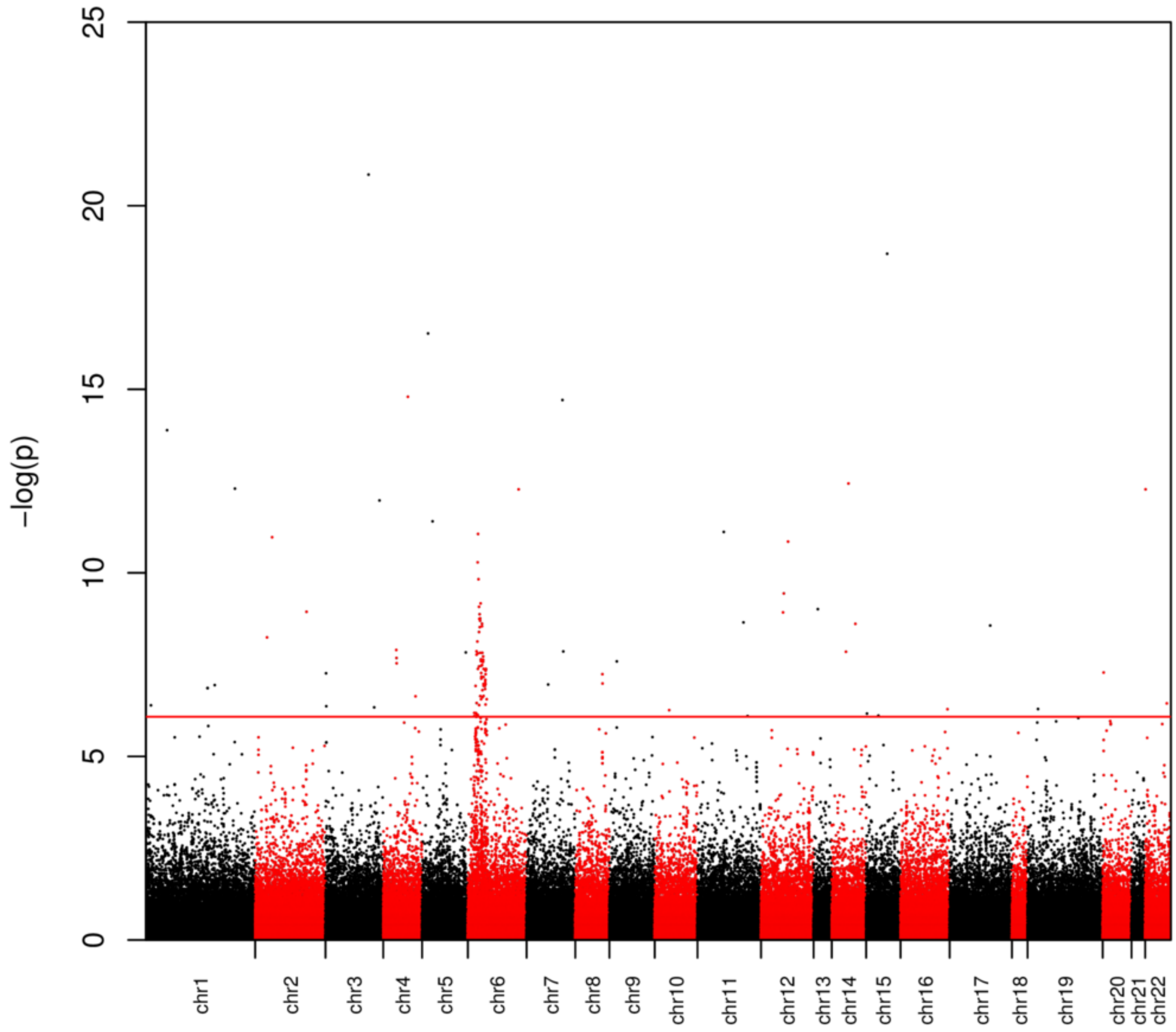

**Figure S22: Manhattan plot of bins at $MAF_T=0.01$ for the SCZD data set. Dots represent the middle position of bins (average between the start and end positions). Values of $-\log_{10}(p)\leq 2.5$ are omitted. The significance threshold for the combined the three MAF thresholds (Table 2) and p-values were computed with the GECS software.**

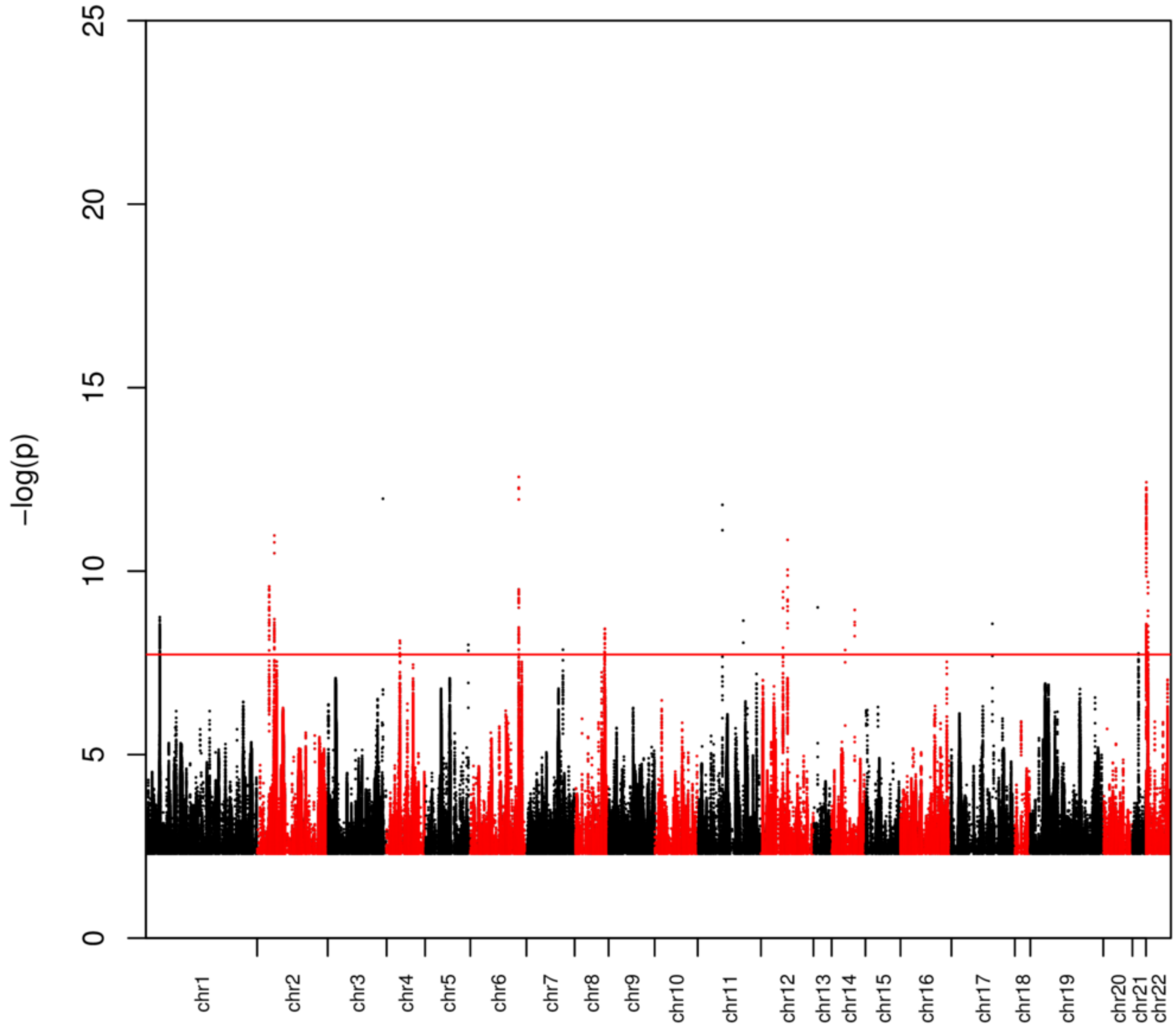

**Figure S23: Manhattan plot of bins at $MAF_T$=0.03 for the SCZD data set. Dots represent the middle position of bins (average between the start and end positions). Values of $-\log_{10}(p) \leq 2.5$ are omitted. The significance threshold for the combined the three MAF thresholds (Table 2) and p-values were computed with the GECS software.**

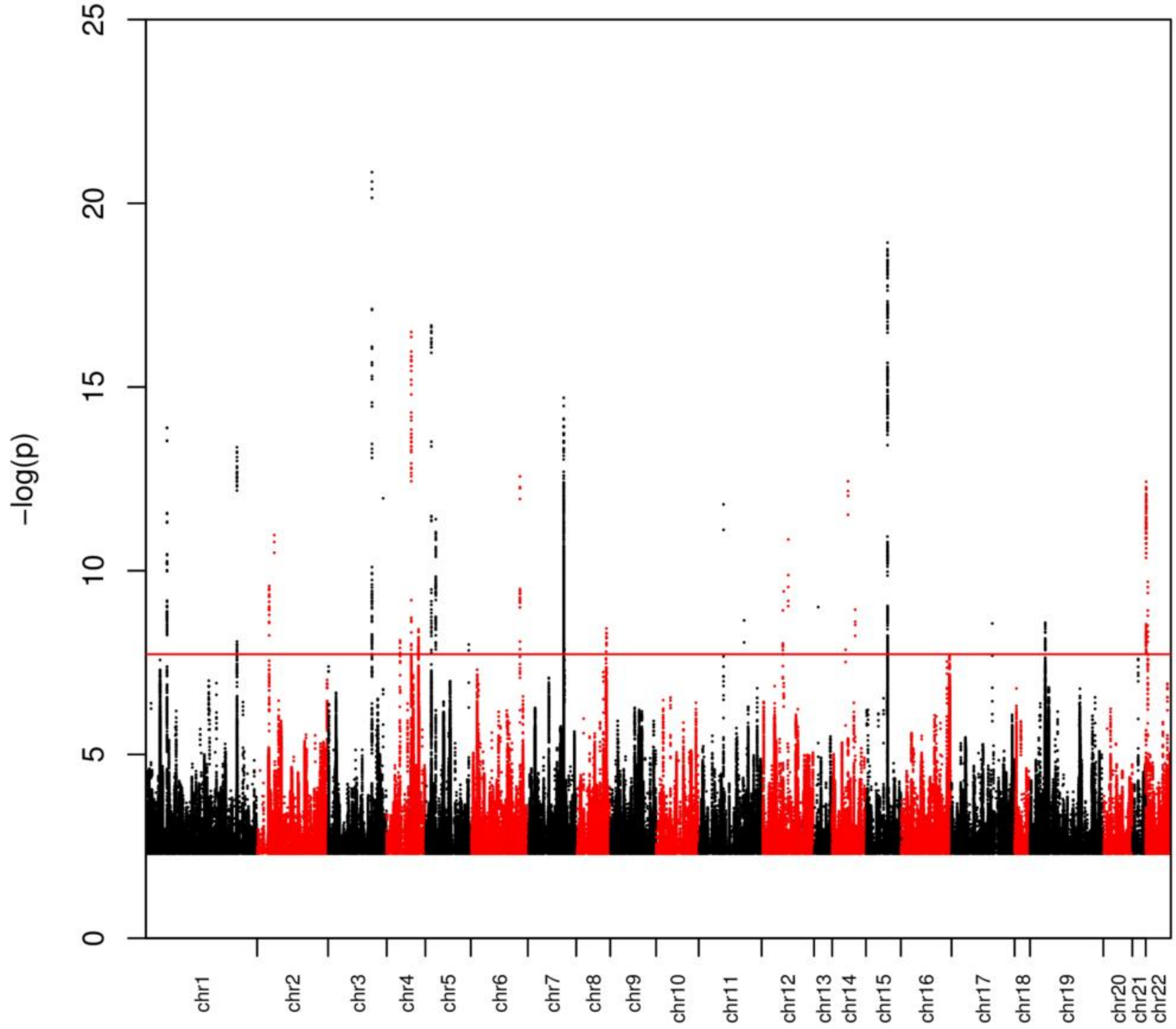

**Figure S24: Manhattan plot of bins at $MAF_T$=0.05 for the SCZD data set. Dots represent the middle position of bins (average between the start and end positions). Values of $-log_{10}(p) \leq 2.5$ are omitted. The significance threshold for the combined the three MAF thresholds (Table 2) and p-values were computed with the GECS software.**

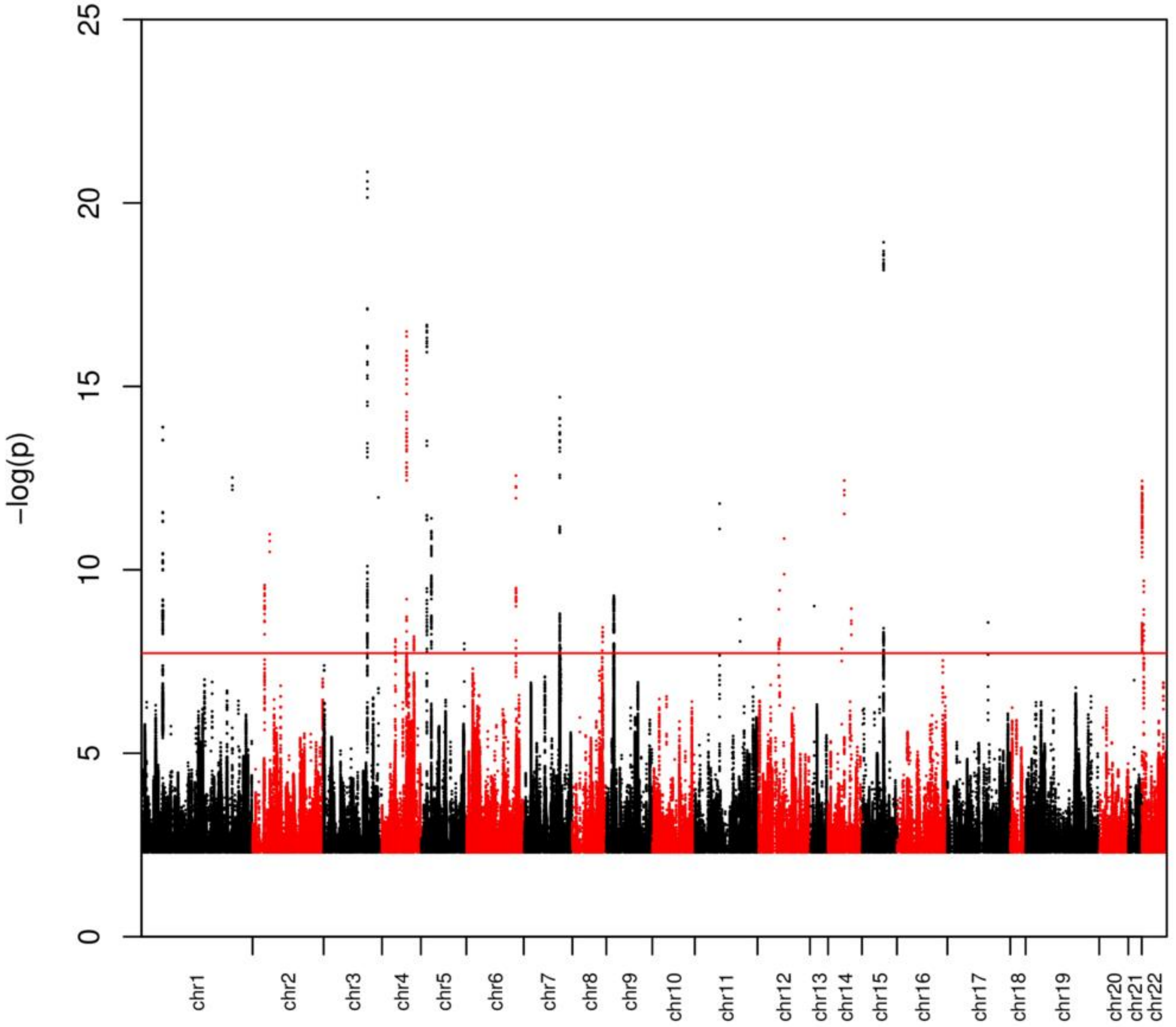

**Figure S25:**

The extended region [8999386-9028410] on chromosome 19 for NCT=636 in the SCZD data set. Numbers in cells represent the $-\log_{10}$ of p*-values. Cells with the lowest p*-values are shown in red.

Figure S25

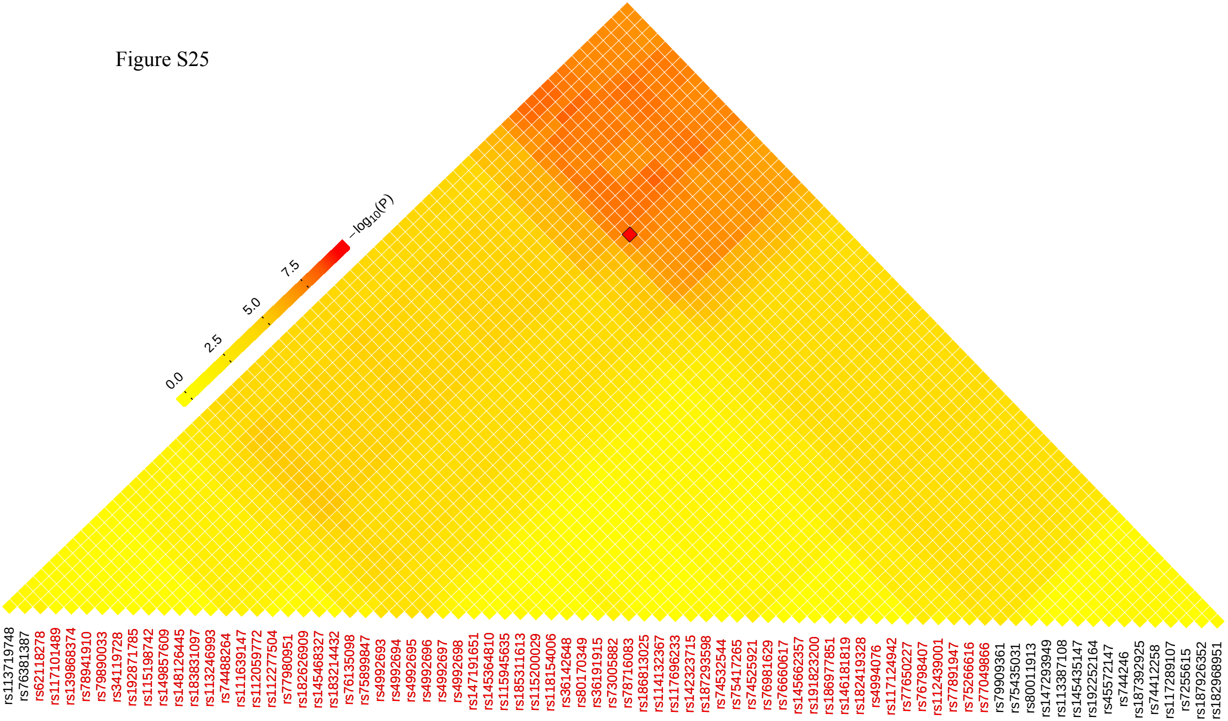

0.0
2.5
5.0
7.5
$-\log_{10}(P)$
rs113719748
rs76381387
rs62118278
rs117101489
rs139868374
rs78941910
rs79890033
rs34119728
rs192871785
rs115198742
rs149857609
rs148126445
rs183831097
rs113246993
rs74488264
rs111639147
rs112059772
rs112277504
rs77980951
rs182626909
rs145468327
rs183214432
rs76135098
rs75899847
rs4992693
rs4992694
rs4992695
rs4992696
rs4992697
rs4992698
rs147191651
rs145364810
rs115945635
rs185311613
rs115200029
rs118154006
rs36142648
rs80170349
rs36191915
rs73005882
rs78716083
rs186813025
rs114132367
rs117696233
rs142323715
rs187293598
rs74532544
rs75417265
rs74525921
rs76981629
rs76660617
rs145662357
rs191823200
rs186977851
rs146181819
rs182419328
rs4994076
rs117124942
rs77650227
rs76798407
rs112439001
rs77891947
rs75266616
rs77049866
rs79909361
rs75435031
rs80011913
rs147293949
rs113387108
rs145435147
rs192252164
rs45572147
rs744246
rs187392925
rs74412258
rs117289107
rs7255615
rs187926352
rs182968951

### Supplementary Notes:

#### a) Collapsing test (COLL)

Given a set of variants, the collapsing test COLL dichotomizes individuals by their carrier status, i.e. by presence of at least one minor allele. The resulting contingency table,

| #Individuals | Carrier | Non-carrier |
|---|---|---|
| Affected | $N_A^C$ | $N_A^N$ |
| Unaffected | $N_U^C$ | $N_U^N$ |

can be evaluated with Pearson $\chi^2$ -test with the test statistic

$$T_{COLL} = \frac{(N_A+N_U)(N_U N_A^C+N_A N_U^C)^2}{N_A N_U (N_A^C+N_U^C)(N_A^N+N_U^N)}.$$

The odds ratios (OR) are obtained from the table by computing

$$OR = \frac{N_A^C N_U^N}{N_U^C N_A^N}.$$

A rare-variant analysis procedure requires a definition of ”rareness”, which is usually provided by setting a minor allele frequency threshold ($MAF_T$) and considering all variants with minor allele frequencies below the threshold value as ”rare”.

#### b) GECS with Variable Binning (VB), an example:

In the following, we present a small example for which we specify all steps of the algorithm. Our goal is to find the locally distinct bins from the contiguous region that includes 8 variants in 4 individuals: $\left(v_1 = (1000), v_2 = (0100), v_3 = (0100), v_4 = (0010), v_5 = (0110), v_6 = (1000), v_7 = (1010), v_8 = (0001)\right)$

There are overall $8\,(8+1)/2 = 36$ combinations of start- end positions.

The full matrix $B$ is:

$$
B = \begin{bmatrix}
(1000) & (1100) & (1100) & (1110) & (1110) & (1110) & (1110) & (1111) \\
 & (0100) & (0100) & (0110) & (0110) & (1110) & (1110) & (1111) \\
 & & (0100) & (0110) & (0110) & (1110) & (1110) & (1111) \\
 & & & (0010) & (0110) & (1110) & (1110) & (1111) \\
 & & & & (0110) & (1110) & (1110) & (1111) \\
 & & & & & (1000) & (1010) & (1011) \\
 & & & & & & (1010) & (1011) \\
 & & & & & & & (0001)
\end{bmatrix}
$$

The algorithm executes as follows:

- Compute first bin to be analyzed: $B_{1,2} = v_1 \vee v_2 = (1100)$. This bin is locally distinct from both (1) $B_{1,1} = (1000)$and (2) $B_{2,2} = (0100)$.
- We keep track of the first identified distinct bin $B_{1,1}$by setting $I_1 = \{1\}$.
- Next bin to be analyzed is $B_{1,3} = B_{1,2} \vee v_3 = (1100)$. Variation of start and end positions identifies (1) $B_{1,3}$ as non-distinct from $B_{1,2}$ and (2) distinct from $B_{2,3} = (0100)$.
- Continuing with $B_{1,4}$, we find that $B_{1,4} = (1110) \neq B_{1,3} = (1100) \neq B_{2,3} = (0110) \Rightarrow B_{1,3}$ is a new distinct element: $I_1 = \{1,3\}$.
- $B_{1,5}$: (1) $B_{1,5} = B_{1,4} = (1110)$; (2) $B_{1,5} \neq B_{2,5}$, continue.
- $B_{1,6}$: (1) $B_{1,6} = B_{1,5} = (1110)$, but (2) $B_{1,6} = B_{2,6}$ implies that $B_{1,7} = B_{2,7}$ and $B_{1,8} = B_{2,8}$, so that $B_{1,6}, B_{1,7}, B_{1,8}$ are not distinct from bins that will be encountered in the family of bins starting at $v_2$, the algorithm terminates for the first row of $B$ and the final $I_1$ is $I_1 = \{1,3\}$, resulting in two distinct bins with start positions at $v_1$: $B_{1,1}$ and $B_{1,3}$.
-$B_{2,3}$: (1) and (2) are both satisfied: $B_{2,3} = B_{2,2} = B_{3,3}$, so that $I_2 = \{\}$ and the algorithm terminates for row 2.
-$B_{3,4}$: (1) and (2) are both not satisfied: $B_{3,4} = (0110) \neq B_{3,3} = (0100) \neq B_{4,4} = (0010) \Rightarrow$ $I_3 = \{3\}$.
-$B_{3,5}$: (1) and (2) satisfied: $B_{3,5} = B_{3,4} = B_{4,5} = (0110)$, terminate row.
-$B_{4,5}$: (1) $B_{4,5} = (0110) \neq B_{4,4} = (0010) \Rightarrow I_4 = \{4\}$, (2) $B_{4,5} = B_{55} = (0110)$, terminate row.
-$B_{5,6}$: (1) and (2): $B_{5,6} = (1110) \neq B_{5,5} = (0110) \neq B_{6,6} = (1000) \Rightarrow I_4 = \{5\}$.
-$B_{5,7}$: (1) $B_{5,7} = B_{5,6} = (1110)$, (2) $B_{5,7} \neq B_{6,7}$.

-$B_{5,8}$: (1) $B_{5,8} = (1111) \neq B_{5,7} = (1110) \Rightarrow I_5 = \{5{,}7\}$; $B_{5,8} = (1111)$ is a fully collapsed bin, terminate row.

-$B_{6,7}$: (1) $B_{6,7} = (1010) \neq B_{6,6} = (1000) \Rightarrow I_6 = \{6\}$, (2) $B_{6,7} = B_{7,7}$, terminate row.

-$B_{7,8}$: (1) and (2):$B_{7,8}(1011) \neq B_{7,7} = (1010) \neq B_{8,8} = (1000) \Rightarrow I_7 = \{7{,}8\}$

- Finally, include $B_{8,8}$: $I_8 = \{8\}$.

The algorithm finds the index sets $I = \{\{1{,}4\}, \{\}, \{3\}, \{4\}, \{5{,}7\}, \{6\}, \{7{,}8\}, \{8\}\}$. This results in identification of all single-variant bins $B_{i,i} = v_i$, apart from $v_2$ (since $v_2 = v_3$) and three bins consisting of two variants each: $B_{1,3} = v_1 \vee v_3 = (1100), B_{5,7} = v_5 \vee v_7(1110), B_{7,8} = v_7 \vee v_8 = (1011)$. Only 10 locally distinct, non-fully-collapsed bins remain out of 36 possible bins.

We wish to re-emphasize at this point that the algorithm identifies *locally* distinct bins. It can be stated that all overlapping bins that are left after application of the algorithm are distinct. Non-overlapping bins, however, may have identical carrier status also after application of the algorithm. This is actually a desirable feature, since bins from chromosomal regions that are far apart can have identical carrier status. In such a case, it is important to know the locations of the identical bins, since they will point to different genes. In addition, the output of the analysis software shall report also the boundaries locally equivalent bins. While they can be neglected for computation, their exact location is of interest.

#### c) GECS with variable threshold (VT):

The GECS method used till now a fixed threshold $N_T^C$, so that the number of carriers (i.e. the count of '1's in each variant's array) for each variant is below or equal to $N_T^C$:

$$\forall i : N_i^C = N_i^C(v_i) \equiv \text{BitCount}(v_i) \leq N_T^C \qquad (1)$$

, where the function 'BitCount' literally counts the number of '1's in each variant's array. We call $N_k^C$ the "level" of variant *k*.

The idea of VT analysis for a fixed bin is to find the "optimal level" that minimizes the p-value among a set of the locally distinct bins, followed by correction for multiple testing.

For the GECS approach with VT, it turns out that it is possible to combine the VB and VT methods and to shift through all combinations of start position, end position and level in a computationally efficient manner by only considering one representative bin from each family of locally non-distinct bins. The objective of GECS is to find all locally distinct bins for all combinations of three parameters.

We define $B_{i,j}$ (” $B_{i,j}$ at level $l$”) by combining all variants $v_k$ *with* $i \le k \le j$ *and* $N_k^C \le l$ using the logical ”OR”.

$$B_{i,j}^{l} = \begin{cases} 0 & : \quad \left\{ N_k^C \,\middle|\, i \le k \le j \,;\, N_k^C \le l \right\} = \emptyset \\ \bigvee_k v_k \,, k \in \{ i \le k \le j \mid N_k^C \le l \} & : \quad otherwise \end{cases} \quad (2)$$

Local non-distinctiveness at neighboring levels occurs if variants at level l + 1 do not contribute carries $l$ that are not in $B_{i,j}$ already:

$$B_{i,j}^{l} = B_{i,j}^{l+1}. \ (3)$$

We traverse the parameters $i, j$ as in the case of VB analysis, with additional traversal of possible l-values for fixed $i, j$.

If local non-distinctiveness is encountered during traversal of l, conditions (A1), (A2) have further consequences.

$$B_{i,j}^{l} = B_{i,j-1}^{l} \text{ with } l \le l_j \quad \text{(B1)}$$

$$\Rightarrow \ B_{i,j}^{l+k} = B_{i,j-1}^{l} \ \forall k\colon 0 \le N_T^C - l$$

$\Rightarrow$ stop traversing $l$, continue with next $j$.

This is a consequence of the fact that variant $v_j$ only contributes to bins at levels $l_j$ and higher. If (B1) is satisfied, we can stop traversing l for the current j. Likewise, if a fully collapsed bin is encountered; all bins at higher levels and larger j are guaranteed to be fully collapsed:

$$B_{i,j}^{l} = B_{i+1,j}^{l} \text{ or } B_{i,j}^{l} = 1 \quad \text{(B2)}$$

$$\Rightarrow \ B_{i,j+k_2}^{l+k_1} = B_{i+1,j+k_2}^{l+k_1} \ \forall k_1, k_2\colon 0 \le k_1 \le N_T^C - l; 0 \le k_2 \le n - j$$

$\Rightarrow$ stop traversing l; for fixed $i$, the highest level we need to consider is $l_{max} = l - 1$.

In other words, if $v_i$ does not contribute carriers to $B_{i,j}$, it will not contribute carriers to bins at either higher levels or bins with larger end coordinates.

Finally, local non-distinctiveness is possible at different levels for fixed $i$ and $j$:

$$B_{i,j}^{l} = B_{i,j}^{l-1} \ \forall l\colon l_i < l < l_{max} \quad \text{(B3)}$$

$\Rightarrow$ eliminate $B_{i,j}^{l}$ .

This condition is trivially satisfied if $B^{l}_{i,j-1} = B^{l-1}_{i,j-1}$.

As opposed to $i$ and $j$, most values of l do not correspond to potentially new distinct bins. For a fixed start position $i$, the lower bound of l is given by $l_i$. The highest possible non-trivial level for bins with coordinates $(i, j)$ is given by $l_{max} = \max(l_i, ..., l_j)$, although it can be lower due to (B2). Furthermore, for a fixed $j > i$, potentially new locally non-distinct bins can be found at levels $l \geq \max(l_i, l_j)$ were either $\hat{B}^{l}_{i,j-1} \neq \hat{B}^{l+1}_{i,j-1}$, or at level $l_j$ if $l_j > l_i$.

#### d) Combined results and correction for multiple testing

Let us consider a study with large sample size $N$. To make the conducting of GECS on such studies more feasible and less time consuming, maybe we need to conduct the analysis separately on the 22 chromosomes. Moreover, we want to conduct GECS with different $MAF_T$. As a result, we get for each of these sub-studies a separate list of maxT/minP of $n_0$ permutations. The combined result from all separate sub-studies must be also corrected for multiple testing. To hold the correction for multiple testing across the whole data, we have to conduct the maxT/minP approach for each permutation across all sub-studies, and subsequently the 5% level will be determined. In particular, for $m_0$ separate sub-studies and $n_0$ minP permutations conducted for each sub-study, the combined correction for multiple testing is

$$\alpha = 0.95\_quantile\ (P),$$

where

$$P = \left(P_1, P_2, \ldots, P_{n_0}\right),$$

and

$$P_i = \min\left\{P^i_1, P^i_2, \ldots, P^i_{m_0}\right\}.$$

Note that for sub-studies $s_1, s_2, \ldots, s_{m_0}$, the produced list of minP's of $n_0$ permutations for the *j-th* sub-study will be $(P^1_j, P^2_j, \ldots, P^{n_0}_j)$ and here the order of the permutation is very essential to get the aimed result.

In analogy, if we want to combine results from multiple $MAF_T$, we have first to conduct the maxT/minP approach for each permutation, and subsequently the 5% level will be determined.

#### The power in models of common diseases.

Applying GECS and SMA to diseases of higher prevalence (*K*=0.1) yielded some similarities to the results for rare diseases (*K*=0.01), but also some marked differences. In general, the power of both approaches slightly decreased. Differences between the two prevalence classes where most pronounced for small sample sizes (N=1,000) and a recessive inheritance mode.

Again, SMA performed better than GECS for PDV<0.3, but the power did not exceed 30% (SMA) and 10% (GECS), respectively, for the highest OR interval [15, 25]. Again, higher PDV values (PDV≥0.3) always yielded superior or at least equal power of GECS compared to SMA (Figures S6-7, S8). GECS attained about 100% power only for extreme PDV values (PDV=0.9) and extremely large OR values (≥15) and all but the recessive inheritance modes. Moderate sample size (N=10,000) strongly improved the power, ranging between 60-90% for studies with small to moderate OR values (1<OR≤5) and PDV ≥0.3 (Figure S7). Higher prevalence values also led to more nuanced results between the DOM, ADD and MULTI inheritance modes, although these remained marginal compared to the recessive mode. The moderate sample sizes provided substantial power (30-75%) even with small window sizes, although the power was consistently lower than for rare diseases (*K*=0.01) (Figures 3, S7). Increasing the sample size to N=20,000 resulted in an only negligible deviation from simulations with 10,000 individuals (Figures 3, S5, S7-8).

#### e) Extended results from AAMD data set

We applied GECS to the whole-genome imputed case-control data of the subset of samples with European ancestry and cases with AAMD from the international AMD genomics consortium (Table S2-3). The *CFI* gene was reported as associated with the AAMD phenotype and its subtypes in multiple studies (ALEXANDER *et al.* 2014; SEDDON *et al.* 2013; VAN DE VEN *et al.* 2013). We found a well-known rare variant in this region, rs141853578, on chromosome 4 (pos: 110,685,820bp), to be significantly associated ($p=3.1\times10^{-10}$). This SNP was covered by bin 4.I (chr4:110,685,721-110,685,820bp) that included five rare variants ($MAF_T \leq 0.01$) and was found to be significant with $p = 2.16\times10^{-11}$, $p' = 7.03\times10^{-10}$ and OR = 3.43 (Figure S20). The bin was more significant than the single rare SNP, but the SNP possessed slightly larger OR of 3.8. Additionally, in the local eSKAT analysis, we found the bin 4.II (chr4:110,685,721-110,685,962bp) with 6 variants, including all variants of the bin 4.I, with a more significant p'-value of $3.31\times10^{-10}$. In general, regions that showed a strong association of common variants (MAF>0.05) with the phenotype were covered by huge bins, being detected as significant at $MAF_T \leq 0.05$. This is because previously reported high-risk common variants may tag haplotypes that harbor rare variants included in these bins (some bins up to 5000 rare variants) (Table S13). As expected, local SKAT-based association testing with associated common variants as covariates led to disappearance of the association signal, confirming that the association seen in rare variants was only due to LD with nearby common variants (Table S13-14).

#### g) Benchmarking of GECS

We investigated the impact of the number of included variants and of the sample size on the computation time and the required amount of memory for GECS. For real and simulated studies, this algorithm results in performance increase of multiple orders of magnitude and allows for exhaustive computation and correction for multiple testing via permutations. Since the number of variants included in the analyses depended on the applied $MAF_T$ ($MAF_T$=0.5 for SMA; $MAF_T$=0.01, 0.03, 0.05, respectively, for GECS), it is useful to consider the number of performed tests of associations in each study. The study was conducted on Cheops, a high-performance cluster of university of Cologne, Germany (https://rrzk.uni-koeln.de/cheops.html). Nodes used for the calculations possess 24 GB RAM, 2.66 GHz each.

For small-sized studies (n=1,000), GECS performed on average 270 million (M) association tests, presenting the, number of the distinct bins in about 3 hours, requiring 3-4 GB (Table S18). In particular, GECS performed 276, 285, and 259 M association tests for 3.2, 5.3, and 6.2 M included variants, respectively. On the other hand, SMA performed 12.3 M association tests, representing the number of variants at $MAF_T$=0.5, in 10 minutes, requiring 7 GB. For 3.2 M variants, COLL-based GECS reduced the number of actually performed tests to 276 M distinct bins down from a possible ~300,000 M bins that result from the $O(n^2)$ complexity of a naïve exhaustive search, i.e. a reduction by more than three orders of magnitude. Interestingly, the reduction rate became higher with increasing $MAF_T$.

In general, the running time of both methods increased with increasing sample size, although to a lesser extent for SMA than for GECS. Moreover, the average reduction rate decreased slightly, which was more notable in for $MAF_T$=0.01. The GECS analysis of the imputed whole-genome data of AMD, with 27,259 samples and round 900,000 variants, took about 3-4 hours as we analyzed each chromosome in parallel. On average, GECS required fewer computational resources than SMA, with memory usage ranging between 1-5 GB for GECS and about 14 GB for SMA (Table S19). The analysis of the schizophrenia WES data (~10,000 samples, ~300,000 SNPs) took at maximum 14 h for GECS and 6 h for SMA (Table S19).
